# Histologic and Proteomic Remodeling of the Pulmonary Veins and Arteries in a Porcine Model of Chronic Pulmonary Venous Hypertension

**DOI:** 10.1101/2021.03.26.437051

**Authors:** Ahmed U. Fayyaz, Michael S. Sabbah, Surendra Dasari, Leigh G. Griffiths, Hilary M. DuBrock, M. Cristine Charlesworth, Barry A. Borlaug, Sarah M. Jenkins, William D. Edwards, Margaret M. Redfield

## Abstract

**AIM:** In heart failure (HF), pulmonary venous hypertension (PVH) produces pulmonary hypertension (PH) with remodeling of pulmonary veins (PV) and arteries (PA). In a porcine PVH model, we performed proteomic-based bioinformatics to investigate unique pathophysiologic mechanisms mediating PA and PV remodeling.

**METHODS:** Large PV were banded (PVH, n= 10) or not (Sham, n=9) in piglets. At sacrifice, PV and PA were perfusion labeled for vessel specific histology and proteomics. The PA and PV were separately sampled with laser-capture micro-dissection for mass spectrometry.

**RESULTS:** Pulmonary vascular resistance (Wood Units; 8.6 versus 2.0) and PA (19.9 versus 10.3) and PV (14.2 versus 7.6) wall thickness/external diameter (%) were increased in PVH (p<0.01 for all). Similar numbers of proteins were identified in PA (2093) and PV (2085) with 94% overlap, but biological processes differed. There were more differentially expressed proteins (287 versus 161), altered canonical pathways (17 versus 3) and predicted up-stream regulators (PUSR; 22 versus 6) in PV than PA. In PA and PV, bioinformatics indicated activation of the integrated stress response and mTOR signaling with dysregulated growth. In PV, there was also activation of Rho/Rho kinase signaling with decreased actin cytoskeletal signaling and altered tight and adherens junctions, ephrin B, and caveolar mediated endocytosis signaling; all indicating disrupted endothelial barrier function. Indeed, protein biomarkers and the top PUSR in PV (TGF-β) indicated endothelial mesenchymal transition (EndoMT) in PV. Findings were confirmed in human autopsy specimens.

**CONCLUSION:** These findings provide new therapeutic targets to oppose pulmonary vascular remodeling in HF-related PH.

**TRANSLATIONAL PERSPECTIVE:** In heart failure (HF) related (Group 2) PH, despite remodeling of pulmonary veins (PV) and arteries (PA), therapies targeting PA biology altered in Group 1 PH have not shown consistent benefit. In a porcine Group 2 PH model, microdissection allowed vessel specific (PV and PA) proteomics/bioinformatics. In PA and PV, the integrated stress response and mTOR signaling were activated with evidence of dysregulated growth. In PV, many more pathways were altered with broad evidence of disrupted endothelial barrier function and endothelial mesenchymal transition. Findings were confirmed in human specimens and provide new therapeutic targets in Group 2 PH.

## INTRODUCTION

Heart failure (HF) related pulmonary venous hypertension (PVH) is a common cause of pulmonary hypertension (Group 2 PH). Group 2 PH is a poor prognostic factor and a potential therapeutic target in HF.^1^ However, Group 1 PH therapies in HF have shown mixed but mostly non-beneficial effects.^2^ In human Group 2 PH, pulmonary veins (PV) and arteries (PA) are remodeled.^3, 4^ Several pathophysiologic pathways contribute to disrupted PA biology in non-Group 2 PH and representative animal models^5–13^ and may or may not mediate PVH induced pulmonary vascular remodeling. Further, as PV are developmentally, physiologically and structurally distinct from PA,^4, 14, 15^ we hypothesized that mechanisms involved in PV versus PA remodeling may differ, with therapeutic implications.

To investigate biologic responses to PVH in PV and PA, we used an established porcine model produced by banding of the large PV.^16–18^ Post-mortem PA versus PV perfusion labeling allowed vessel specific histomorphology and laser capture micro-dissection (LCMD) for proteomics (liquid chromatography tandem mass spectrometry (LC–MS/MS)). Differentially expressed proteins (DEP) versus Sham controls were analyzed using Ingenuity Pathways Analysis (IPA) software to identify key canonical pathways and predicted upstream regulators (PUSR) altered in PA and PV in experimental Group 2 PH. Whole lung (WL) samples were harvested for LC-MS/MS to provide context to LCMD findings and owing to larger sample and protein recovery, additional discovery potential. In addition to IPA analysis, select proteins relevant to current PH biology and therapeutics were analyzed individually. Endothelial mesenchymal transition (EndoMT) is an important pathophysiologic mechanism in PH of other etiologies^19–21^ but lacks well-established bioinoformatics protein/gene sets. Thus, we tabulated previously reported EndoMT protein biomarkers^19–24^ and analyzed their change in PVH vs Sham pigs. Lastly, we performed LCMD of PA and PV in autopsy specimens from patients with HF related PH (n=3) and Controls (n=3) to assess generalizability of our findings to the human.

## METHODS

This study was approved by the Mayo Clinic Institutional Review Board and the Institutional Animal Care and Use Committee (IACUC) and performed in accordance with the Guidelines for the Care and Use of Laboratory Animals. See Supplemental Materials for detailed methods regarding each of the study procedures outlined briefly below.

### Anesthesia, analgesia and euthanasia

For thoracotomy, all animals were given intramuscular Telazol (5.0 mg/kg) and Xylazine (1-2 mg/kg) prior to intubation. General anesthesia was maintained with 1.5-3.0% isoflurane. Additional analgesia was provided through fentanyl (intravenous bolus of 100 mcg followed by infusion of 2-5 mcg/kg/hr for the duration of the procedure). All animals received Buprenorphine SR immediately following thoracotomy closure. For monthly RHC, animals were administered Carprofen 4 mg/kg subcutaneously for perioperative analgesia. Anesthesia utilized Telazol, Xyalzin and isoflurane as for thoracotomy. Experimental animals underwent a final RHC and were euthanized with pentobarbital sodium at end study according to National Institute of Health (NIH) and Mayo IACUC guidelines.

### Study design (Supplemental Figure 1)

In male domestic piglets (n=23, weight approximately 10 kg), PVH was produced (n=14) by surgical banding of the left anterior PV and the posterior common PV.^16–18^ Sham animals (n=9) underwent thoracotomy without banding. A right heart catheterization (RHC) with pulmonary arteriogram and pulmonary venogram were performed at baseline and monthly intervals. In a randomly selected sub-group (n=7), CardioMEMS^®^ PA pressure monitors (Abbott Laboratories, Chicago IL) were implanted at the one-month RHC. Conscious measurements were recorded intermittently.

The PVH pigs underwent a final RHC and were euthanized when they met end-study (ES) criteria defined as clinical instability (n=4), severe combined pre-capillary and post-capillary hypertension (CpcPH) (n=5) or 16 weeks post-banding (n=1). Sham pigs were sacrificed at similar time points to PVH pigs.

### Hemodynamic evaluation

Right atrial (RA), right ventricular (RV), pulmonary artery (PA), and pulmonary capillary wedge (PCW) pressures were recorded. Cardiac output (CO) was measured by thermodilution. Stroke volume (SV), pulmonary artery capacitance (cPA), transpulmonary gradient (TPG), pulmonary vascular resistance (PVR) and diastolic pressure gradient (DPG) were calculated using standard formulae.

### Post-Mortem Pulmonary Vasculature Labeling and Lung Histology

As small PA and PV can be difficult to discriminate, vessel specific perfusion using microbead-agarose or barium-agarose solutions was used to label PA and PV.^4^ Lung sections were stained with hematoxylin and eosinophil (H&E) and Verhoeff-van Gieson (VVG) stains, later stain highlighting collagen/fibrosis in red, smooth muscle cells in brown and elastic fibers/nuclei in black. Digital images of whole slides were captured and annotated for computer assisted morphometry^3^ (**Supplemental Figure 2**). Pressure-perfused labelling dilates vessels being labeled and results in lower % wall thickness for the same severity of remodeling. To quantify this, we also measured unlabeled PA in sections where PV were labeled and vice versa.

### LCMD and LC-MS/MS of Pulmonary Vein and Artery Tissue

A subset of six PVH and six Sham pigs was assembled for LCMD sampling of PA and PV. The area of tissue collected was measured (approximately 500,000 μm^2^ per vessel; **Supplemental Figure 3**) and loading was normalized to tissue sample area.

### Whole Lung Tissue Collection for LC-MS/MS

Prior to ex-vivo vessel labeling, a lung section was harvested with a surgical cutting stapler. Technical issues precluded use of one PVH sample. Samples (n=8 PVH and n=9 Sham) were processed for LC-MS/MS with loading normalized to protein content.

### Bioinformatics Analysis of LC-MS/MS Data

Quality of raw data was evaluated utilizing the NIST MS^25^ quality metrics. MaxQuant software (Max Plank Institute of Biochemistry, Martinsried, Germany) was configured to use either UniProt porcine or human protein sequence database. Reversed sequences of the proteins were appended to the database for estimating false discovery rates (FDRs). MaxQuant reported protein intensities using an overall protein group FDR of ≤ 0.01.^26^ An in-house R script performed differential expression analysis.^27^ Normalized intensities between groups were modeled using a Gaussian-linked generalized linear model and *p* values were FDR corrected using the Benjamini–Hochberg–Yekutieli procedure. Protein groups with a FDR p value of <0.05 and absolute Log_2_ fold change of at least 0.5 (≈1.4 fold change) were considered DEP.

### Proteomic Data Analysis using Ingenuity Pathway Analysis Software

The Log_2_ fold changes, p-values, FDR p values and gene symbols for the DEP were analyzed in IPA to identify key canonical pathways and PUSR altered in experimental Group 2 PH versus Sham. For key canonical pathways and PUSR, an activation *z* score is computed to infer likely activation or inhibition states.^28^ A z-score is not calculated if there is insufficient knowledge base on activation status implications of DEP in the pathway. For canonical pathways and PUSR, we focus on rigorous requirements for statistical significance (FDR corrected p value < 0.01). For PUSR, we further require an absolute value of z-score > 2.0. See supplemental data for pathways/regulators meeting less rigorous statistical metrics.

### Human Pulmonary Vascular Remodeling

From our previous study of pulmonary vascular remodeling in autopsy specimens^3^, we selected three Control and three Group 2 PH patients for LCMD of PV and PA followed by LC-MS/MS and bioinformatics as above. Group 2 PH patients with PA and PV remodeling similar to that seen in the porcine model were selected. Given the smaller sample size, designation of DEP used absolute Log_2_ fold change of at least 0.5 and p values < 0.05 and canonical pathways and PUSR report p values without FDR correction.

### Statistical Analysis

Piglets were growing after surgery with expected effects on hemodynamics and vessel morphology. Pigs met ES criteria at different ages. Thus, serial data were summarized with medians and interquartile ranges (IQR), and were compared between groups (Sham vs PVH) and by time (baseline vs ES) using mixed models adjusting for repeated data in each pig, including the group*time interaction, and allowing for heterogeneous variance across the group*time categories. Distributions appeared normal (assessed visually) for these data and thus raw data were used in models.

The percent (%) wall thickness data were skewed (visual inspection) and were log transformed for analysis. Mixed models were used to compare vessel morphology between groups (Sham vs PVH) and vessel labeling status (along with group*labeling interaction), accounting for correlated data within each pig (hundreds of vessel measurements within each pig), and also allowing for heterogeneous variance across the group*label categories. These models were used to estimate the mean % wall thickness (along with the standard error of the mean, SEM) on the log scale, and the mean estimates were back-transformed for ease of interpretation. A p-value <0.05 was considered statistically significant. In human subjects, histologic characteristics for each subject were calculated from the average of all measurements performed in that subject. These and clinical characteristics were compared to Control subjects with students t-test. Analyses were performed using SAS version 9.4 (SAS Institute Inc., Cary, NC).

## RESULTS

### Pulmonary hemodynamics in experimental Group 2 PH

Venograms confirmed obstruction of the posterior PV confluence in PVH vs Sham (**Figure 1**) pigs. In PVH pigs, ES criteria were met at one (n=6), two (n=1, hemodynamic data only), three (n=2) or four (n=1) months after surgery. Sham pigs were sacrificed at one (n=4), three (n=4) and four (n=1) months after surgery. The time from surgery to ES did not differ significantly by group (**Table 1**).

**Figure 1.**
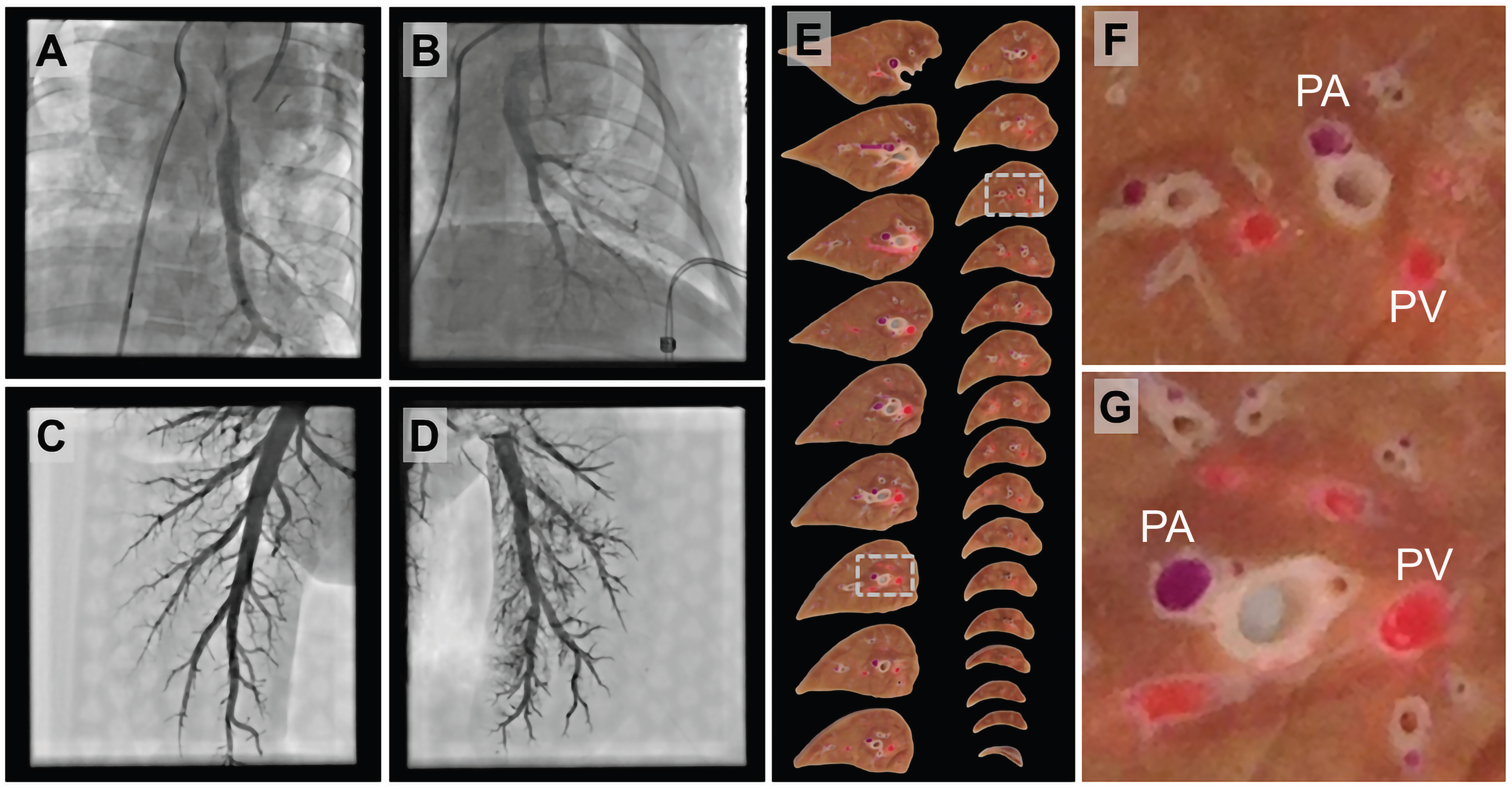
Experimental PH venograms, arteriograms and gross lung specimens. In-vivo venograms confirming an obstructed (A) and patent (B) common posterior vein in an experimental and sham pig, respectively are shown. Ex-vivo, post-mortem angiograms displaying barium-agarose solution in pulmonary arteries of right lung (C) and pulmonary veins of left lung (D) are shown. In E are shown serial, transversely sliced gross specimens of right posterior lung lobe with microbead-aragose perfusion labeling for histological section sampling. In F and G, are expanded images of lung sections displaying purple (pulmonary arteries, PA) and red (pulmonary veins, PV) microbeads-agarose solution.

**Table 1.**
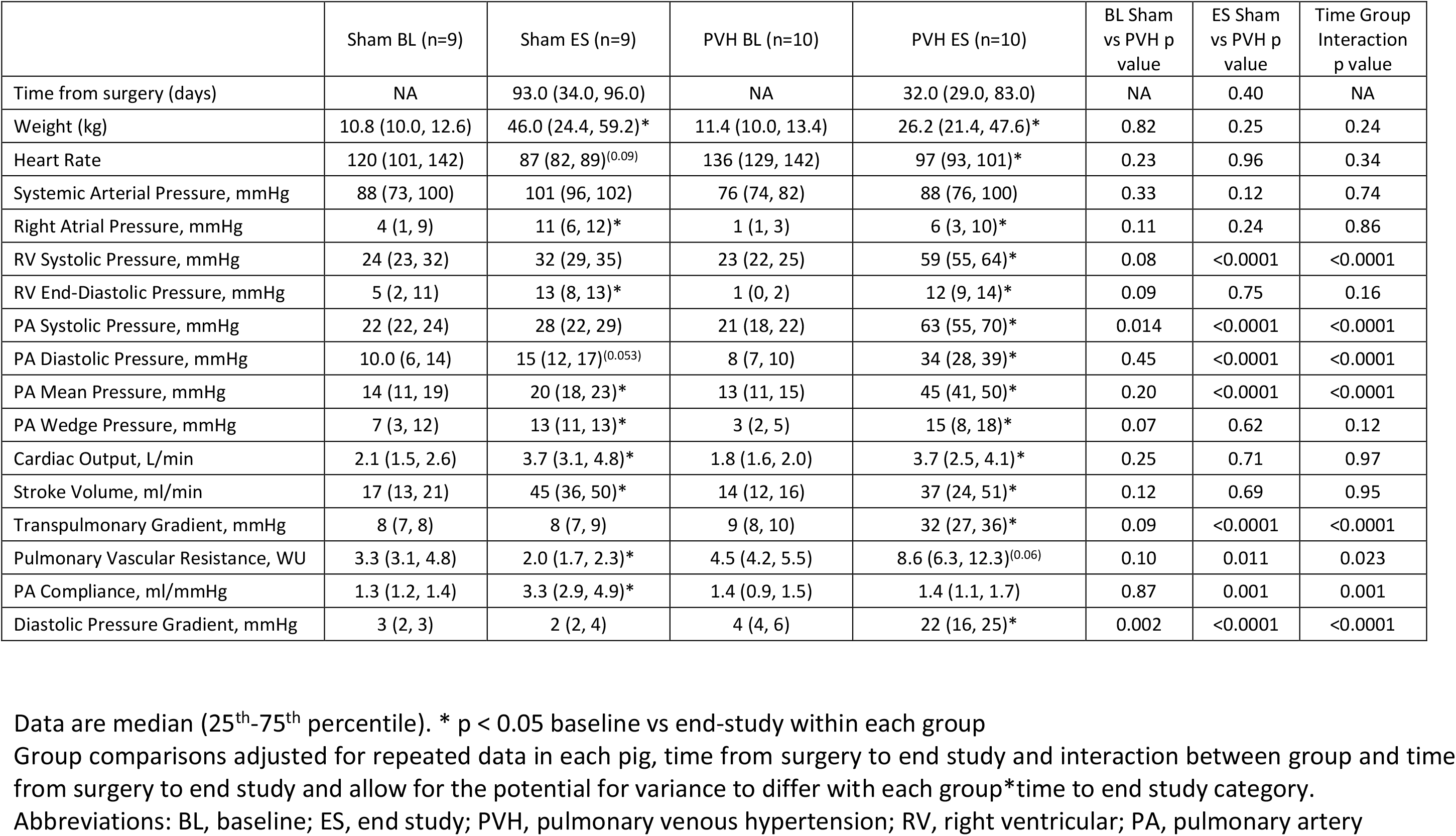
Hemodynamic Characterization of Experimental Group 2 PH.

At baseline (**Table 1**) there were no physiologically significant differences between groups. In Sham pigs, weight, RA, RV end-diastolic, PA mean and PCW pressures, SV, CO and cPA increased from baseline to ES while PVR decreased; all reflecting growth related changes. As compared to Sham pigs, the increases in RV systolic and PA pressures, TPG, DPG and PVR over time were significantly greater and the increase in cPA over time was significantly less in PVH pigs (Table 1, time group interaction p < 0.05 for all).

In the conscious PA measurement subgroup of pigs (**Supplemental Figure 4**), conscious mean PA pressures were 3-fold higher in PVH (n=3; 51±6 mmHg) as compared to Sham (n=4; 17±1 mmHg, p<0.001 vs PVH), differences that exceeded the corresponding measurements under general anesthesia in PVH (31±3 mmHg; p=0.008 vs conscious) and Sham (14±1, p=0.024 vs conscious).

### Pulmonary vascular remodeling in experimental Group 2 PH

Post-mortem perfusion labeling of the PA and PV (**Figure 1**) allowed measurement of labeled PA (n= 11,735), unlabeled veins in the lungs with PA labeling (n= 5,268), labeled PV (n= 13,080) and unlabeled arteries (n=6,727) in the lungs with PV labeling. The labeled and unlabeled PA and PV wall thicknesses were both greater in PVH than Sham (**Table 2, Figure 2**). As expected, the % wall thicknesses were smaller in labeled vessels vs unlabeled vessels due to the impact of pressurized perfusion labeling on lumen diameter but this effect was not significantly different in Sham and PVH groups (group*labeling interaction p>0.05). The PA remodeling was primarily in the media and cells in media stained brown/red (consistent with prominent smooth muscle cell phenotype) whereas the adventitia surrounding the PA and the PV wall thickening had more pink/red staining consistent with a prominent fibrotic component (**Figure 2 and Supplemental Figure 5**)

**Figure 2.**
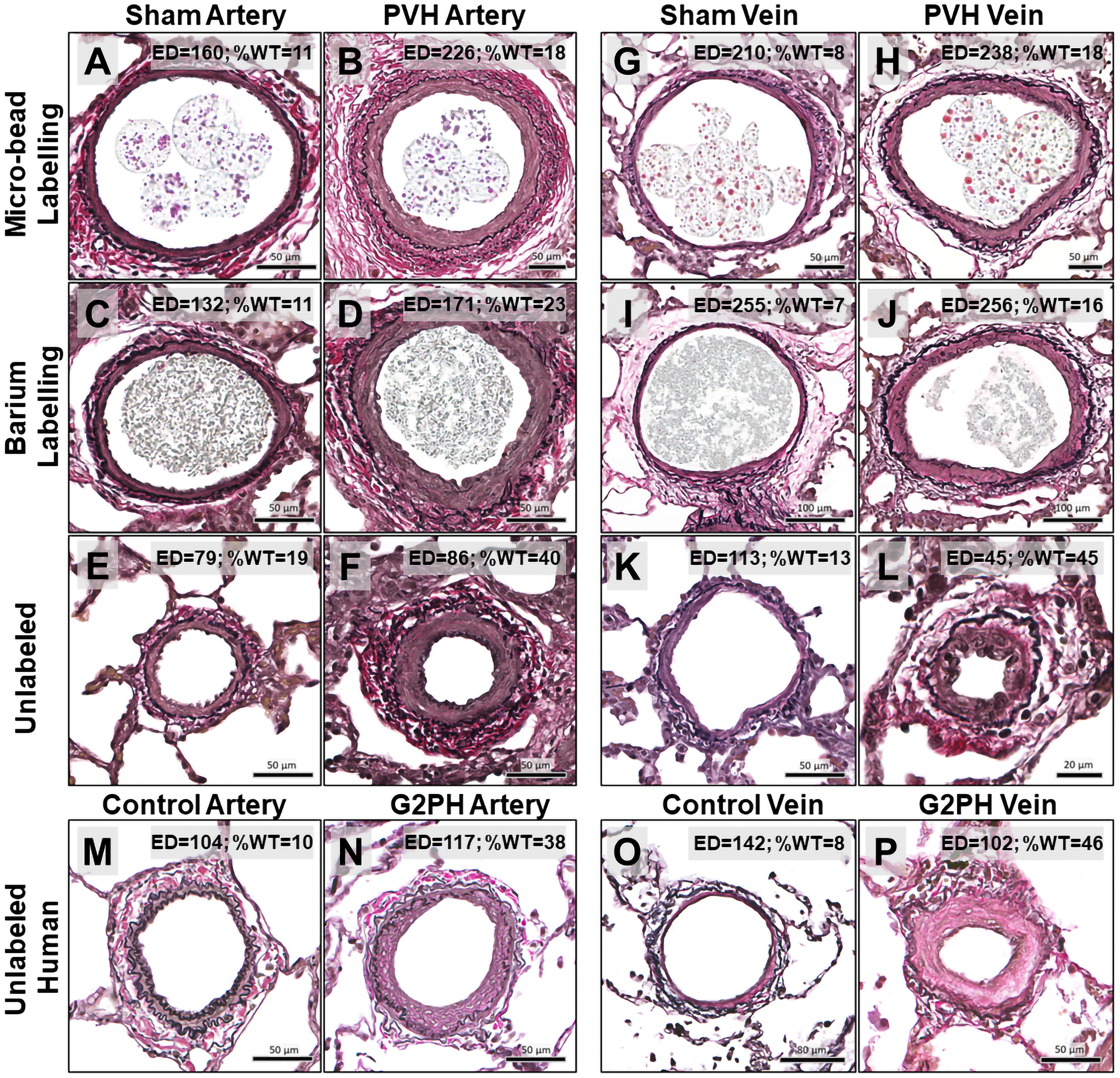
Pulmonary vascular remodeling in experimental PVH and human Group 2 PH. Verhoeff-van Gieson stained histomicrographs of representative pulmonary vessels with % wall thickness (%WT) approximating the median value in each study group. Rows represent the labelling status and columns represent the group and vessel type. In pigs, A-F show purple microbeads (A.B), barium (C,D) or no labeling (E,F) in porcine arteries. In pigs, G-L show red microbeads (G,H), barium (I,J) or no labeling (K,L) in porcine veins. In humans, M-P show unlabeled Control and Group 2 PH arteries (M,N) and veins (O,P). Abbreviations: ED, external diameter (μm); G2PH, group 2 pulmonary hypertension; PVH, pulmonary venous hypertension and % WT, percent wall thickness.

**Table 2.** Pulmonary Vascular Remodeling in Experimental and Human Group 2 PH.

|  | Log Estimated Mean %<br>WT (SEM) | Estimated Mean % WT* | Group p<br>value† | Label p<br>value‡ | Group*Label<br>Interaction p |
| --- | --- | --- | --- | --- | --- |
| **Arteries** |  |  |  |  | 0.20 |
| Sham Label | 2.34 (0.07) | 10.3 |  | 0.001 |  |
| PVH Label | 2.99 (0.10) | 19.9 | <0.0001 | 0.004 |  |
| Sham No Label | 2.67 (0.05) | 14.5 |  |  |  |
| PVH No Label | 3.60 (0.15) | 36.5 | <0.0001 |  |  |
| Human Data (No Label) |  | Mean % WT |  |  |  |
| Human Control |  | 13±1 |  |  |  |
| Human HF+PH |  | 33±12 | 0.049# |  |  |
| **Veins** |  |  |  |  | 0.06 |
| Sham Label | 2.03 (0.06) | 7.6 |  | <0.0001 |  |
| PVH Label | 2.65 (0.15) | 14.2 | 0.002 | <0.001 |  |
| Sham No Label | 2.60 (0.03) | 13.4 |  |  |  |
| PVH No Label | 3.68 (0.15) | 39.8 | <0.0001 |  |  |
| Human Data (No Label) |  | Mean % WT |  |  |  |
| Human Control |  | 12±3 |  |  |  |
| Human HF+PH |  | 33±5 | 0.004# |  |  |

\* Back transformed from model estimated mean, † vs respective Sham, ‡ vs respective No Label
#Human data are group mean ± standard deviation with p values calculated from Students t test which does not account for the measurement of multiple arteries and veins in each subject.
Abbreviations: HF, heart failure; PH, pulmonary hypertension; WT, wall thickness; PVH, pulmonary venous hypertension; SE, standard error

### Pulmonary vascular proteomics in experimental Group 2 PH

#### Differentially Expressed Proteins (DEP)

In PVH and Sham pigs, there were 2093 proteins identified in PA samples, 2085 in PV and 5968 in WL (**Figure 3**). In PA and PV, there was near complete overlap of detected proteins with 94% of protein in PV also detected in PA and 93% of proteins detected in PA also detected in PV.

**Figure 3.**
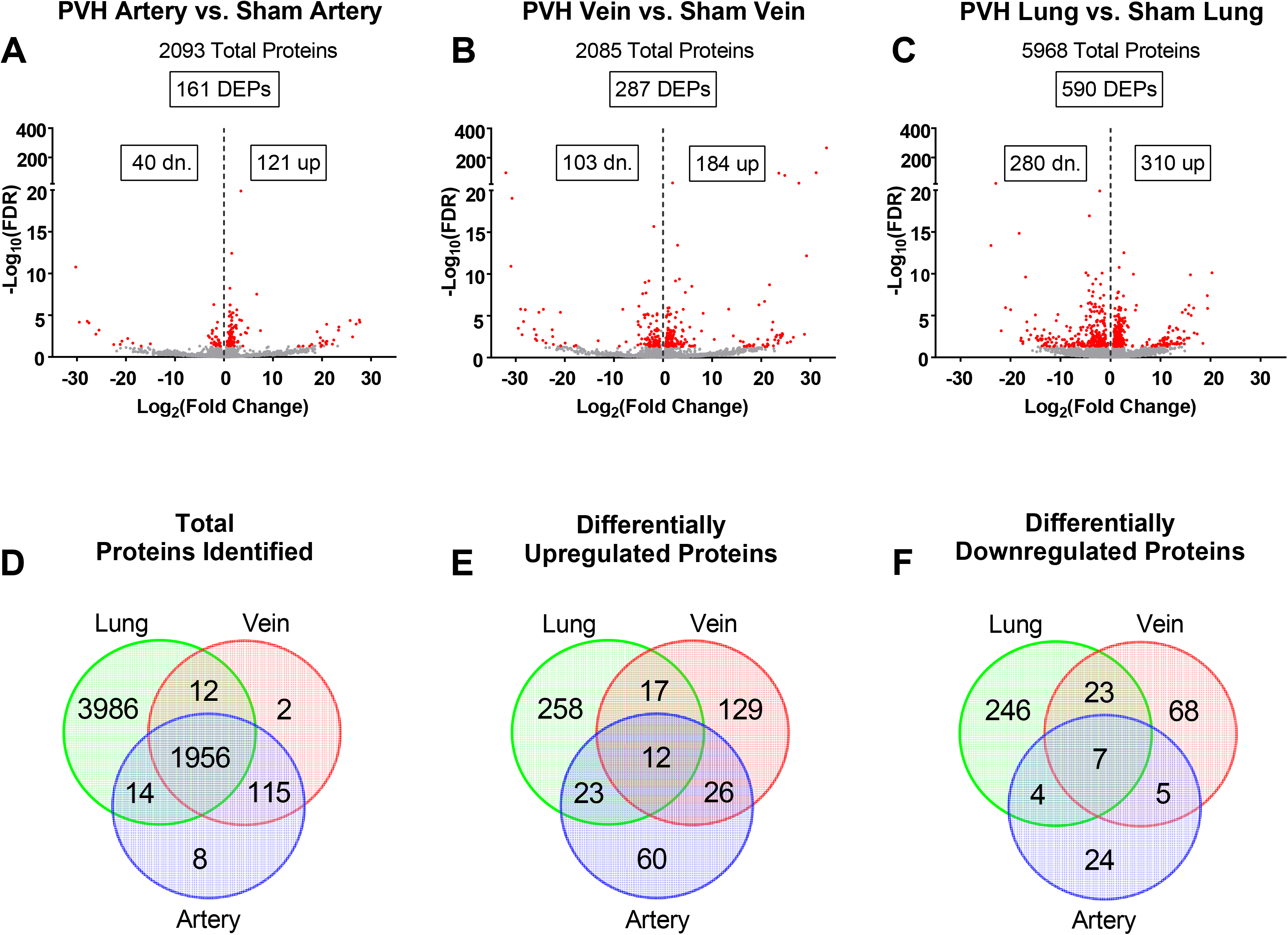
Proteomic remodeling in experimental PVH and human Group 2 PH. Volcano plots (A-C) displaying proteins in arteries (A), veins (B) and whole lung (C) porcine specimens. The −Log_10_ of the Benjamini-Hochberg (BH) false discovery rate corrected p value is on the *y*-axis and the Log_2_ fold change of protein expression is on *x*-axis; significantly altered proteins are shown as red dots. In porcine experimental PVH, Venn diagrams (D-F) display overlapping total (D), up-regulated (E) and down-regulated (F) proteins in whole lung, vein and artery.

In PA, there were 161 DEP (**Figure 3**) in PVH (121 up-regulated and 40 down-regulated; **Supplemental Table 1**). In PV samples, there were 287 DEP in PVH (184 up-regulated and 103 down-regulated, **Supplemental Table 1**). Of the 448 DEP in PA and/or PV, only 50 (11%) were differentially expressed in both PA and PV. In WL samples, there were 590 DEP in PVH (310 upregulated and 280 down-regulated in PVH; **Supplemental Table 1**). Of the 161 DEP in PA, 46 (29%) were also DEP in WL. Of the 287 DEP in PV, 59 (21%) were also DEP in WL. Only 19 proteins were DEP in PA, PV and WL (**Supplemental Table 1**).

#### Select Proteins of Interest in Pulmonary Vascular Biology and PH Therapeutics

While not detected in LCMD samples, bone morphogenic protein receptor 2 (BMPR2) was down regulated in WL (Log_2_ fold change −14.5; FDR p=0.010) in PVH versus Sham. Phosphodiesterase 5a, prostacyclin synthase and soluble guanylyl cyclase proteins were detected in PA, PV and WL but were not DEP. Endothelin receptor protein was not detected in PA, PV or WL. Endothelin converting enzyme protein was detected in WL only and not DEP. Endothelial nitric oxide synthase (NOS3) protein was down regulated in PV (Log_2_ fold change - 30.8; FDR p < 0.0001) and WL (Log_2_ fold change −16.7; FDR p <0.0001) and tended to be down regulated in PA (Log_2_ fold change −9.3; p = 0.080, FDR p 0.33). Receptors involved in adrenomedullin signaling in the endothelium (CALCRL, RAMP-2 and RAMP-3) were only detected in WL and were (CALCRL Log_2_ fold change −14.6; FDR p=0.008; RAMP-3 Log_2_ fold change −12.2; FDR p=0.047) or tended to be (RAMP-2 Log_2_ fold change –5.3, p=0.046; FDR p=0.21) down regulated in PVH.

#### Proteins Reflecting Endothelial to Mesenchymal Transition (EndoMT)

Of protein biomarkers of EndoMT from the literature,^19–24^ 216 were present in our PA, PV or WL protein set; 73 of which were DEP in PVH in some sample type. Of the EndoMT markers, 15 were DEP in PA (**Figure 4**) with all but two proteins (beta 2 microglobin (B2M) and chloride intracellular channel 4 (CLIC4)) changed in a manner consistent with EndoMT. In PV, there were 30 DEP that were markers of EndoMT and all but two (B2M (above) and Yes-associate protein 1 (YAP-1)) were altered in a manner consistent with EndoMT. Of note, while YAP-1 is believed to be up-regulated in EndoMT, it is also intimately involved in cellular responses to mechanical stress, interacts with VE-cadherin and catenins (all down regulated in PV here) and expression can vary by tissue and cell density.^29^ Of the markers of EndoMT, 52 were DEP in WL with 40 altered in a manner consistent with EndoMT. Collectively, these data indicate the presence of EndoMT (particularly in the PV) in experimental Group 2 PH.

**Figure 4.**
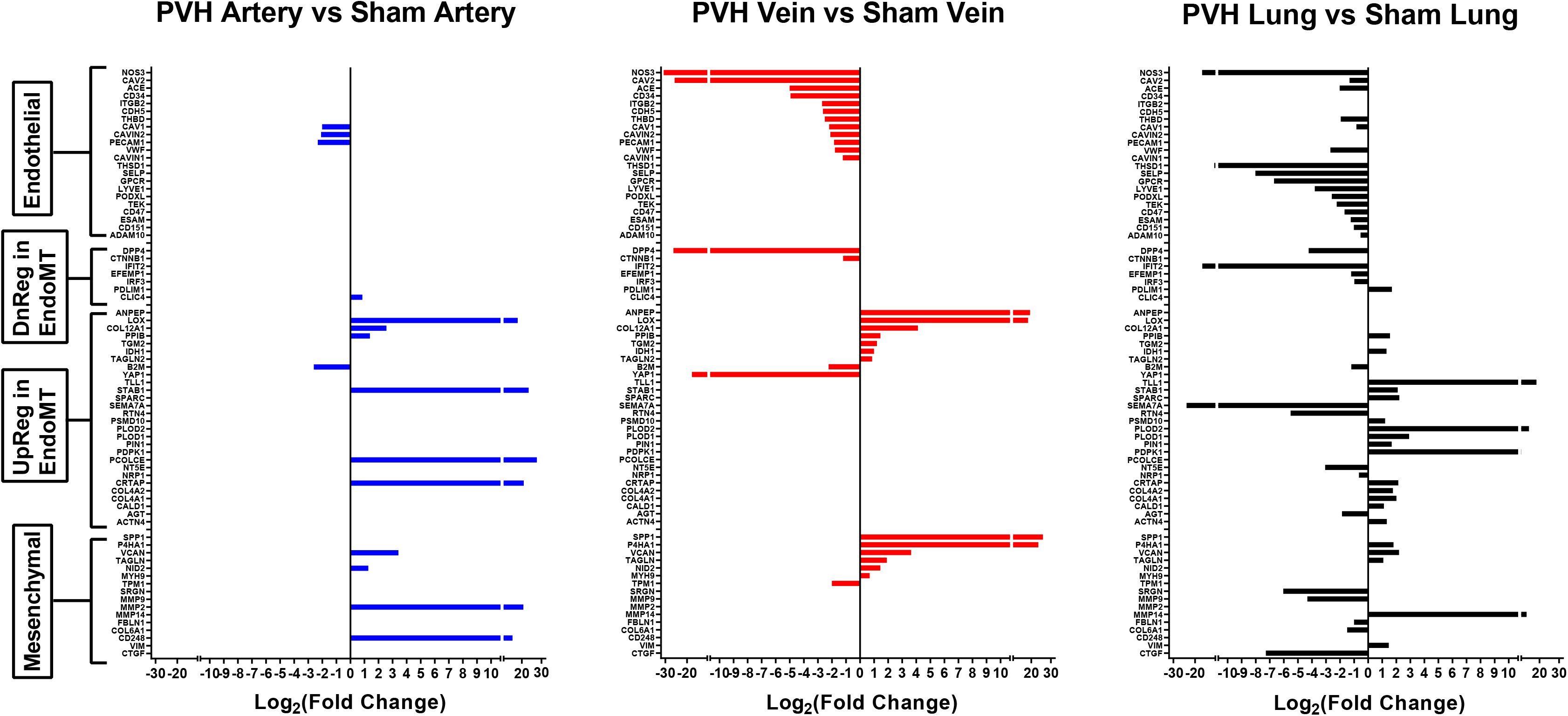
Biomarkers of endothelial–to–mesenchymal transition (EndoMT) in experimental PVH. Bar graphs displaying the log_2_ fold change of differentially expressed proteins reflective of EndoMT (endothelial or other markers down-regulated and mesenchymal or other markers up-regulated in EndoMT) plotted on *x*-axis and protein coding gene symbols on *y*-axis. Abbreviations: DnReg, downregulated; EndoMT, endothelial-mesenchymal transition; PVH, pulmonary venous hypertension; and UpReg, upregulated.

#### Canonical Pathways altered in Pulmonary Arteries

In PA from PVH vs Sham pigs, there were nominally significant differences in 49 canonical pathways (p value < 0.05; **Supplemental Table 2**) but only three pathways were significant with a FDR corrected p value < 0.01 (**Figure 5**). These included the related eukaryotic initiation factor 2 (EIF2), regulation of eukaryotic translation initiation factor 4 and p70S6K signaling (eIF4-p70S6K) and mammalian target of rapamycin (mTOR) pathways. These three pathways were also significantly altered in PV and WL (FDR p value < 0.01 for all). For EIF2, z scores were > 2.0 in PA, PV and WL indicating activation of the pathway. For eIF4-p70S6K and mTOR pathways, z-scores could not be calculated but the majority (≥88%) of DEP detected in each pathway were up-regulated based on Log_2_ fold change values. These pathways are involved in the integrated stress response, regulate protein synthesis in response to a wide range of cellular stressors and have been implicated in pathobiology of other forms of PH.^30–32^

**Figure 5.**
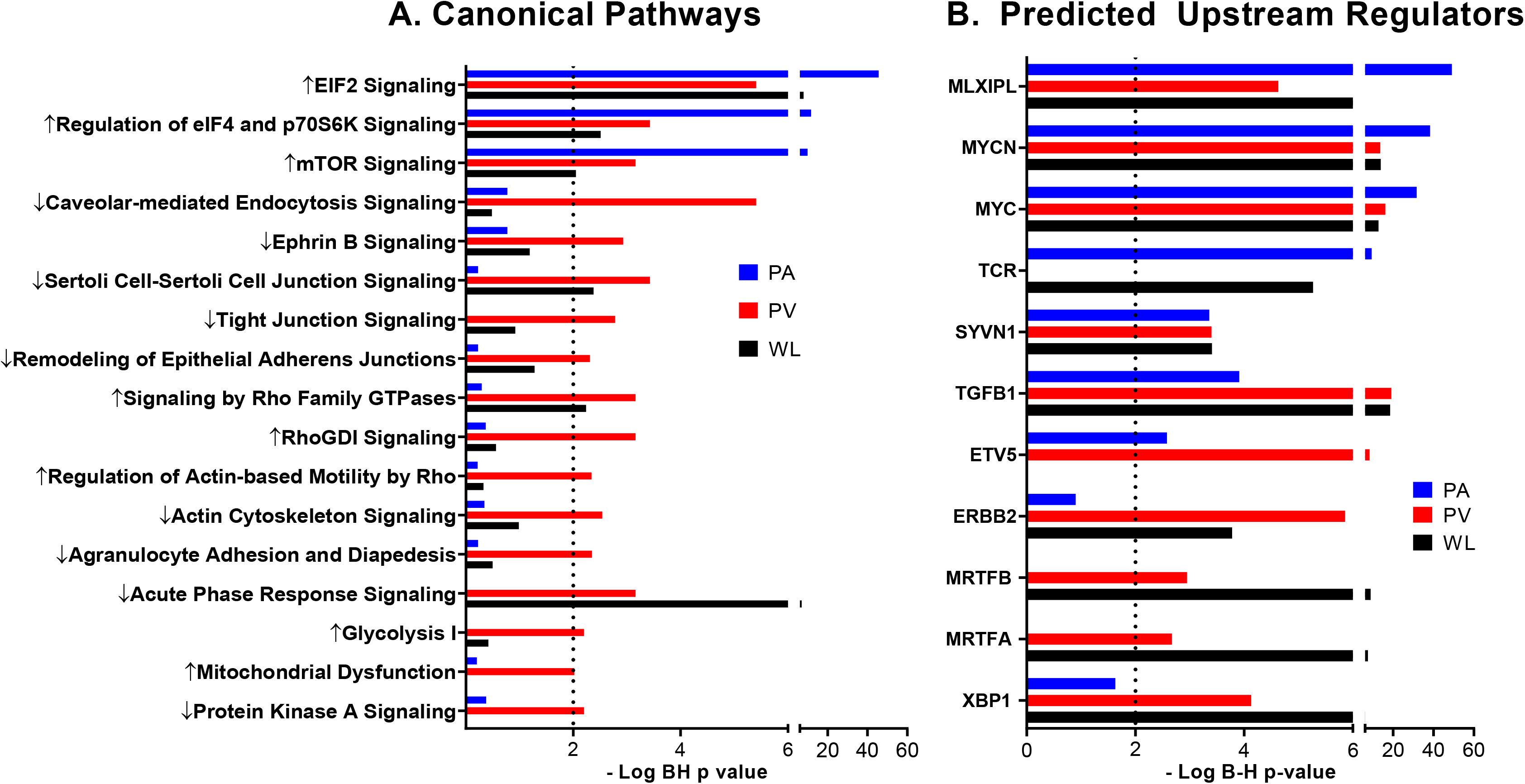
Canonical Pathways and Predicted Up-stream Regulators (PUSR) in pulmonary vessels and whole lung specimens in experimental PVH. Bar graphs outlining the canonical pathways (A) or PUSR (B) on *y*-axis and the −Log_10_ of Benjamini-Hochberg (BH) false discovery rate corrected p value for pulmonary arteries (PA, blue), pulmonary veins (PV, red) or whole lung (WL, black) on the *x*-axis. A missing bar indicates that there was not sufficient overlap in that tissue to generate a p value or fold change. For canonical pathways (A), arrows next to pathway name indicate probable direction of change (↑ activated, ↓ inhibited; see text). Abbreviations: MLXIPL, MLX interacting protein-like; MYCN, N-myc proto-oncogene protein; MYC, Myc proto-oncogene; TCR, T-cell receptor; SYVN1, synoviolin 1; TGFB1, transforming growth factor beta 1; ETV5, ETS variant transcription factor 5; ERBB2, erb-b2 receptor tyrosine kinase 2; MRTFB, myocardin related transcription factor B; MRTFA, myocardin related transcription factor A; and XBP1, X-box binding protein 1.

#### Canonical Pathways altered in pulmonary veins

In PV from PVH vs Sham pigs, there were nominally significant differences in 108 pathways (p value < 0.05; Supplemental Table 2) and 17 pathways which were significant with a FDR corrected p value < 0.01 (**Figure 5**). As above, the EIF2, eIF4-p70S6K and mTOR pathways were altered in PV. Physical forces are known to activate the RhoA/Rho kinase system.^33^ In PV, Signaling by Rho Family GTPases, Rho GDI Signaling and Regulation of Actin-based Motility by Rho were all significantly altered in PVH PV and z-scores were ≥ 0 (albeit < 2), suggesting activation. Rho/Rho kinase signaling has been reported to be up-regulated as an early step in different etiologies of PH, including Group 2 PH.^33, 34^ Consistent with activation of Rho/Rho kinase pathway and its known interactions with the actin cytoskeleton and adherens junction signaling^33^, the Actin Cytoskeleton Signaling pathway was significantly altered and this pathway also tended to be down regulated in WL (FDR p = 0.10, z-score – 0.71). Remodeling of Epithelial Adherens, Sertoli Cell-Sertoli Cell and Tight Junction Signaling pathways were all significantly altered in PV. While z-scores could not be calculated for these three cell-cell interaction pathways, the majority of DEP in these pathways were down regulated including VE-cadherin, catenin A and B and tight junction protein-1 (aka Zonula occludens-1, ZO-1). Activation of Rho/Rho kinase was also consistent with the down-regulation of NOS3 (above) and significant down-regulation of Ephrin B signaling (z-score = −2.3) in PVH PV. The Eph/ephrin proteins principally modify cytoskeletal organization and cell–cell and cell–extra-cellular matrix adhesion^35^ and may contribute to increased endothelial cell permeability, especially with inflammation.

Caveolar-mediated endocytosis signaling was significantly different in PVH PV and while a z-score could not be calculated, caveolin-1 and −2 and caveolar associated protein-1 and −2 were all significantly down regulated (FDR p < 0.01 for all; Log_2_ fold change < −1.0 for all) suggesting deficiency in caveolae in PV as described in other forms of PH^36, 37^ This pathway tended to be altered in PA (p value 0.014, FDR p value 0.169) and WL (p value 0.095, FDR p value 0.327).

Collectively, this bioinformatic analysis suggests activation of Rho/Rho kinases with impairments in endothelial cell actin cytoskeletal signaling, sertoli cell, tight and epithelial adherens junctions, ephrin B and caveloar-mediated signaling in PV. These pathways have been demonstrated to mediate increased pulmonary vascular permeability in other forms of lung injury.^38, 39^

Agranulocyte Adhesion and Diapedesis, and Acute Phase Response Signaling pathways were both significantly altered in PV. While z-scores were not calculable, 70% of the DEP in the Agranulocyte Adhesion and Diapedesis pathway were down regulated, including platelet endothelial cell adhesion molecule (PECAM1). For Acute Phase Response Signaling, the z-score was – 1.0 and 90% of the DEP were down regulated, including VonWillebrand factor (vWF) and this pathway was also down regulated in WL (FDR p < 0.0001, z-score −1.3). Other significantly altered pathways in PV include glycolysis (FDR p< 0.007, z-score 1.0), protein kinase A signaling (FDR p < 0.01, z-score −1.3) and mitochondrial dysfunction (FDR p< 0.007, z score not calculated but 78% of DEP up-regulated).

#### Canonical Pathways altered in WL samples

In WL specimens, there were nominally significant differences in 90 pathways (p value < 0.05) and 27 pathways which were significant with a FDR corrected p value < 0.01 (**Supplemental Table 2**). Confirming findings in PV, Sertoli Cell-Sertoli Cell Junction Signaling, Clathrin-mediated Endocytosis Signaling, Signaling by Rho Family GTPases and RhoA Signaling were all significantly altered in WL. Additionally, several related or redundant pathways were or tended to be altered including eNOS signaling (FDR p < 0.02, z-score −1.7), Remodeling of Epithelial Adherens Junctions (FDR p 0.052) and Ephrin B signaling (FDR p 0.06) Additional significantly altered pathways dealt with coagulation, inflammation, metabolism, protein synthesis, the unfolded protein response and aldosterone signaling (**Supplemental Table 2**).

#### Predicted Up-stream Regulators (PUSR)

Highly significant (FDR corrected p value < 0.01 and absolute value of z-score > 2.0) PUSR in PVH included six in the PA, 22 in the PV and 32 in WL (**Supplemental Table 3**). Of the top five (smallest FDR p value and activated) PUSR in PA (**Figure 6**), the top 3 proteins (MLX interacting protein like (MLXIPL), MYCN Proto-Oncogene, BHLH Transcription Factor (MYCN) and MYC Proto-Oncogene, BHLH Transcription Factor (MYC) were also among the top 5 activated PUSR for PV and/or WL. These transcription factors belong to Myc/Max/Mad family of transcription regulators which regulate cellular proliferation and apoptosis and have been implicated in cancer^40^ and inflammation related PA remodeling.^41^ The T-cell receptor (TCR) protein was among the Top 5 PUSR in PA, was a significant PUSR in WL and has been implicated in Group 1 PH with up-regulation suggesting a failure of regulatory T cell activity to control vascular endothelial injury.^42^ The fifth top PUSR in PA was E3 ubiquitin-protein ligase synoviolin (SYVN1) which was also a statistically significant PUSR in PV and WL and is involved in endoplasmic reticulum (ER)-associated degradation and removal of unfolded proteins, including tumor suppressor gene p53.^43^ Notably, p53 deficiency is linked to hypoxia mediated pulmonary vascular remodeling^44^ and synoviolin sequesters and metabolizes p53, negatively impacting its intracellular level and biological function.^43^

**Figure 6.**
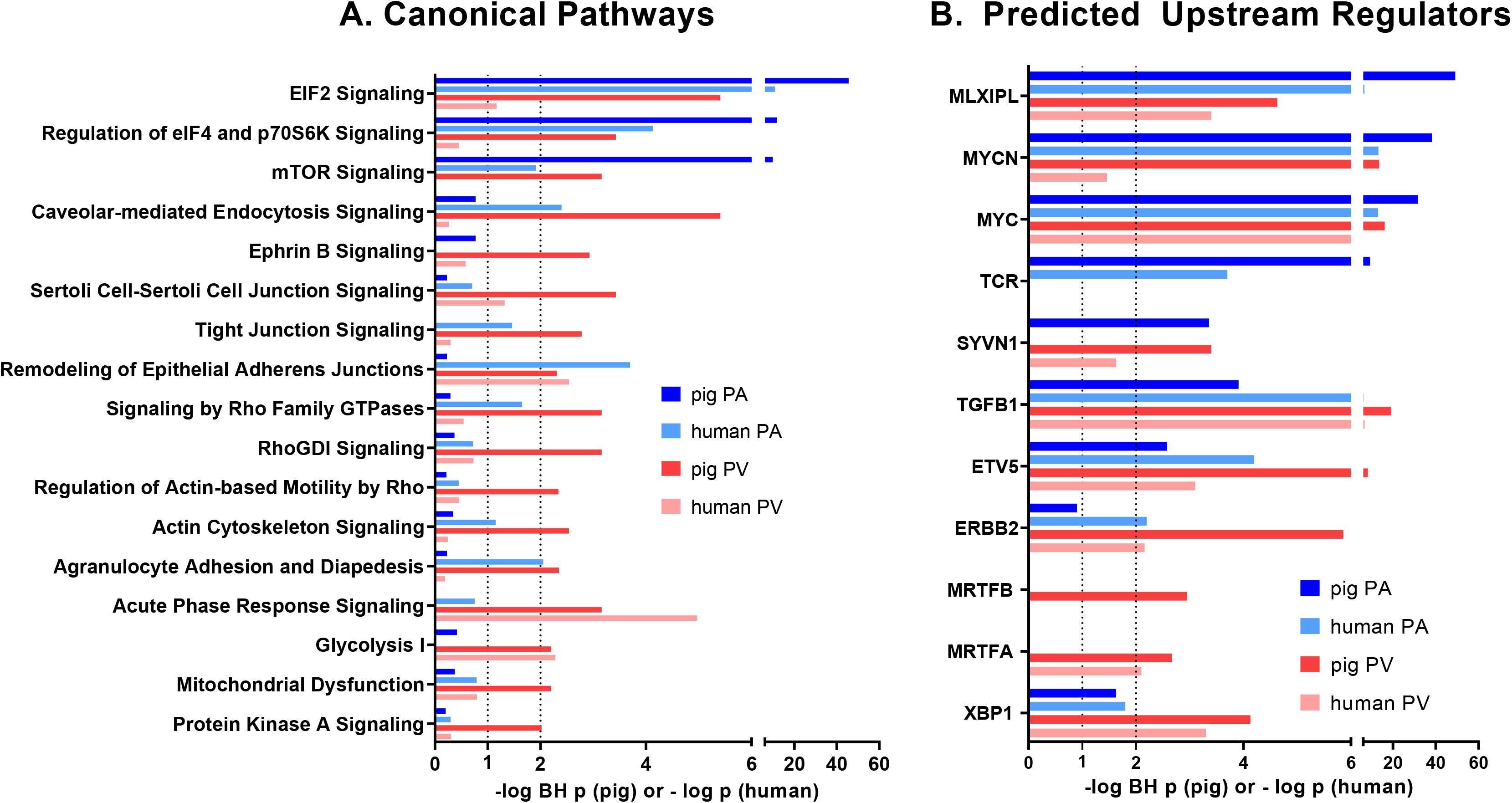
Canonical Pathways and Predicted Up-stream Regulators (PUSR) in human Group 2 PH pulmonary arteries and veins. Bar graphs outlining the canonical pathways (A) or PUSR (B) on *y*-axis and the −Log_10_ p value for human (light blue) pulmonary arteries (PA) and human (pink) pulmonary veins (PV). The findings in porcine PA (dark blue) and PV (red) are duplicated here for reference. For porcine tissue, the −Log_10_ of the BH false discovery rate corrected p value is shown. A missing bar indicates that there was not sufficient overlap with proteins in that tissue to generate a p value or fold change. Abbreviations as in Figure 5.

Transforming growth factor beta (TGF-β) was the most significant PUSR in PV and was also significantly activated in PA and WL. TGF-β is well recognized as a key driver of EndoMT and the pathophysiology of several forms of PH.^45^ Other PUSR in PV include ETS translocation variant 5 protein (ETV5) which is a transcription factor whose gene was up-regulated in lung tissue from Group 1 PH and is functionally associated with BMPR2 signaling and cell proliferation.^10^ Epidermal growth factor receptor 2 (ERBB2, aka HER2) is a key regulator of cell proliferation and survival in cancer, was the 5^th^ top PUSR in PV and has previously been implicated in the pathobiology of Group 1 PH.^46^

In WL samples, two myocardin-related transcription factors (genes MRTFB and MRTFA) were PUSR in WL (and significant PUSR in PV), are stimulated by Rho GTPases, involved in vascular smooth muscle cell hypertrophy and dis-inhibited by loss of SMAD due to chronic TGF-β activation in Group-1 PH.^47^ X-box binding protein 1 (XBP1) was the final top 5 PUSR in WL (and a significant PUSR in PV), functions as a transcription factor during endoplasmic reticulum stress by regulating the unfolded protein response and can function via growth factor signaling pathways to regulate endothelial cell proliferation and angiogenesis.^48^

### Pulmonary vascular remodeling and proteomics in Human Group 2 PH

The HF and Control patients were elderly (82±4 versus 73±17 years old; p=0.45) and all female. In HF, the PA systolic pressure was 58±15 mmHg and the EF was 54±11% (echo unavailable in Controls). By design, the severity of PA and PV remodeling was similar to that observed in the porcine model (Figure 2 and Table 2). The time from formalin fixation to LCMD in human samples averaged 4,564 ± 1922 days versus < 500 days in the pigs. There were 833 proteins detected in human PA (634 (76%) also seen in pig) and 824 proteins in human PV (633 (77%) also seen in pig). As expected with the smaller sample size and less robust protein coverage, there were fewer differentially expressed proteins (100 in PA and 96 in PV) in the human PVH versus Control specimens. Of EndoMT biomarkers, nine were DEP in PA with 6 changing in a direction consistent with EndoMT. In PV, 10 EndoMT biomarkers were DEP, all changing in a manner consistent with EndoMT.

All three of the canonical pathways highly significant in the porcine PA were also significant in human PA (Figure 6). In PV, 5 of the 17 canonical pathways which were highly significant in porcine PV were (n=4) or tended (n=1; p value ≤ 0.1) to be significant in human PV. In human PA and PV, there was nearly complete overlap of pig and human findings for the top PUSR in both PA and PV (Figure 6).

## DISCUSSION

The porcine PVH model recapitulated hemodynamic and histologic findings in human Group 2 PH with elevated PVR and both PV and PA remodeling. The LCMD and LC–MS/MS proteomics strategy identified a similar number of proteins in PA and PV with nearly complete (94%) overlap. However, there were more DEP and more statistically significant EndoMT biomarkers, altered canonical pathways and PUSR identified in PV than PA. The proteomic profile of remodeled PA and PV suggested activation of the integrated stress response and mTOR signaling, dysregulated growth, cellular proliferation and anti-apoptotic signaling which was also seen in WL; consistent with the “cancer hypothesis” in PH vascular remodeling.^49^ In PV, bioinformatics also suggested activation of Rho/Rho kinase signaling with altered actin cytoskeletal signaling, disrupted tight junction, adherens junction, ephrin B, and caveolar mediated endocytosis signaling; all indicating disrupted endothelial cell barrier function in PV. Consistent with this, there was clear biomarker evidence of EndoMT in PVH, particularly in PV. Findings in human Group 2 PH samples were generally consistent with those in the porcine PVH model. The proteomic bioinformatics approach used here leverages the tremendous breadth of existing knowledge related to pulmonary vascular biology, allows generation of highly plausible mechanistic information from a single experiment using < 25 μg of tissue per vessel per subject and suggests multiple potential therapeutic targets to oppose pulmonary vascular remodeling in Group 2 PH.

### Porcine Model of Group 2 PH

Our hemodynamic findings under anesthesia are consistent with previous studies using this model.^16–18, 50^ The conscious PA pressure measurements performed here better demonstrate the severity of PH in this model. The severity of PH here is consistent with levels that produced right ventricular failure in the study of Aguero et al^16^ and based on PVR, is somewhat more severe than in most^1, 3^ but not all^51^ humans with HF related combined pre- and post-capillary PH (CpcPH).

### Endothelial Mesenchymal Transition and Pulmonary Vascular Remodeling in Group 2 PH

In human Group 1 PH and representative animal models including mutant BMPR2 genetic models, Ranchoux et al established that EndoMT was present in remodeled PA and linked to alterations in BMPR2 signaling.^21^ During EndoMT, endothelial cells lose their tight and adherens junctions and thus polarity to the endothelium and change from endothelial to mesenchymal cell types with characteristic changes in EndoMT markers.^52^ In large PV proximal (“upstream”) to the PV stenosis in this Group 2 PH model, Kato et al reported loss of PECAM-1, vWF and VE-cadherin by immunostaining and western analysis and increases in mesenchymal markers of EndoMT and TGF-β.^50^ Our findings confirm and extend the study of Kato et al with significantly lower expression of these same endothelial markers in the small pulmonary venules by LC-MS/MS, significant changes in multiple other markers of EndoMT in PV, down-regulation of BMPR2 in WL and bioinformatics which reveal TGF-β, a key regulator of EndoMT, as the most significant activated PUSR in the PV. EndoMT has also been demonstrated in the remodeling of venous conduit grafts after exposure to arterial level pressures, supporting its role as a maladaptive response to increased transmural pressure loading in veins.^53^

Multiple stressors are believed to initiate the EndoMT process in Group 1 PH and atherosclerosis^52^ including hypoxia, inflammation and mechanical stress induced loss of endothelial junctions as is seen with acute stress failure due to PVH.^1, 54^ Here we also show evidence of enhanced Rho/Rho kinase signaling which further contributes to loss of tight and adherens junctions and caveolae signaling and thus, promotes EndoMT.^52^

While EndoMT has been demonstrated in PA samples from human and experimental Group 1 PH, evidence for EndoMT in PA remodeling here was weaker, with fewer biomarkers of EndoMT altered in PA samples. This may suggest that the PV are more susceptible to the levels of mechanical stress provided by HF related PVH given their developmental and structural differences^4^. Many of the canonical pathways identified here in PA and PV from experimental Group 2 PH were also identified with a proteomic bioinformatics approach in PA smooth muscle cells harvested from patients with Group 1 PH.^13^

### Dysregulated Growth and Pulmonary Vascular Remodeling in Group 2 PH

The EIF2/eIF4/mTOR activation and the PUSR including the Myc/Max/Mad family of transcription regulators and synoviolin seen in remodeled PA and PV are consistent with the integrated stress response, dysregulated growth and the “cancer theory” of adverse pulmonary vascular remodeling in PH. The metabolic derangements (enhanced glycolysis and mitochondrial dysfunction) seen in the canonical pathway analysis in PV and the bioinformatics analysis predicting ERBB2 as a PUSR in PV are further evidence of the hyper-proliferative anti-apoptotic phenotype in PV.

The prominent venous remodeling in PVH here and in human HF^3^ is reminiscent of pulmonary veno-occlusive disease (PVOD). Interestingly, mutations in the gene (EIF2AK4) coding for the amino acid deficiency sensing kinase (GCN2) in the integrative stress response have been found in familial and sporadic cases of PVOD, suggesting that the integrated stress response is prominently involved in venous remodeling in PVOD.^55^ While the mutations of EIF2AK4 in PVOD are *loss of function* mutations, downstream proteins activated as part of the integrated stress response (CHOP and HO-1) are paradoxically increased in human and experimental PVOD with loss of GCN2.^56^ Consistent with this, our findings also suggest activation of the integrated stress response in PVH induced PA and PV remodeling.

### Therapeutic targets

There are few therapies for HF with preserved ejection fraction^57^ to prevent PVH and Group 2 PH. Studies of PH therapies targeting endothelin and NO-cGMP signaling in HF have shown mixed effects with some signals of benefit in HF with reduced but not generally in HF with preserved ejection fraction.^2, 57^ In the present study, there was no up-regulation of PDE5 in the pulmonary vasculature and no clear evidence of activation of endothelin signaling here in PA, PV or WL. This suggests important and fundamental differences in the biology of Group 1 and Group 2 PH that will require distinct treatments.

The canonical pathways and PUSR implicated in PV and PA remodeling here provide further support for different approaches, many already suggested as potential therapies in Group 1 PH. mTOR inhibition with or without concomitant platelet derived growth factor inhibitors may reduce EndoMT by induction of VE-cadherin expression or anti-proliferative effects in Group 1 PH^21, 32^ and our data would support potential benefit in Group 2 PH. Of note, mTOR inhibition has potential favorable effects on right ventricular structure and function as well.^30^ Opportunities and barriers to development of other methods to oppose EndoMT in cardiovascular disease have been well described.^19^ Rho-kinase inhibitors have already been tested and showed benefits in human and experimental Group 2 PH^33, 58^ and the current data provide further support for this strategy. Novel strategies to enhance caveolin in animal models of PH had dramatic beneficial effects and caveolin deficiency was clear in PV here.^37^ A number of therapeutics targeting the integrated stress response are available or being tested.^55^

### Generalizability to the human

Differences in time from death to formalin fixation, time from formalin fixation to LCMD, age, sex, stage of disease (early in pigs and likely late in human) and differences in the comparator non-PH group (aged human non-HF controls vs completely normal sham operated pigs) may all contribute to fewer and/or different proteins detected in the human versus pig specimens. None-the-less, while there was less robust protein recovery from the human autopsy specimens, the bioinformatics findings in human Group 2 PH were largely similar in to those from the porcine PVH model, adding confidence that the targets identified in pigs are worthy of further study and steps toward therapeutic translation.

### Limitations

While we examined consistency across sampling sites and with human tissue, specific DEP within pathways, consistency in canonical pathway and PUSR analyses and consistency with previous studies, we did not perform independent validation for any one pathway with other methodologies. The small tissue volume from LCMD limits use of alternate methodologies. We did not perform assessments of different time points in the model or band adult pigs. We also did not band female pigs although the human specimens were all from females with similar findings to pigs.

### Conclusions

Based on pulmonary vessel specific proteomic bioinformatics in experimental PVH, we found that PVH produces significant vascular remodeling that is associated with disrupted endothelial barrier function and EndoMT in PV where as in PA and PV, there is activation of the integrated stress response and mTOR signaling and evidence of dysregulated growth. Our findings provide new insight into the pathobiology of Group 2 PH and support multiple novel therapeutic targets to oppose pulmonary vascular remodeling in HF.

## Acknowledgment

We thank Carrie Jo Holtz-Heppelmann for assisting with the methods of mass spectrometry.

