## Supplemental for "Histologic and Proteomic Remodeling of the Pulmonary Veins and Arteries in a Porcine Model of Chronic Pulmonary Venous Hypertension"

**Supplementary Appendix**


| Context | Page |
|---|---|
| Supplemental Methods | 2-7 |
| Supplemental Table 1. Differentially Expressed Proteins | 8-31 |
| Supplemental Table 2. Significantly Altered Ingenuity® Canonical Pathways at Uncorrected P < 0.05 Level. | 32-37 |
| Supplemental Table 3. Ingenuity® Predicted Upstream Regulators with FDR p Value <0.01 (-log p > 2.0) AND I Z-score I ≥ 2.0 | 38-41 |
| Figure 1. Study Design | 42 |
| Figure 2. Whole slide digital microscopic scanning of lung specimens with annotation of pulmonary vessels with quantitative histomorphometry measurement | 43 |
| Figure 3. Workflow of laser capture microdissection of pulmonary arteries and veins using ZEISS PALM Microbeam System | 44 |
| Figure 4. Conscious Pulmonary Artery Pressure Assessment | 45 |
| Figure 5. Histological demonstration of Verhoeff-van Gieson (VVG) staining | 46 |
| References | 47 |

**Supplemental Methods:**

**Study design (Supplemental Figure 1)**:

In male domestic piglets (n=23), PVH was produced (n=14) by surgical banding of the left anterior PV and the posterior common PV via a left thoracotomy at age approximately 30 days when weight was approximately 10 kg. Sham animals (n=9) of similar size underwent thoracotomy without PV banding. In the PVH group, two animals were euthanized due to infection, and three animals had unobserved death presumed due to RV failure during the study. A right heart catheterization (RHC) with pulmonary arteriogram and pulmonary venogram were performed at baseline (prior to PV banding or Sham procedure), intraoperatively (confirming band placement) and at one month intervals. One of the PVH pig's unobserved death occurred two days after the two month RHC and hemodynamic data was used for that pig but no tissue was available.

In a randomly selected sub-group (n=7), CardioMEMS PA pressure monitors (Abbott Laboratories, Chicago IL) were implanted at the one month RHC when piglets were large enough to allow placement. Conscious measurements of PA pressure were recorded intermittently. Experimental animals underwent a final RHC and were euthanized with pentobarbital sodium at end study. Criteria for end study included clinical instability (n=4), severe combined pre-capillary and post-capillary hypertension (CpcPH) at monthly RHC (n=5) or 16 weeks post-banding (n=1). Sham pigs were sacrificed in similar fashion and at similar time points to experimental pigs. At end study, pulmonary arteries and veins were specifically labeled using pressurized perfusion with microbead- or barium-agarose solution (below).

**Anesthesia and peri-operative analgesia:**

For thoracotomy, all animals were given intramuscular Telazol (5.0 mg/kg) and Xylazine (1-2 mg/kg) prior to intubation. General anesthesia was maintained with 1.5-3.0% isoflurane. Additional analgesia was provided fentanyl (intravenous bolus of 100 mcg followed by infusion of 2-5 mcg/kg/hr for the duration of the procedure). All animals received Buprenorphine SR immediately following thoracotomy closure. For monthly RHC, animals were administered Carprofen 4 mg/kg subcutaneously for perioperative analgesia. Anesthesia utilized Telazol, Xyalzin and isoflurane as for thoracotomy.

**Pulmonary Vein Banding:**

Briefly, a left lateral thoracotomy at the 5$^{th}$ or 6$^{th}$ intercostal space was performed. After mobilization of the pericardium, the left anterior PV and common posterior PV were bluntly dissected. A 3.5mm plastic cylinder was then applied to the left anterior PV and surgical umbilical tape was fixed around the cylinder and left anterior PV at the pulmonary vein-atrial junction. After fixation, the plastic cylinder was removed, leaving the umbilical tape in place. A 6 mm lumen inflatable vascular occluder (DocXS Biomedical, Ukiah, CA) was then secured via eyelet holes around the posterior common PV. Successful banding was confirmed via intra-operative pulmonary venogram as pilot studies had demonstrated that slight angulation differences in the posterior PV confluence band placement resulted in banding of only the left posterior PV and no significant PH. Two skin incisions were then made over the left side of the animal's mid-back just left of the spine, and the occluder tubing was subcutaneously tunneled through the chest to the mid back. The proximal pocket incision harbored coiled line to account for animal growth, while the distal pocket contained a subcutaneous port (DoxXS Biomedical, Ukiag,CA) allowing for diameter adjustment of the occluder. After confirming cuff inflation, the chest wall and mid-back incisions were closed.

**Hemodynamic evaluation:**

Femoral artery access was obtained via a 5-Fr microcatheter and arterial blood pressure was transduced and recorded. Femoral venous access was obtained and a 7 Fr Swan-Ganz catheter was advanced into the right heart under fluoroscopic guidance. Right atrial (RA), right ventricular (RV), pulmonary artery (PA), and pulmonary capillary wedge (PCW) pressures were transduced and recorded. Cardiac output (CO) was measured using the thermodilution method. Stroke volume (SV) was calculated as CO divided by heart rate (HR). Pulmonary artery capacitance (cPA) was calculated as SV/($PA_{systolic}$-$PA_{diastolic}$ pressure), trans-pulmonary gradient (TPG) was calculated as mean PA-PCW, pulmonary vascular resistance (PVR) was calculated as TPG/CO, and diastolic pressure gradient (DPG) was calculated as $PA_{diastolic}$-PCW.

**Pulmonary Arteriograms and Venograms:**

Following hemodynamic measurements, a Swan-Ganz catheter was advanced into the left posterior PA, the cuff was inflated, 10 mL of iodinated contrast media was injected and the arteriogram was recorded. Immediately following the arteriogram, the balloon was deflated and the venogram was recorded as contrast media flowed toward the left atrium. The Swan-Ganz catheter was then advanced into the right posterior PA and right PA arterio- and venograms were recorded.

**Conscious PA pressure monitoring**:

In a subset (3 PVH and 4 Sham) of animals, at the one month RHC and following angiography, the Swan-Ganz catheter was advanced into the left posterior PA. An exchange length wire was advanced into the left PA via the Swan Ganz distal port. The Swan-Ganz catheter was subsequently exchanged for the PA pressure monitor (CardioMEMS®) deployment catheter after the device had been primed in sterile heparinized saline. The position of the device in the left posterior PA was confirmed via fluoroscopy, and the device was deployed. After detachment, the delivery catheter was removed and device position was confirmed via fluoroscopy. The device was then calibrated with the Swan Ganz catheter per manufacture's guidelines. Conscious PA pressure measurements were made sequentially and compared to temporally similar anesthetized measurements.

**Post-Mortem Pulmonary Vasculature Labeling:**

In the initial three PVH and three Sham animals, a microbead-agarose solution was infused. To prepare the solution, 1 g of either purple (artery) or red (vein) 20-45 µm microspheres (Cospheric LLC, Santa Barbara, CA) was added to 3 mL of 0.1% Tween, emulsified, and then added to 57 mL of 1% agarose solution. To prepare the agarose solution, one g of UltraPure low-melting point agarose (Thermo Fisher Scientific, Waltham, MA) was added to 100 mL of distilled water, and heated to $90^{o}$ C until completely dissolved. The solution was kept at approximately $70^{o}$ C until it was injected. In the remaining 12 animals, 200 g of barium sulfate (Sigma-Aldrich, St. Lois, MO) was added to 200 mL of the 1% agarose solution. After mixing, the solution was kept at $70^{o}$ C until ready to be infused.

After in-vivo measurements were recorded, 100 U/kg heparin was administered. The animal was subsequently euthanized and the heart and lungs were removed *en bloc* via median sternotomy taking care not to damage lung tissue. Organs were gently rinsed with saline, and the pulmonary vessels were identified at the lung hilum. Using an 8-Fr Swan-Ganz catheter, the distal end was inserted into the PA or PV, the balloon was inflated and the vessel was additionally clamped at the catheter insertion site to allow pressure perfusion. Vessels were injected with either a microbead-agarose solution (n=6) where PA and PV could be labeled in the same lung segment (right posterior lobe) or with barium-agarose solution (n=12) where PA in the right lung and PV in the left lung were labeled. Infusion used moderate constant manual pressure in 15 mL increments. In vessels injected with the microbead-agarose solution, the injection was stopped and the vessel was clamped once back-flow of the labeling solution was visualized. In vasculature injected with the barium-agarose solution, the injection was stopped and clamped distal to the incision once the spread of the post-mortem labeling solution appeared similar to the in-vivo arterio- or venogram under fluoroscopic guidance. The lung was then placed in an ice bath to allow the labeling solution to cool and solidify prior to any further post-mortem tissue processing. The process was then repeated injecting the left lung venous system as per the right arterial system.

**Lung Preparation:**

Following vessel labeling, the lungs were distended by intra-tracheal instillation of buffered formalin (under gravity) until the lung margins were defined. The trachea was ligated to maintain distension of the lung. Lungs were fixed in formalin (10 volume equivalents of formalin per volume of lung tissue) for 48-72 hours prior to sectioning. For histologic analysis, blocks were taken from the peripheral regions of only right posterior lobe of 7 (4 PVH, 3 Sham) pigs, and both right and left posterior lobes of 11 (5 PVH and 6 Sham) pigs.

**Quantitative Histomorphometry Analysis:**

All tissue blocks were paraffin embedded, cut at four µm thickness, and stained with hematoxylin and eosinophil (H&E) and Verhoeff-van Gieson (VVG) stains. All stained histology slides were reviewed to select the tissue blocks with adequate samples of pulmonary vessels. One sham pig was excluded due to histologic features consistent with diffuse pulmonary parenchymal disease. Three to seven blocks were selected from each animal for histomorphometry analysis. Digital images of whole slides were captured at 40x magnification with a resolution of 0.25 um per pixel, and scanned into SVS format, using ScanScope AT Turbo (Leica Biosystems Inc., Buffalo Grove, IL, USA). Histomorphometric measurements of the vessels were performed in ImageScope (Leica Biosystem Inc., Buffalo Grove, IL, USA). Each slide was focused to allow analysis of the maximum number of vessels meeting the criteria for histomorphometric analysis and vessels were annotated for computer assisted data collection (**Supplemental Figure 2**). Pulmonary vessels with external diameter > 300 um, and the bronchial, pleural, vasa vasorum and anastomosing vessels were excluded from the analysis.

Pressure-perfused labelling dilates vessels being labeled and results in lower % wall thickness for the same severity of remodeling. Under the direct supervision of a senior pathologist (WDE), the anatomic landmarks to accurately locate the porcine PA and PV on the histologic slides were learnt from the labelling strategy (in over 10,000 arteries and 10,000 veins) to allow measurement of both labeled and unlabeled vessels. Specifically, in the lung labeled for arteries, unlabeled vessels were considered as veins if their localization in interlobular septa and at the edge of pulmonary acini and the histologic features confirmed venous morphology. In the lung labeled for veins, unlabeled vessels were considered as arteries if their localization in vicinity to pulmonary airways was confirmed.

**LCMD and LC-MS/MS of Pulmonary Vein and Artery Tissue:**

A subset of six PVH and six Sham pigs was assembled for proteomic profiling of PA and PV using LC–MS/MS. For each pig, thick (10 µm) sections of formalin fixed paraffin embedded (FFPE) blocks were cut on to polyethylene naphthalate membrane slides and stained with H&E for contrast. Remodeled (defined as %WT ≥40%) and normal small (ED ≤150 µm) intrapulmonary PA and PV were selectively isolated, through LCMD using Zeiss PALM Microbeam System (Bernried, Germany), from PVH and Sham pigs, respectively. For each sample, approximately 500,000 $\mu m^2$ area of tissue was collected. Each vascular profile was cut along the outer side of adventitia following the removal of debris, such as blood cells or labelling material, from vascular lumen (**Supplemental Figure 3**). Loading was normalized to tissue sample area. Formalin-protein cross-linkages were broken and proteins extracted by heating in a closed Thermomixer at 98 °C for 1 h followed by protein reduction, cooling and alkalization. Proteins were digested with trypsin, centrifuged, transferred to MS vials, concentrated to dryness and stored at -80C until LC-MS/MS analysis. After reconstitution with Pierce stable isotope labeled peptide retention time standards, sample aliquots were loaded onto a 0.25 µl bed OptiPak trap (Optimize Technologies) and washed. Using a Dionex UltiMate 3000 RSLC liquid chromatography system, peptides were transferred onto a PicoFrit column, separated using a 400 nl/min LC gradient and re-equilibrated. Eluting peptides were analyzed using a QExactive Plus mass spectrometer (Thermo-Fisher Scientific). The instrument is configured to operate in data-dependent mode by collecting MS1 data at 70,000 resolving power (measured at m/z 200) with an AGC value of 3E6 over a m/z range of 340–1800, using lock masses from background poly-siloxanes at m/z 371.10123 and 445.12002. Precursors were fragmented with normalized with collision energy of 28, fragments measured at 17,500 resolving power and a fixed first mass of 140. Resulting tandem mass spectra (MS/MS) were collected on the top 20 precursor masses present in each MS1 using an AGC value of 1E5, max ion fill time of 60ms, and an isolation window of 3 Da.

**Whole Lung Tissue Collection and Protein Preparation for LC-MS/MS:**

At tissue harvest, prior to ex-vivo vessel labeling, a section from the mid right posterior lobe was chosen to maximize vessel abundance (periphery) and harvested with a surgical cutting stapler (PROXIMATE® Linear Cutter, size: 75mm), sectioned, weighed, placed in a sample tube and then flash frozen in liquid nitrogen.

Lysis buffer consisted of 0.1% SDS, 20mM Tris, pH 8.2, 1 mM $MgCl_2$, Benzonase and protease inhibitors ProBlock, Simple Stop 1, Simple Stop 3 (GoldBio). Twenty volumes (w/v) of cold lysis buffer was added to each tube. Tissue was lysed on a Minilys bead homogenizer for 30 sec at 5000 rpm x 3. The tubes were heated at 80C for 10 min to denature the proteins, and then spun at 10kxg for 10 min. Supernatants were transferred to 1.5 ml tubes. Protein concentrations were determined using the BCA assay (ThermoFisher).

Eighteen µg of protein from each sample was diluted with SDS-PAGE buffer (Laemmli buffer, 5% beta mercaptoethanol), heated for 10 min at 85°C to reduce disulfide bonds, and loaded on a 12.5% Criterion gel (Bio-Rad) for electrophoresis. The gel was then fixed and stained with BioSafe colloidal blue stain (Bio-Rad). The gel lanes were divided into 6 even sections down the length of the lane horizontally aligned across all sample lanes. Each gel segment (6 per sample) was excised, cut into 1-2mm pieces and transferred to 0.5 ml tubes for subsequent in-gel digest.

Proteins were destained with 40% acetonitrile in 50mM Tris pH 8.1 until clear, reduced with 50 mM TCEP in 50 mM Tris pH 8.1 for 40 minutes at 60°C, followed with alkylation using 25 mM iodoacetamide in 50 mM Tris pH 8.1 for 60 minutes in the dark at room temperature. Proteins were digested in-situ with 0.16 ug trypsin (Promega Corporation, Madison WI) in 25 mM Tris pH 8.1 with 0.0002% Zwittergent 3-16, overnight at 37°C, followed by peptide extraction with 2% trifluoroacetic acid and acetonitrile. Extractions were dried and stored at -20°C.

Dried trypsin digested samples were suspended in sample buffer (0.2% formic acid/0.1% TFA/0.002% zwittergent 3-16) containing 2 fmol/uL Pierce Retention Time Calibration Mixture (Thermo Fisher Scientific, Bremen, Germany). One sixth of the sample was analyzed by nano-flow liquid chromatography electrospray tandem mass spectrometry (nanoLC-ESI-MS/MS) using a Thermo Scientific Q-Exactive Mass Spectrometer (Thermo Fisher Scientific, Bremen, Germany) coupled to a Thermo Ultimate 3000 RSLCnano HPLC system. The digest peptide mixture was loaded onto a 330 nL Halo 2.7 ES-C18 trap (Optimize Technologies, Oregon City, OR). Chromatography was performed using solvent A (98% water/2% acetonitrile/0.2 % formic acid) and solvent B (80% acetonitrile/10%/ isopropanol/10% water/0.2 % formic acid), from 2% to 45% B gradient over 90 minutes at 400 nL/min through a PicoFrit (New Objective, Woburn, MA) 100 µm x 33 cm column hand packed with Agilent Poroshell 120 EC C18 packing. The Q-Exactive mass spectrometer was set to acquire an ms1 survey scans from 350-1600 m/z at resolution 70,000 (at 200 m/z) with an AGC target of 3e6 ions and a maximum ion inject time of 60 msec. Survey scans were followed by HCD MS/MS scans on the top 15 ions at resolution 17,500 with an AGC target of 2e5 ions and a maximum ion inject time of 60 msec. Dynamic exclusion placed selected ions on an exclusion list for 40 seconds.

**Bioinformatics Analysis of LC-MS/MS Data:**

Quality of raw data was evaluated utilizing the NIST MS quality metrics encoded in the Swift proteomic data processing pipeline. MaxQuant software (Max Plank Institute of Biochemistry, Martinsried, Germany), version 1.5.1.2, was configured to use UniProt porcine protein sequence database (downloaded on 02/2019), semitryptic digestion strategy, and the following variable modifications when identifying peptides and proteins present in the samples: carbamidomethylation of cysteine, oxidation of methionine, and n-terminal pyroglutamic acid. Reversed sequences of the proteins were appended to the database for estimating peptide and protein false discovery rates (FDRs). MaxQuant detected proteins present in each sample, grouped them into protein groups based on peptide evidence and reported their intensities using an overall protein group FDR of <= 0.01. An in-house script written in R programming language performed differential expression analysis using protein group intensities. First, protein group intensities of each sample were $\log_2$ transformed and normalized using the quantile method. One PVH and one Sham PV sample showed many fewer proteins and a highly skewed intensity distribution even after normalization suggesting inadequate protein extraction and these samples were excluded from further analysis. The arterial samples from these two pigs did not show a similar issue with distribution and were included in analysis.

For each protein group, the normalized intensities observed in two groups of samples were modeled using a Gaussian-linked generalized linear model. A *t* test was used to detect differentially expressed proteins (DEP) between experimental groups. Differential expression *p* values were false discovery rate (FDR) corrected using the Benjamini–Hochberg–Yekutieli procedure. Protein groups with a FDR p value of <0.05 and absolute $\log_2$ fold change of at least 0.5 (≈1.4 fold increase) were considered as significantly differentially expressed.

**Proteomic Data Analysis using Ingenuity Pathway Analysis Software:**

The log2 fold changes, p-values, FDR p values and gene symbols for the DEP in remodeled PV, PA and WL samples vs their corresponding Sham controls were analyzed in IPA to identify key canonical pathways and PUSR altered in experimental Group 2 PH. In IPA, right-tailed Fischer's Exact test is used to test overlap between DEP in our data sets and datasets given in canonical pathway or upstream regulator and are FDR corrected (Benjamini–Hochberg procedure). The upstream regulator analysis is based on prior knowledge of expected effects between transcriptional regulators and their target genes stored in the Ingenuity® Knowledge Base. The term "upstream regulator" as used in IPA refers to any molecule that can affect the expression of another molecule. Upstream regulators cover a wide variety of molecule types found in the literature such as transcription regulators, receptors, kinases, cytokines, microRNA etc.; including others, drugs and chemicals which were excluded for presentation here. The analysis examines how many known targets of each transcription regulator are present in the dataset, and also compares their direction of change (i.e. expression in the experimental sample(s) relative to control) to what is expected from the literature in order to predict likely relevant transcriptional regulators. If the observed direction of change is mostly consistent with a particular activation state of the transcriptional regulator ("activated" or "inhibited"), then a prediction is made about that activation state.

For key canonical pathways and predicted upstream regulators, an activation *z* score is computed. The activation *z*-score is used to infer likely activation states based on comparison with a model that assigns random regulation directions. The statistical approach defines a quantity (*z*-score) that determines whether a pathway/regulator has significantly more "activated" predictions than "inhibited" predictions ($z>0$) or vice versa ($z<0$). Here, significance means that the hypothesis that predictions are random with equal probability is rejected. If the absolute value of the *z*-score calculated from those numbers is large (i.e. falls into the "tail" of the Gaussian distribution) it would be unlikely to obtain that value of *z* by chance. Moreover, the sign of the calculated *z*-score will reflect the overall predicted activation state of the regulator (<0: inhibited, >0: activated). In practice, *z*-scores greater than 2 or smaller than -2 can be considered statistically significant. A z-score is not calculated if there is insufficient knowledge base on activation status implications of detected pathway proteins in the data set.

For canonical pathway (FDR corrected p value < 0.01) and PUSR analysis, we focus on pathways/regulators meeting rigorous requirements for statistical significance (FDR corrected p value < 0.01 and absolute value of z-score > 2.0) but provide supplemental data on pathways/regulators meeting less rigorous requirements.

**Human Pulmonary Vascular Remodeling**:

From our previous (Mayo IRB approved) study of pulmonary vascular remodeling in autopsy specimens from Control patients and patients with Group 2 PH[1], we selected three Controls and three HF with PH patients for LCMD of PV and PA from FFPE blocks followed by LC-MS/MS and bioinformatics as above. While protein recovery is anticipated to be somewhat impaired given the age of the autopsy FFPE blocks, we sought to provide some information regarding generalizability of the proteomic bioinformatics information provided by the porcine model.

Human HF with PH subjects were selected to have a similar severity of remodeling (% PA and PV wall thickening in unlabeled PA and PV) as observed in the pig model. The entry criteria and histologic methodology, including the methods for discriminating PA from PV were previously described[1]. In human PA, the internal elastic lamina clearly separates intima and media and the % of intima and media thickening (relative to the external diameter of the vessel) are reported. In pigs, the elastic lamina was less consistently distinct and thus total % wall thickening was reported and this convention was followed for the human specimens as well.

Proteomic and Bioinformatics methods were as above for porcine samples except that the UniProt human protein sequence database (downloaded 10/04/2019) was used.

**Supplemental Table 1.** Differentially Expressed Proteins

| UniProtKB Accesion | Entrez Gene Name | Gene Symbol | $Log_2$ Fold Change | $-Log_{10}$ (P-value) | $-Log_{10}$ (FDR P-value) |
|---|---|---|---|---|---|
| **PVH Pulmonary Artery vs. Sham Pulmonary Artery** | | | | | |
| I3LUM8 | phenylalanyl-tRNA synthetase subunit beta | FARSB | 3.5 | 23.24 | 19.88 |
| P62863 | FAU ubiquitin like and ribosomal protein S30 fusion | FAU | 1.6 | 15.47 | 12.42 |
| I3LSA5 | Alpha-amylase | AMY1A | -30.2 | 13.64 | 10.77 |
| F1SPH1 | proliferation-associated 2G4 | PA2G4 | 1.2 | 10.98 | 8.23 |
| Q6EEI7 | mannose receptor C-type 1 | MRC1 | 6.7 | 10.18 | 7.53 |
| A1YH85 | MHC class I antigen | HLA-2 | -2.1 | 8.84 | 6.27 |
| A0A287AVQ1 | DEAD-box helicase 3 X-linked | DDX3X | 1.3 | 8.68 | 6.23 |
| F2Z5G8 | ribosomal protein S25 | RPS25 | 1.2 | 8.70 | 6.23 |
| F1RTH3 | apoptosis inducing factor mitochondria associated 1 | AIFM1 | 2.6 | 8.04 | 5.65 |
| P67985 | ribosomal protein L22 | RPL22 | 1.2 | 7.78 | 5.43 |
| A0A287B7J9 | ribosomal protein S24 | RPS24 | 1.9 | 7.58 | 5.28 |
| F2Z5F5 | ribosomal protein S8 | RPS8 | 1.8 | 7.55 | 5.28 |
| A0A287AWS4 | ribosomal protein L27a | RPL27A | 1.4 | 7.28 | 5.05 |
| F1RQI0 | collagen type XII alpha 1 chain | COL12A1 | 2.6 | 7.22 | 5.01 |
| Q29205 | ribosomal protein L11 | RPL11 | 1.1 | 7.05 | 4.88 |
| Q2YGT9 | ribosomal protein L6 | RPL6 | 1.6 | 6.89 | 4.74 |
| F1SMS8 | lectin, mannose binding 1 | LMAN1 | 3.2 | 6.57 | 4.44 |
| I3LEE6 | procollagen C-endopeptidase enhancer | PCOLCE | 27.6 | 6.52 | 4.42 |
| A0A287ACK3 | DAB adaptor protein 2 | DAB2 | 25.7 | 6.40 | 4.37 |
| F1S935 | ribosomal protein L18a | RPL18A | 3.6 | 6.38 | 4.37 |
| I3LJ87 | ribosomal protein S2 | RPS2 | 1.6 | 6.40 | 4.37 |
| A0A287AKC5 | ribosomal protein L38 | RPL38 | 1.5 | 6.40 | 4.37 |
| P18137 | lactalbumin alpha | LALBA | -27.9 | 6.26 | 4.27 |
| A0A287A2G6 | glutathione peroxidase 7 | GPX7 | 27.7 | 6.14 | 4.16 |
| F2Z5K2 | proteasome 20S subunit alpha 5 | PSMA5 | -1.3 | 6.12 | 4.16 |
| A0A287BB44 | multimerin 2 | MMRN2 | -29.5 | 6.09 | 4.16 |
| H2F098 | macrophage receptor with collagenous structure | MARCO | -27.6 | 6.00 | 4.08 |
| A0A287BRY6 | SPARC like 1 | SPARCL1 | 27.0 | 5.84 | 3.94 |
| I3L6R1 | cartilage associated protein | CRTAP | 20.9 | 5.80 | 3.92 |
| I3LP78 | ribosomal protein L9 | RPL9 | 1.5 | 5.76 | 3.88 |
| A0A286ZRU9 | serpin family H member 1 | SERPINH1 | 2.5 | 5.66 | 3.80 |
| F1SAE9 | laminin subunit beta 1 | LAMB1 | 2.4 | 5.60 | 3.77 |
| P49666 | ribosomal protein L21 | RPL21 | 1.3 | 5.61 | 3.77 |
| F2Q9A3 | peptidylprolyl isomerase A | PPIA | 1.0 | 5.51 | 3.69 |
| F2Z512 | ribosomal protein S23 | RPS23 | 2.1 | 5.42 | 3.62 |
| A0A287AR45 | stabilin 1 | STAB1 | 23.4 | 5.38 | 3.59 |
| B1Q039 | crystallin alpha B | CRYAB | 2.4 | 5.34 | 3.56 |
| A0A286ZXA0 | insulin like growth factor binding protein 7 | IGFBP7 | 4.9 | 5.25 | 3.48 |
| A0A287A8T0 | ribosomal protein L7 | RPL7 | 1.7 | 5.17 | 3.43 |
| G9F6X8 | prolyl 4-hydroxylase subunit beta | P4HB | 1.6 | 5.18 | 3.43 |

| | | | | | |
|---|---|---|---|---|---|
| H6TBN0 | thioredoxin | TXN | -1.6 | 5.18 | 3.43 |
| I3LTQ6 | sideroflexin 3 | SFXN3 | 2.5 | 5.14 | 3.43 |
| A0A287AE76 | ribosomal protein L4 | RPL4 | 1.7 | 5.14 | 3.43 |
| F1RTR7 | YOD1 deubiquitinase | YOD1 | -25.5 | 4.92 | 3.22 |
| I3LKU0 | Rac family small GTPase 2 | RAC2 | -2.4 | 4.90 | 3.21 |
| M9TGS8 | enoyl-CoA hydratase 1 | ECH1 | 23.4 | 4.87 | 3.20 |
| A0A287BM53 | ribosomal protein L7a | RPL7A | 1.3 | 4.85 | 3.18 |
| F1S7U3 | chitinase 3 like 1 | CHI3L1 | 7.4 | 4.82 | 3.16 |
| K7GNA9 | platelet and endothelial cell adhesion molecule 1 | PECAM1 | -2.3 | 4.76 | 3.11 |
| F2Z567 | ribosomal protein L8 | RPL8 | 1.3 | 4.72 | 3.07 |
| A0A286ZQ40 | ribosomal protein S26 | RPS26 | 1.3 | 4.70 | 3.06 |
| F1SUQ3 | ER membrane protein complex subunit 1 | EMC1 | 19.0 | 4.66 | 3.03 |
| A0A287AJT7 | ribosomal protein L28 | RPL28 | 1.8 | 4.63 | 3.01 |
| K7GKC0 | ribosomal protein S16 | RPS16 | 1.3 | 4.56 | 2.95 |
| A0A287B356 | eukaryotic translation initiation factor 5A | EIF5A | 0.8 | 4.55 | 2.94 |
| F1SUU4 | filamin binding LIM protein 1 | FBLIM1 | 4.3 | 4.53 | 2.93 |
| A0A287AP66 | ribosomal protein L12 | RPL12 | 1.1 | 4.51 | 2.92 |
| A0A286ZUW0 | glutamine--fructose-6-phosphate transaminase 1 | GFPT1 | 4.6 | 4.43 | 2.85 |
| I3LV17 | RAS like proto-oncogene B | RALB | -2.4 | 4.39 | 2.82 |
| A0A287AYJ0 | dihydropyrimidinase like 3 | DPYSL3 | 1.6 | 4.31 | 2.75 |
| K7GLN4 | peroxiredoxin 4 | PRDX4 | 1.0 | 4.31 | 2.75 |
| A0A286ZQ63 | SEC14 like lipid binding 3 | SEC14L3 | -26.1 | 4.28 | 2.73 |
| I3LFP3 | versican | VCAN | 3.4 | 4.23 | 2.68 |
| A0A287AEG8 | ribosomal protein L10 | RPL10 | 1.2 | 4.08 | 2.54 |
| A5D9J3 | transporter 1, ATP binding cassette subfamily B member | TAP1 | -3.2 | 4.06 | 2.53 |
| F2Z4Y8 | ribosomal protein S11 | RPS11 | 1.7 | 3.95 | 2.42 |
| F1SGM3 | proteasome activator subunit 2 | PSME2 | -1.8 | 3.93 | 2.41 |
| A0PA01 | Serine protease inhibitor 9 | PI-9 | -1.5 | 3.91 | 2.40 |
| P79385 | milk fat globule EGF and factor V/VIII domain containing | MFGE8 | 26.2 | 3.91 | 2.40 |
| F1S4D7 | guanylate binding protein 1 | GBP1 | -3.0 | 3.85 | 2.35 |
| A0A286ZLH8 | ribosomal protein L35a | RPL35A | 2.1 | 3.83 | 2.34 |
| F1RZ28 | ribosomal protein S10 | RPS10 | 1.3 | 3.83 | 2.34 |
| B8XZY6 | high density lipoprotein binding protein | HDLBP | 1.7 | 3.82 | 2.34 |
| F1RJP9 | kinesin family member 13B | KIF13B | 19.7 | 3.78 | 2.30 |
| F2Z557 | poly(A) binding protein cytoplasmic 1 | PABPC1 | 1.1 | 3.74 | 2.27 |
| A0A287AK65 | argininosuccinate synthase 1 | ASS1 | -19.6 | 3.67 | 2.21 |
| I3LS12 | sorting nexin 2 | SNX2 | -2.4 | 3.60 | 2.14 |
| A0A286ZMW0 | endoplasmic reticulum protein 44 | ERP44 | 1.3 | 3.59 | 2.13 |
| A0A287BGN7 | ribosomal protein S6 | RPS6 | 2.1 | 3.58 | 2.13 |
| A0A286ZUZ5 | microtubule associated protein 1S | MAP1S | -1.2 | 3.51 | 2.07 |
| A0A287AN50 | agrin | AGRN | 1.2 | 3.49 | 2.06 |
| E1CAJ6 | Protein disulfide isomerase P5 | PDI-P5 | 1.2 | 3.47 | 2.04 |
| I3LRZ4 | PPFIA binding protein 1 | PPFIBP1 | 22.4 | 3.46 | 2.03 |
| I3LSD3 | ribosomal protein L13 | RPL13 | 1.0 | 3.45 | 2.02 |
| A0A287AH85 | adenylate kinase 2 | AK2 | 1.3 | 3.44 | 2.02 |
| F2Z4Z8 | G protein subunit beta 2 | GNB2 | -1.1 | 3.43 | 2.02 |

| | | | | | |
|---|---|---|---|---|---|
| I3LT81 | RPL17-C18orf32 readthrough | RPL17-C18orf32 | 1.5 | 3.41 | 2.00 |
| A0A287AEU6 | methionine adenosyltransferase 2B | MAT2B | 1.3 | 3.36 | 1.96 |
| A0A287AGN9 | spondin 1 | SPON1 | 20.2 | 3.32 | 1.93 |
| A0A287AMZ2 | chaperonin containing TCP1 subunit 3 | CCT3 | 0.9 | 3.32 | 1.93 |
| I3LP35 | Peptidase S1 domain-containing protein | LOC100154047 | -20.8 | 3.28 | 1.89 |
| A0A286ZQS7 | sec1 family domain containing 1 | SCFD1 | 21.0 | 3.24 | 1.87 |
| P62831 | ribosomal protein L23 | RPL23 | 2.8 | 3.23 | 1.87 |
| F2Z505 | eukaryotic translation termination factor 1 | ETF1 | 1.4 | 3.24 | 1.87 |
| F1SEN2 | glutamate dehydrogenase 1 | GLUD1 | 1.0 | 3.23 | 1.87 |
| Q56P20 | ADP ribosylation factor 4 | ARF4 | 0.9 | 3.23 | 1.87 |
| A0A287ADH9 | chloride intracellular channel 4 | CLIC4 | 0.8 | 3.23 | 1.87 |
| F1RQQ7 | glycogen phosphorylase B | PYGB | 0.6 | 3.23 | 1.87 |
| I3LD72 | EH domain containing 2 | EHD2 | -1.1 | 3.24 | 1.87 |
| F1RYA3 | Cysteinyl-tRNA synthetase | CARS | 20.9 | 3.17 | 1.82 |
| A0A286ZRM3 | FLII actin remodeling protein | FLII | 1.9 | 3.15 | 1.81 |
| A0A286ZJQ9 | caveolin 1 | CAV1 | -2.0 | 3.15 | 1.81 |
| F1SU03 | heparan sulfate proteoglycan 2 | HSPG2 | 1.2 | 3.14 | 1.81 |
| Q29361 | ribosomal protein L35 | RPL35 | 2.7 | 3.09 | 1.77 |
| Q52NJ3 | secretion associated Ras related GTPase 1A | SAR1A | 1.3 | 3.09 | 1.77 |
| F8TEL6 | serine and arginine rich splicing factor 10 | SRSF10 | 0.7 | 3.10 | 1.77 |
| F1S431 | Alanine--tRNA ligase | AARS | 2.9 | 3.08 | 1.77 |
| I3LNY6 | nestin | NES | 2.1 | 3.08 | 1.77 |
| Q29092 | heat shock protein 90 beta family member 1 | HSP90B1 | 1.5 | 3.08 | 1.77 |
| F1RF11 | matrix metallopeptidase 2 | MMP2 | 20.8 | 3.03 | 1.73 |
| Q5S1U1 | heat shock protein family B (small) member 1 | HSPB1 | 0.7 | 3.01 | 1.72 |
| A0A287B5V1 | ribosomal protein L3 | RPL3 | 1.3 | 2.99 | 1.69 |
| I3LEF8 | hydroxysteroid 17-beta dehydrogenase 4 | HSD17B4 | 2.0 | 2.98 | 1.69 |
| F1SFF3 | nidogen 2 | NID2 | 1.3 | 2.98 | 1.69 |
| B2NJ26 | beta-2-microglobulin | B2M | -2.6 | 2.96 | 1.67 |
| P35750 | calpain 1 | CAPN1 | -0.8 | 2.95 | 1.67 |
| F1SC47 | aldehyde dehydrogenase 18 family member A1 | ALDH18A1 | 19.6 | 2.93 | 1.65 |
| A0A287AZ68 | scribble planar cell polarity protein | SCRIB | -18.6 | 2.92 | 1.65 |
| B6EAV5 | SLC9A3 regulator 2 | SLC9A3R2 | -1.7 | 2.91 | 1.65 |
| K7GS03 | LDL receptor related protein 1 | LRP1 | 1.1 | 2.90 | 1.63 |
| F1RYZ0 | ribosomal protein lateral stalk subunit P2 | RPLP2 | 0.9 | 2.87 | 1.61 |
| I3LNH3 | glucosidase II alpha subunit | GANAB | 1.0 | 2.83 | 1.57 |
| A0A287AXD1 | myocardial zonula adherens protein | MYZAP | -15.0 | 2.83 | 1.57 |
| Q6QAT1 | ribosomal protein S28 | RPS28 | 0.9 | 2.82 | 1.57 |
| Q6JLA8 | four and a half LIM domains 3 | FHL3 | 22.1 | 2.78 | 1.53 |
| A0A287AN95 | PDZ and LIM domain 5 | PDLIM5 | 1.2 | 2.78 | 1.53 |
| A0A287BJ88 | basic transcription factor 3 | BTF3 | 20.9 | 2.77 | 1.53 |
| A0A287B771 | ribosomal protein S4 X-linked | RPS4X | 1.3 | 2.76 | 1.52 |
| F1RGJ2 | catenin alpha 1 | CTNNA1 | -1.2 | 2.75 | 1.52 |
| A0A287B600 | synemin | SYNM | -1.2 | 2.75 | 1.52 |
| F1RLC4 | lysyl oxidase | LOX | 17.9 | 2.74 | 1.51 |
| A0A287A2P1 | ciliary rootlet coiled-coil, rootletin family member 2 | CROCC2 | -21.2 | 2.73 | 1.50 |

| | | | | | |
|---|---|---|---|---|---|
| F2Z546 | ribosomal protein L19 | RPL19 | 1.0 | 2.72 | 1.50 |
| Q95342 | ribosomal protein L18 | RPL18 | 1.7 | 2.72 | 1.50 |
| I3LAB6 | proteasome 20S subunit alpha 2 | PSMA2 | -1.2 | 2.71 | 1.50 |
| I3LDR9 | caveolae associated protein 2 | CAVIN2 | -2.1 | 2.71 | 1.50 |
| F1S6S9 | proteinase 3 | PRTN3 | -22.5 | 2.70 | 1.49 |
| F1SJ86 | chondroitin sulfate proteoglycan 4 | CSPG4 | 1.7 | 2.69 | 1.49 |
| B9TRW9 | G protein subunit gamma 12 | GNG12 | -1.3 | 2.69 | 1.49 |
| A0A286ZL65 | ribosomal protein L15 | RPL15 | 1.3 | 2.67 | 1.47 |
| A0A286ZIL9 | collagen type XVIII alpha 1 chain | COL18A1 | 1.7 | 2.64 | 1.45 |
| A0A287AEH1 | laminin subunit alpha 5 | LAMA5 | 1.3 | 2.64 | 1.45 |
| G8FUN5 | Y-box binding protein 1 | YBX1 | 1.1 | 2.64 | 1.45 |
| F1RUX1 | coronin 1B | CORO1B | -0.9 | 2.63 | 1.44 |
| A0A286ZW70 | peptidylprolyl isomerase B | PPIB | 1.4 | 2.62 | 1.43 |
| F2Z554 | ribosomal protein L30 | RPL30 | 1.0 | 2.60 | 1.42 |
| F1SB57 | armadillo repeat containing 10 | ARMC10 | 0.6 | 2.60 | 1.42 |
| A0A287A3C9 | Inositol 1,4,5-triphosphate receptor associated 1 | MRVI1 | 20.0 | 2.57 | 1.40 |
| F1RJX8 | COPI coat complex subunit alpha | COPA | 1.3 | 2.57 | 1.40 |
| I3LQ79 | major vault protein | MVP | 1.4 | 2.50 | 1.34 |
| A0A286ZN31 | Sad1 and UNC84 domain containing 2 | SUN2 | -0.8 | 2.51 | 1.34 |
| F1SJE6 | phosphoglucomutase 5 | PGM5 | -1.7 | 2.49 | 1.33 |
| F1SJQ6 | ribosomal protein L14 | RPL14 | 2.1 | 2.49 | 1.33 |
| K7GPF5 | CD248 molecule | CD248 | 15.2 | 2.48 | 1.32 |
| A0A287BHE7 | ribosomal protein S14 | RPS14 | 1.0 | 2.48 | 1.32 |
| A0A287B9Y4 | UPF1 RNA helicase and ATPase | UPF1 | 3.9 | 2.47 | 1.31 |
| A0A287A1A0 | ankyrin repeat and FYVE domain containing 1 | ANKFY1 | -16.9 | 2.46 | 1.31 |
| F1SJY1 | endoplasmic reticulum-golgi intermediate compartment 1 | ERGIC1 | 17.5 | 2.45 | 1.31 |
| A0A287A8P0 | OS9 endoplasmic reticulum lectin | OS9 | 16.1 | 2.45 | 1.31 |
| F1S0M9 | FKBP prolyl isomerase 10 | FKBP10 | 19.8 | 2.45 | 1.30 |
| F1RX00 | monoamine oxidase A | MAOA | -0.7 | 2.44 | 1.30 |
| **PVH Pulmonary Vein vs. Sham Pulmonary Vein** | | | | | |
| P79385 | milk fat globule EGF and factor V/VIII domain containing | MFGE8 | 35.6 | 270.45 | 269.40 |
| A0A287A2G6 | glutathione peroxidase 7 | GPX7 | 33.3 | 270.45 | 267.40 |
| F1RNR2 | SET domain containing 1B, histone lysine methyltransferase | SETD1B | 31.2 | 100.72 | 97.85 |
| F1SJ07 | pleckstrin | PLEK | -32.0 | 99.90 | 97.15 |
| A1X898 | prolyl 4-hydroxylase subunit alpha 1 | P4HA1 | 23.6 | 97.37 | 94.72 |
| F1RKK5 | proline rich coiled-coil 1 | PRRC1 | 24.8 | 81.70 | 79.13 |
| A0A287AEU6 | methionine adenosyltransferase 2B | MAT2B | 1.9 | 30.73 | 28.28 |
| A0A287BJG3 | 3'-phosphoadenosine 5'-phosphosulfate synthase 1 | PAPSS1 | 27.7 | 29.15 | 26.76 |
| B4YYD8 | nitric oxide synthase 3 | NOS3 | -30.8 | 21.40 | 19.05 |
| A0A287BFZ2 | erythrocyte membrane protein band 4.1 like 2 | EPB41L2 | -1.9 | 17.99 | 15.68 |
| F1SJ86 | chondroitin sulfate proteoglycan 4 | CSPG4 | 3.0 | 15.71 | 13.44 |
| A0A287BR41 | protein kinase C and casein kinase substrate in neurons 2 | PACSIN2 | 29.2 | 14.41 | 12.17 |
| H2F098 | macrophage receptor with collagenous structure | MARCO | -31.0 | 13.13 | 10.92 |
| A0A286ZIL9 | collagen type XVIII alpha 1 chain | COL18A1 | 3.3 | 11.57 | 9.40 |

| | | | | | |
|---|---|---|---|---|---|
| A0A287AKW4 | glycoprotein nmb | GPNMB | 2.5 | 11.29 | 9.17 |
| A1YH85 | MHC class I antigen | HLA-2 | -2.9 | 11.29 | 9.17 |
| F1SFI5 | histidine rich glycoprotein | HRG | -3.7 | 11.08 | 8.98 |
| F1RFT3 | proline, glutamate and leucine rich protein 1 | PELP1 | 21.7 | 10.78 | 8.71 |
| A0A287BL58 | PBX homeobox interacting protein 1 | PBXIP1 | 5.9 | 10.57 | 8.53 |
| A0A286ZUW0 | glutamine--fructose-6-phosphate transaminase 1 | GFPT1 | 4.5 | 9.82 | 7.80 |
| A0A287B2U6 | synaptosome associated protein 23 | SNAP23 | -3.5 | 9.74 | 7.73 |
| Q1ACV4 | transporter 2, ATP binding cassette subfamily B member | TAP2 | -4.1 | 9.63 | 7.64 |
| F1S8Z3 | von Willebrand factor A domain containing 5A | VWA5A | 20.7 | 8.68 | 6.72 |
| A0A287BBE9 | TBC1 domain family member 9B | TBC1D9B | 19.5 | 8.26 | 6.31 |
| A0A286ZFW3 | apolipoprotein H | APOH | -4.9 | 8.08 | 6.14 |
| I3LKF3 | fascin actin-bundling protein 1 | FSCN1 | 1.3 | 7.88 | 5.96 |
| A0A287B356 | eukaryotic translation initiation factor 5A | EIF5A | 1.0 | 7.82 | 5.92 |
| A0A287B792 | UDP-N-acetylglucosamine pyrophosphorylase 1 | UAP1 | 13.4 | 7.66 | 5.79 |
| A5GFX6 | tubulin beta 1 class VI | TUBB1 | -29.0 | 7.66 | 5.79 |
| F1RQI0 | collagen type XII alpha 1 chain | COL12A1 | 4.1 | 7.63 | 5.78 |
| A0A286ZZB2 | syntaxin binding protein 1 | STXBP1 | -8.2 | 7.61 | 5.77 |
| A0A286ZNQ7 | protein kinase cAMP-dependent type II regulatory subunit beta | PRKAR2B | -24.4 | 7.61 | 5.77 |
| F2Z5R5 | programmed cell death 10 | PDCD10 | -28.1 | 7.55 | 5.73 |
| F1SHL9 | pyruvate kinase M1/2 | PKM | 1.0 | 7.34 | 5.54 |
| I3LRS5 | aldehyde dehydrogenase 1 family member A1 | ALDH1A1 | 2.8 | 7.22 | 5.43 |
| A0A287ASF0 | SH3 and multiple ankyrin repeat domains 3 | SHANK3 | -25.3 | 7.20 | 5.41 |
| A0A287AXD1 | myocardial zonula adherens protein | MYZAP | -20.9 | 7.18 | 5.41 |
| F1S7U3 | chitinase 3 like 1 | CHI3L1 | 8.2 | 7.05 | 5.29 |
| B9TRW9 | G protein subunit gamma 12 | GNG12 | -1.2 | 7.00 | 5.25 |
| A0A287ARI1 | eukaryotic translation initiation factor 3 subunit F | EIF3F | 1.6 | 6.95 | 5.22 |
| A0A287A0A6 | collagen type VI alpha 6 chain | COL6A6 | -3.2 | 6.95 | 5.22 |
| A0A287AHT5 | small nuclear ribonucleoprotein D1 polypeptide | SNRPD1 | 1.0 | 6.87 | 5.15 |
| Q8WMN8 | lactotransferrin | LTF | -3.0 | 6.83 | 5.12 |
| B6EAV5 | SLC9A3 regulator 2 | SLC9A3R2 | -3.6 | 6.81 | 5.12 |
| A0A287BQW3 | Histone H1.0 | H1F0 | -1.2 | 6.80 | 5.12 |
| A0A287A853 | G protein subunit alpha 13 | GNA13 | -3.9 | 6.79 | 5.11 |
| Q2EN76 | NME/NM23 nucleoside diphosphate kinase 2 | NME2 | 1.3 | 6.78 | 5.11 |
| I3L816 | heterogeneous nuclear ribonucleoprotein H1 | HNRNPH1 | 0.7 | 6.63 | 4.98 |
| I3LDR9 | caveolae associated protein 2 | CAVIN2 | -2.1 | 6.63 | 4.98 |
| C4MXZ1 | ADP ribosylation factor 1 | ARF1 | 0.7 | 6.48 | 4.84 |
| A0A287ARZ1 | thioredoxin domain containing 5 | TXNDC5 | 2.5 | 6.41 | 4.78 |
| F1RRD6 | programmed cell death 6 interacting protein | PDCD6IP | -3.7 | 6.39 | 4.76 |
| A0A287B1J4 | spectrin alpha, non-erythrocytic 1 | SPTAN1 | -1.3 | 6.01 | 4.40 |
| F1SDX6 | transglutaminase 2 | TGM2 | 1.2 | 5.91 | 4.31 |
| I3LKU0 | Rac family small GTPase 2 | RAC2 | -5.1 | 5.90 | 4.31 |
| A0A287BJ70 | coagulation factor V | F5 | -28.4 | 5.91 | 4.31 |
| A0A287BT04 | prolyl 3-hydroxylase 1 | P3H1 | 22.4 | 5.83 | 4.24 |
| I3LNY6 | nestin | NES | 3.7 | 5.80 | 4.22 |
| A0A287AB52 | microfibril associated protein 4 | MFAP4 | -2.5 | 5.63 | 4.06 |

| | | | | | |
|---|---|---|---|---|---|
| F1RRW5 | angiotensin I converting enzyme | ACE | -5.0 | 5.48 | 3.92 |
| P80015 | azurocidin 1 | AZU1 | -4.6 | 5.23 | 3.68 |
| I3LLD5 | eukaryotic translation initiation factor 4A1 | EIF4A1 | 2.5 | 5.22 | 3.67 |
| F1SFI7 | alpha 2-HS glycoprotein | AHSG | 5.5 | 5.19 | 3.65 |
| F1S5D8 | protein tyrosine phosphatase receptor type C | PTPRC | -2.9 | 5.18 | 3.64 |
| P23687 | prolyl endopeptidase | PREP | 2.6 | 5.13 | 3.60 |
| A0A287AH93 | extended synaptotagmin 1 | ESYT1 | 1.0 | 5.05 | 3.53 |
| F1SJT7 | apolipoprotein A4 | APOA4 | -29.5 | 5.02 | 3.50 |
| A0A286ZNI3 | caveolae associated protein 1 | CAVIN1 | -1.2 | 4.99 | 3.48 |
| P22411 | dipeptidyl peptidase 4 | DPP4 | -26.2 | 4.88 | 3.38 |
| A0A287B5D0 | UBX domain protein 6 | UBXN6 | -22.6 | 4.83 | 3.33 |
| F1RJK5 | GCN1 activator of EIF2AK4 | GCN1 | 4.0 | 4.81 | 3.32 |
| A0A287AAG0 | UDP-N-acetylglucosamine pyrophosphorylase 1 like 1 | UAP1L1 | 22.2 | 4.80 | 3.32 |
| I3LQ79 | major vault protein | MVP | 1.2 | 4.78 | 3.31 |
| A0A287BEK4 | NADH:ubiquinone oxidoreductase subunit V3 | NDUFV3 | 1.0 | 4.79 | 3.31 |
| P83686 | cytochrome b5 reductase 3 | CYB5R3 | -1.0 | 4.77 | 3.30 |
| A0A287AR67 | sorcin | SRI | -1.0 | 4.76 | 3.30 |
| A0A287AT38 | AKAP2_C domain-containing protein | AKAP2 | -1.7 | 4.75 | 3.30 |
| A0A287AG48 | keratin 7 | KRT7 | 4.0 | 4.74 | 3.28 |
| Q56P20 | ADP ribosylation factor 4 | ARF4 | 1.5 | 4.70 | 3.25 |
| A0A286ZRU9 | serpin family H member 1 | SERPINH1 | 2.1 | 4.69 | 3.24 |
| F1RGY5 | nidogen 1 | NID1 | -1.5 | 4.66 | 3.23 |
| A0A287ACQ4 | catenin beta 1 | CTNNB1 | -1.2 | 4.61 | 3.18 |
| A0A287B0Q6 | dynamin 1 like | DNM1L | 7.8 | 4.60 | 3.18 |
| F1RKJ9 | isochorismatase domain containing 1 | ISOC1 | 4.1 | 4.58 | 3.17 |
| F1SAE9 | laminin subunit beta 1 | LAMB1 | 2.0 | 4.55 | 3.14 |
| K7GM40 | apolipoprotein A1 | APOA1 | -2.4 | 4.51 | 3.10 |
| F1RRK9 | exportin 5 | XPO5 | 2.3 | 4.48 | 3.08 |
| E1CAJ6 | Protein disulfide isomerase P5 | PDI-P5 | 1.5 | 4.42 | 3.02 |
| A0A287AJU1 | coactosin like F-actin binding protein 1 | COTL1 | 1.6 | 4.36 | 2.97 |
| P00339 | lactate dehydrogenase A | LDHA | 0.9 | 4.32 | 2.94 |
| B0FWK5 | ribosomal protein L5 | RPL5 | 0.9 | 4.32 | 2.94 |
| I3LK59 | enolase 1 | ENO1 | 1.0 | 4.29 | 2.91 |
| A0A287A0G6 | MMS19 homolog, cytosolic iron-sulfur assembly component | MMS19 | 24.3 | 4.24 | 2.87 |
| A0A287AN95 | PDZ and LIM domain 5 | PDLIM5 | 1.5 | 4.21 | 2.85 |
| K7GNA9 | platelet and endothelial cell adhesion molecule 1 | PECAM1 | -1.8 | 4.18 | 2.82 |
| A0A286ZRG3 | PRAME family member 27-like | LOC100737973 | 28.8 | 4.13 | 2.78 |
| A3EX84 | galectin 3 | LGALS3 | 2.2 | 4.13 | 2.78 |
| A0A287BGV1 | latent transforming growth factor beta binding protein 4 | LTBP4 | -2.6 | 4.13 | 2.78 |
| A0A287BQG2 | canopy FGF signaling regulator 4 | CNPY4 | 24.4 | 4.12 | 2.78 |
| A0A287B2P1 | chloride intracellular channel 1 | CLIC1 | 0.9 | 4.12 | 2.78 |
| A0A287AGU1 | integrin subunit beta 2 | ITGB2 | -2.7 | 4.07 | 2.74 |
| F1S1C3 | carbonic anhydrase 4 | CA4 | -28.7 | 4.07 | 2.74 |
| I3LMH4 | eukaryotic translation initiation factor 4 gamma 1 | EIF4G1 | 1.5 | 4.06 | 2.73 |
| A0A287AU48 | CD34 molecule | CD34 | -4.9 | 4.04 | 2.72 |

| | | | | | |
|---|---|---|---|---|---|
| P79293 | myosin heavy chain 7 | MYH7 | -25.8 | 4.04 | 2.72 |
| K7GLN4 | peroxiredoxin 4 | PRDX4 | 0.9 | 4.02 | 2.71 |
| I3LRZ4 | PPFIA binding protein 1 | PPFIBP1 | 23.8 | 4.00 | 2.69 |
| A0A287AM26 | arginyl aminopeptidase | RNPEP | 2.0 | 3.99 | 2.68 |
| I3LA63 | Methionine--tRNA ligase, cytoplasmic | MARS | 23.2 | 3.97 | 2.67 |
| A0A286ZRI5 | galectin 1 | LGALS1 | 2.0 | 3.92 | 2.62 |
| F1S1M8 | diaphanous related formin 2 | DIAPH2 | 14.3 | 3.89 | 2.59 |
| F1RY68 | carnitine palmitoyltransferase 1A | CPT1A | 24.2 | 3.87 | 2.58 |
| K7GNZ3 | nascent polypeptide associated complex subunit alpha | NACA | 1.4 | 3.87 | 2.58 |
| K9IWG6 | spectrin beta, non-erythrocytic 1 | SPTBN1 | -1.0 | 3.87 | 2.58 |
| F1RJX8 | COPI coat complex subunit alpha | COPA | 1.9 | 3.86 | 2.57 |
| F2Q9A3 | peptidylprolyl isomerase A | PPIA | 1.0 | 3.84 | 2.56 |
| F1RWT2 | plastin 3 | PLS3 | 0.9 | 3.82 | 2.55 |
| F1SIX3 | UDP-glucose pyrophosphorylase 2 | UGP2 | 3.8 | 3.80 | 2.53 |
| L7PBE6 | chaperonin containing TCP1 subunit 5 | CCT5 | 1.3 | 3.80 | 2.53 |
| F2Z512 | ribosomal protein S23 | RPS23 | 1.4 | 3.79 | 2.53 |
| Q9GMA6 | serpin family A member 3 | SERPINA3 | -22.5 | 3.77 | 2.51 |
| M3TYW5 | Glutaminyl-tRNA synthetase | QARS | 2.0 | 3.76 | 2.51 |
| A0A287BQQ3 | adducin 3 | ADD3 | -1.1 | 3.76 | 2.51 |
| F1SFF8 | glycogen phosphorylase L | PYGL | -1.3 | 3.74 | 2.49 |
| F1RJ93 | transgelin 2 | TAGLN2 | 0.9 | 3.73 | 2.48 |
| A0A286ZXH3 | NECAP endocytosis associated 2 | NECAP2 | 23.0 | 3.72 | 2.48 |
| F1SV89 | histone deacetylase 1 | HDAC1 | -1.6 | 3.71 | 2.48 |
| A0A287BKM2 | xin actin binding repeat containing 2 | XIRP2 | 26.5 | 3.66 | 2.42 |
| A0A286ZJQ9 | caveolin 1 | CAV1 | -2.2 | 3.65 | 2.42 |
| F1RUK8 | GDP dissociation inhibitor 2 | GDI2 | 0.7 | 3.63 | 2.41 |
| B2NJ26 | beta-2-microglobulin | B2M | -2.2 | 3.62 | 2.40 |
| G1FJ20 | ATP citrate lyase | ACLY | 0.9 | 3.61 | 2.40 |
| F1SPF9 | COPI coat complex subunit gamma 1 | COPG1 | 1.6 | 3.60 | 2.39 |
| F1RGG1 | ribosomal protein S19 | RPS19 | 1.3 | 3.60 | 2.39 |
| F1S3W0 | ubiquinol-cytochrome c reductase hinge protein | UQCRH | -3.1 | 3.58 | 2.37 |
| F1SUP2 | signal peptidase complex subunit 2 | SPCS2 | 23.9 | 3.58 | 2.37 |
| I3LDC7 | isocitrate dehydrogenase (NADP(+)) 1 | IDH1 | 1.0 | 3.56 | 2.36 |
| A0A287AWI9 | eukaryotic translation elongation factor 2 | EEF2 | 0.9 | 3.56 | 2.36 |
| K9IVP5 | myosin heavy chain 9 | MYH9 | 0.7 | 3.54 | 2.35 |
| Q08094 | calponin 2 | CNN2 | 0.6 | 3.55 | 2.35 |
| A0A287B278 | annexin A4 | ANXA4 | 1.0 | 3.50 | 2.31 |
| F1RU28 | DR1 associated protein 1 | DRAP1 | 2.5 | 3.47 | 2.29 |
| F1SJS8 | transgelin | TAGLN | 1.9 | 3.45 | 2.26 |
| M3V7Z1 | cytoplasmic FMR1 interacting protein 1 | CYFIP1 | -0.7 | 3.41 | 2.23 |
| F1SPS7 | acylaminoacyl-peptide hydrolase | APEH | -4.3 | 3.41 | 2.23 |
| A0A286ZW70 | peptidylprolyl isomerase B | PPIB | 1.4 | 3.40 | 2.23 |
| A0A287ANL7 | erythrocyte membrane protein band 4.1 | EPB41 | -26.0 | 3.40 | 2.23 |
| F1SK03 | alanyl aminopeptidase, membrane | ANPEP | 19.3 | 3.39 | 2.23 |
| A0A287BMG2 | trans-2,3-enoyl-CoA reductase | TECR | 6.3 | 3.38 | 2.22 |
| A0A286ZI90 | immunoglobulin heavy constant mu | IGHM | -2.1 | 3.38 | 2.22 |

| | | | | | |
|---|---|---|---|---|---|
| A0A287BNU5 | RAB5B, member RAS oncogene family | RAB5B | -3.6 | 3.39 | 2.22 |
| A0A287B341 | nudix hydrolase 16 | NUDT16 | 10.4 | 3.32 | 2.17 |
| A0A287AVQ1 | DEAD-box helicase 3 X-linked | DDX3X | 1.8 | 3.31 | 2.16 |
| A0A287AYT7 | SEC61 translocon subunit alpha 1 | SEC61A1 | 4.5 | 3.31 | 2.16 |
| I3LJT0 | IF rod domain-containing protein | LOC100525573 | 22.7 | 3.28 | 2.14 |
| P06867 | plasminogen | PLG | -3.4 | 3.27 | 2.13 |
| A0A287AIJ3 | glutathione peroxidase 3 | GPX3 | -2.8 | 3.27 | 2.13 |
| B2D2K7 | solute carrier family 3 member 2 | SLC3A2 | 21.3 | 3.26 | 2.12 |
| P31950 | S100 calcium binding protein A11 | S100A11 | 0.7 | 3.25 | 2.12 |
| A0A287B808 | phosphatidylinositol binding clathrin assembly protein | PICALM | 1.0 | 3.24 | 2.11 |
| A0A287A1G6 | hexokinase 1 | HK1 | 0.8 | 3.20 | 2.07 |
| A0A286ZXH8 | elastase, neutrophil expressed | ELANE | -24.5 | 3.19 | 2.06 |
| F1RKW8 | proteasome 26S subunit, non-ATPase 11 | PSMD11 | 2.8 | 3.16 | 2.04 |
| A0A286ZRK0 | thymopoietin | TMPO | -1.3 | 3.16 | 2.04 |
| G8GCE6 | caveolin 2 | CAV2 | -25.7 | 3.16 | 2.04 |
| F1RWS8 | small nuclear ribonucleoprotein polypeptide A' | SNRPA1 | 23.5 | 3.14 | 2.03 |
| Q6QR67 | resistin | RETN | -13.2 | 3.13 | 2.02 |
| I3LT16 | chromosome 19 open reading frame 25 | C19orf25 | 15.7 | 3.07 | 1.97 |
| P00761 | Trypsinogen isoform X1 | LOC100302368 | -1.8 | 3.08 | 1.97 |
| I3LK68 | phosphofructokinase, platelet | PFKP | -1.1 | 3.07 | 1.96 |
| F1RGJ2 | catenin alpha 1 | CTNNA1 | -0.7 | 3.05 | 1.95 |
| A6M930 | eukaryotic translation initiation factor 4A2 | EIF4A2 | 0.7 | 3.04 | 1.94 |
| Q2EHH8 | protein phosphatase 1 catalytic subunit alpha | PPP1CA | 0.8 | 3.02 | 1.93 |
| A0A286ZUI9 | glycogenin 1 | GYG1 | 3.0 | 3.01 | 1.92 |
| Q2XVP4 | tubulin alpha 1b | TUBA1B | 0.7 | 3.01 | 1.92 |
| I3LEF8 | hydroxysteroid 17-beta dehydrogenase 4 | HSD17B4 | 1.7 | 3.01 | 1.92 |
| F1RPW9 | eukaryotic translation elongation factor 1 gamma | EEF1G | 0.7 | 3.00 | 1.92 |
| F2Z5R6 | cullin associated and neddylation dissociated 1 | CAND1 | 0.8 | 3.00 | 1.91 |
| Q8WNW5 | cadherin 5 | CDH5 | -2.6 | 2.99 | 1.91 |
| I3LFP3 | versican | VCAN | 3.6 | 2.98 | 1.90 |
| D0G7G0 | secreted phosphoprotein 1 | SPP1 | 26.0 | 2.98 | 1.90 |
| K7GNN0 | von Willebrand factor | VWF | -1.8 | 2.97 | 1.90 |
| F1RZT9 | methyl-CpG binding protein 2 | MECP2 | -7.5 | 2.96 | 1.89 |
| P00355 | glyceraldehyde-3-phosphate dehydrogenase | GAPDH | 0.8 | 2.95 | 1.88 |
| F1RXR1 | sushi repeat containing protein X-linked | SRPX | 16.3 | 2.94 | 1.88 |
| F1SVB0 | capping actin protein, gelsolin like | CAPG | 1.9 | 2.94 | 1.87 |
| B9TSR6 | advanced glycosylation end-product specific receptor | AGER | -1.9 | 2.94 | 1.87 |
| F1RQH9 | CD109 molecule | CD109 | 4.8 | 2.93 | 1.87 |
| Q95339 | ATP synthase membrane subunit f | ATP5MF | 1.0 | 2.89 | 1.83 |
| A0A287BPW3 | Protein NPG1 | NPG1 | -2.8 | 2.84 | 1.79 |
| M3UZ96 | uridine-cytidine kinase 1 like 1 | UCKL1 | 25.3 | 2.83 | 1.78 |
| G9F6X8 | prolyl 4-hydroxylase subunit beta | P4HB | 1.5 | 2.83 | 1.78 |
| F1S982 | COPI coat complex subunit beta 1 | COPB1 | 1.0 | 2.83 | 1.78 |
| A0A286ZIH0 | hypoxia up-regulated 1 | HYOU1 | 1.5 | 2.81 | 1.77 |
| F8TEL6 | serine and arginine rich splicing factor 10 | SRSF10 | 0.7 | 2.80 | 1.76 |
| F2Z557 | poly(A) binding protein cytoplasmic 1 | PABPC1 | 1.1 | 2.78 | 1.75 |

| | | | | | |
|---|---|---|---|---|---|
| A0A287AWP8 | complement C5 | C5 | -23.8 | 2.78 | 1.75 |
| A0A286ZSD7 | stomatin like 2 | STOML2 | 4.3 | 2.78 | 1.75 |
| I3LIR9 | autophagy related 9B | ATG9B | -1.5 | 2.76 | 1.73 |
| A0A287BCR9 | junctional cadherin 5 associated | JCAD | -19.9 | 2.76 | 1.73 |
| F1S7E0 | COPI coat complex subunit epsilon | COPE | 23.4 | 2.73 | 1.71 |
| F1RFY1 | profilin 1 | PFN1 | 0.9 | 2.73 | 1.71 |
| F1SR53 | integrin subunit alpha 5 | ITGA5 | 2.9 | 2.72 | 1.70 |
| F1SL24 | CD9 molecule | CD9 | -0.9 | 2.72 | 1.70 |
| A0A287A9I8 | ATP synthase F1 subunit gamma | ATP5F1C | 0.7 | 2.69 | 1.68 |
| Q06AS7 | G protein subunit alpha i2 | GNAI2 | -1.1 | 2.69 | 1.68 |
| A0A286ZJD8 | dipeptidyl peptidase 7 | DPP7 | 3.5 | 2.69 | 1.68 |
| A0A287A784 | keratin 8 | KRT8 | 1.6 | 2.69 | 1.68 |
| F1RRU7 | mannose receptor C type 2 | MRC2 | 2.2 | 2.66 | 1.65 |
| Q9XSI4 | hypoxanthine phosphoribosyltransferase 1 | HPRT1 | -1.9 | 2.65 | 1.64 |
| F1SIT7 | ribosomal protein lateral stalk subunit P1 | RPLP1 | 1.0 | 2.65 | 1.64 |
| I3LPB5 | creatine kinase B | CKB | 2.4 | 2.64 | 1.64 |
| F1SMW4 | vacuolar protein sorting 4 homolog B | VPS4B | 21.3 | 2.63 | 1.64 |
| F1RZ28 | ribosomal protein S10 | RPS10 | 0.9 | 2.63 | 1.64 |
| F1RUM4 | inter-alpha-trypsin inhibitor heavy chain 2 | ITIH2 | -3.1 | 2.64 | 1.64 |
| F1SH96 | inter-alpha-trypsin inhibitor heavy chain 1 | ITIH1 | -4.6 | 2.63 | 1.64 |
| A0A287BKP9 | von Willebrand factor A domain containing 1 | VWA1 | 4.2 | 2.62 | 1.63 |
| F1SQN1 | chaperonin containing TCP1 subunit 4 | CCT4 | 0.9 | 2.62 | 1.63 |
| F2Z5P1 | Histone H2A | H2AFV | -0.9 | 2.62 | 1.63 |
| A0A286ZIU2 | chloride intracellular channel 5 | CLIC5 | -1.9 | 2.61 | 1.62 |
| F1S935 | ribosomal protein L18a | RPL18A | 1.7 | 2.61 | 1.62 |
| F1S4D7 | guanylate binding protein 1 | GBP1 | -3.8 | 2.61 | 1.62 |
| P62863 | FAU ubiquitin like and ribosomal protein S30 fusion | FAU | 1.1 | 2.60 | 1.62 |
| F1S9K5 | Glutamyl-tRNA synthetase | EPRS | 3.2 | 2.58 | 1.60 |
| Q5S1U1 | heat shock protein family B (small) member 1 | HSPB1 | 0.6 | 2.57 | 1.59 |
| A0A287ADK3 | aldehyde dehydrogenase 7 family member A1 | ALDH7A1 | 1.4 | 2.56 | 1.59 |
| A0A286ZV95 | transmembrane p24 trafficking protein 10 | TMED10 | 1.3 | 2.55 | 1.57 |
| I3LC73 | fatty acid synthase | FASN | 0.8 | 2.54 | 1.57 |
| Q29014 | orosomucoid 1 | ORM1 | -22.3 | 2.54 | 1.57 |
| F1RIP4 | RuvB like AAA ATPase 2 | RUVBL2 | 0.6 | 2.53 | 1.56 |
| A0A2C9F350 | acyl-CoA dehydrogenase medium chain | ACADM | 3.8 | 2.52 | 1.56 |
| I3L742 | RuvB like AAA ATPase 1 | RUVBL1 | 0.8 | 2.52 | 1.55 |
| F1SJY1 | endoplasmic reticulum-golgi intermediate compartment 1 | ERGIC1 | 18.5 | 2.51 | 1.55 |
| F1SFF3 | nidogen 2 | NID2 | 1.4 | 2.51 | 1.55 |
| A0A286ZRR3 | selenium binding protein 1 | SELENBP1 | -1.3 | 2.51 | 1.55 |
| A0A287AQP4 | phosphatidylinositol 4-kinase alpha | PI4KA | 17.5 | 2.50 | 1.54 |
| D0G773 | isovaleryl-CoA dehydrogenase | IVD | -2.5 | 2.50 | 1.54 |
| I3LB23 | thioredoxin related transmembrane protein 4 | TMX4 | -3.4 | 2.49 | 1.54 |
| F1S0R7 | NOP2/Sun RNA methyltransferase 2 | NSUN2 | 20.3 | 2.47 | 1.52 |
| A0A287BEC7 | vesicle associated membrane protein 3 | VAMP3 | -1.0 | 2.47 | 1.52 |
| F1SAT9 | thrombomodulin | THBD | -2.5 | 2.46 | 1.51 |

| | | | | | |
|---|---|---|---|---|---|
| K9J4P3 | UDP-glucose glycoprotein glucosyltransferase 1 | UGGT1 | 2.0 | 2.45 | 1.51 |
| F1SM98 | NADH:ubiquinone oxidoreductase core subunit V2 | NDUFV2 | 1.8 | 2.45 | 1.51 |
| Q2VTP6 | FKBP prolyl isomerase 1A | FKBP1A | 1.1 | 2.45 | 1.51 |
| K9J4M9 | collectin subfamily member 12 | COLEC12 | 21.4 | 2.44 | 1.50 |
| A0A287AI92 | carbonic anhydrase 1 | CA1 | -5.1 | 2.44 | 1.50 |
| Q5MJE5 | cathepsin D | CTSD | 1.1 | 2.43 | 1.49 |
| A0A287AIA3 | Yes1 associated transcriptional regulator | YAP1 | -17.7 | 2.39 | 1.46 |
| F1SSC2 | cyclin dependent kinase 13 | CDK13 | 17.7 | 2.38 | 1.45 |
| Q0R678 | Parkinsonism associated deglycase | PARK7 | 0.7 | 2.38 | 1.45 |
| D3Y2W1 | matrix Gla protein | MGP | 6.4 | 2.38 | 1.45 |
| A0A287B2M3 | ribosomal protein SA | RPSA | 1.1 | 2.37 | 1.44 |
| A0A286ZRD4 | tight junction protein 1 | TJP1 | -0.9 | 2.35 | 1.42 |
| P02540 | desmin | DES | 1.8 | 2.34 | 1.41 |
| A0A286ZZM7 | thioredoxin domain containing 17 | TXNDC17 | 5.4 | 2.33 | 1.41 |
| F1S6R7 | polypyrimidine tract binding protein 1 | PTBP1 | 0.6 | 2.33 | 1.41 |
| F1RHJ2 | heparin binding growth factor | HDGF | 0.6 | 2.33 | 1.41 |
| F1RK01 | carboxypeptidase B2 | CPB2 | -20.9 | 2.33 | 1.41 |
| A0A286ZY10 | glucose-6-phosphate dehydrogenase | G6PD | 0.5 | 2.32 | 1.41 |
| F1RM08 | solute carrier family 1 member 5 | SLC1A5 | 17.1 | 2.32 | 1.41 |
| I3LIL4 | myosin IC | MYO1C | -1.0 | 2.32 | 1.41 |
| F8SUW1 | ubiquitin C-terminal hydrolase L1 | UCHL1 | 22.5 | 2.31 | 1.40 |
| P60662 | myosin light chain 6 | MYL6 | 0.5 | 2.31 | 1.40 |
| F1RLC4 | lysyl oxidase | LOX | 18.3 | 2.31 | 1.40 |
| F1SQ89 | cullin associated and neddylation dissociated 2 (putative) | CAND2 | 17.7 | 2.30 | 1.39 |
| F1S6B5 | fibromodulin | FMOD | 2.6 | 2.29 | 1.39 |
| F1SBS4 | complement C3 | C3 | -1.6 | 2.29 | 1.39 |
| A0A287AVG1 | proteasome 26S subunit, ATPase 5 | PSMC5 | 1.0 | 2.28 | 1.38 |
| B3VMR0 | 5-aminoimidazole-4-carboxamide ribonucleotide formyltransferase/IMP cyclohydrolase | ATIC | 6.0 | 2.27 | 1.37 |
| A0A287BSS3 | hemicentin 1 | HMCN1 | -2.6 | 2.27 | 1.37 |
| C3S7K6 | S100 calcium binding protein A9 | S100A9 | -8.9 | 2.26 | 1.37 |
| A0A287BG97 | lanosterol synthase | LSS | 16.5 | 2.25 | 1.36 |
| A0A287BSA8 | ribosomal protein L10a | RPL10A | 1.3 | 2.25 | 1.36 |
| F1SA52 | KN motif and ankyrin repeat domains 3 | KANK3 | -18.1 | 2.25 | 1.36 |
| F1S564 | RNA 3'-terminal phosphate cyclase | RTCA | 5.7 | 2.24 | 1.35 |
| A0A287BRY0 | tropomyosin 1 | TPM1 | -2.0 | 2.24 | 1.35 |
| K9J4V0 | small nuclear ribonucleoprotein U5 subunit 200 | SNRNP200 | 2.2 | 2.23 | 1.35 |
| A0A286ZRN8 | spectrin beta, non-erythrocytic 2 | SPTBN2 | 4.0 | 2.22 | 1.34 |
| F1RQT6 | VPS51 subunit of GARP complex | VPS51 | 17.0 | 2.22 | 1.34 |
| A0A287AGN9 | spondin 1 | SPON1 | 21.1 | 2.22 | 1.34 |
| F1S431 | Alanine--tRNA ligase | AARS | 3.8 | 2.20 | 1.32 |
| I3LCP8 | serpin family B member 6 | SERPINB6 | -2.7 | 2.20 | 1.32 |
| F1S8B0 | DDRGK domain containing 1 | DDRGK1 | 3.7 | 2.18 | 1.31 |
| A5GFU6 | GNAS complex locus | GNAS | -0.7 | 2.18 | 1.31 |
| K7GK75 | cofilin 1 | CFL1 | 0.9 | 2.18 | 1.31 |
| Q06AT6 | ras homolog family member G | RHOG | -0.8 | 2.17 | 1.30 |

| | | | | | |
|---|---|---|---|---|---|
| F1S0M9 | FKBP prolyl isomerase 10 | FKBP10 | 20.7 | 2.17 | 1.30 |
| **PVH Whole Lung vs. Sham Whole Lung** | | | | | |
| K9IW97 | Egf-like module containing, mucin-like, hormone receptor-like 4 protein | EMR4 | -23.0 | 27.05 | 23.24 |
| I3LKU0 | Rac family small GTPase 2 | RAC2 | -2.2 | 23.41 | 19.91 |
| P22411 | dipeptidyl peptidase 4 | DPP4 | -4.2 | 20.26 | 16.93 |
| F1RSE2 | transmembrane protein 100 | TMEM100 | -18.3 | 18.05 | 14.84 |
| I3LQF4 | semaphorin 7A (John Milton Hagen blood group) | SEMA7A | -23.9 | 16.49 | 13.38 |
| I3LDC2 | RRAD and GEM like GTPase 1 | REM1 | 2.7 | 15.54 | 12.51 |
| I3LKF3 | fascin actin-bundling protein 1 | FSCN1 | 1.7 | 13.73 | 10.76 |
| M3V830 | death associated protein kinase 3 | DAPK3 | 20.3 | 12.97 | 10.11 |
| F1S6S9 | proteinase 3 | PRTN3 | -4.9 | 12.99 | 10.11 |
| A0A286ZZK2 | CD47 molecule | CD47 | -1.7 | 12.89 | 10.08 |
| F1SUU4 | filamin binding LIM protein 1 | FBLIM1 | 4.6 | 12.71 | 9.95 |
| A0A287AXP1 | cyclin dependent kinase 2 | CDK2 | 16.0 | 12.60 | 9.87 |
| Q6QR67 | resistin | RETN | -4.5 | 12.50 | 9.81 |
| F1SLA4 | solute carrier organic anion transporter family member 2A1 | SLCO2A1 | -17.0 | 12.27 | 9.61 |
| F1RRX1 | lipocalin 2 | LCN2 | -2.9 | 12.00 | 9.36 |
| P80015 | azurocidin 1 | AZU1 | -3.6 | 11.39 | 8.78 |
| A0A287APV0 | serpin family B member 10 | SERPINB10 | -2.0 | 11.32 | 8.75 |
| A0A287AN90 | matrix metallopeptidase 9 | MMP9 | -4.3 | 10.55 | 7.99 |
| F1RIE0 | mesoderm development LRP chaperone | MESD | 1.6 | 10.34 | 7.84 |
| Q8WMN8 | lactotransferrin | LTF | -3.5 | 10.34 | 7.84 |
| Q0Z8U2 | ribosomal protein S3 | RPS3 | 1.5 | 10.21 | 7.72 |
| F1RQ08 | phospholipase C beta 3 | PLCB3 | -1.4 | 9.89 | 7.43 |
| F1SF78 | CTP synthase 1 | CTPS1 | 19.4 | 9.83 | 7.38 |
| A0A287B749 | tubulin beta 6 class V | TUBB6 | 2.6 | 9.77 | 7.35 |
| F1RUJ4 | guanosine monophosphate reductase | GMPR | -3.6 | 9.74 | 7.33 |
| I3LEQ7 | serine/threonine kinase receptor associated protein | STRAP | 1.3 | 9.51 | 7.12 |
| A0A287BHR4 | deoxyribonuclease 1 like 1 | DNASE1L1 | -2.0 | 9.46 | 7.08 |
| F1RYS5 | Septin-11 | SPTAN11 | 1.2 | 9.42 | 7.06 |
| F1RMW7 | CD177 molecule | CD177 | -4.1 | 8.91 | 6.56 |
| K7GR72 | solute carrier family 4 member 1 (Diego blood group) | SLC4A1 | -4.5 | 8.79 | 6.46 |
| A0A287B7T3 | phospholipase B domain containing 1 | PLBD1 | -3.8 | 8.70 | 6.38 |
| K7GSR6 | 5'-nucleotidase ecto | NT5E | -3.1 | 8.68 | 6.38 |
| A0A287B481 | aldehyde oxidase 1 | AOX1 | 15.7 | 8.50 | 6.22 |
| F1SKX8 | filamin A interacting protein 1 like | FILIP1L | 5.2 | 8.51 | 6.22 |
| F1RRT3 | DUF1394 domain-containing protein | FAM49B | -1.4 | 8.45 | 6.19 |
| A0A286ZXH8 | elastase, neutrophil expressed | ELANE | -5.9 | 8.37 | 6.12 |
| A0A287BC64 | TEK receptor tyrosine kinase | TEK | -2.2 | 8.26 | 6.03 |
| F1S682 | quiescin sulfhydryl oxidase 1 | QSOX1 | -3.0 | 8.26 | 6.03 |
| F1SJB9 | Hyaluronoglucosaminidase | TMEM2 | -3.4 | 8.25 | 6.03 |
| F1SAD9 | protein disulfide isomerase family A member 4 | PDIA4 | 1.7 | 8.23 | 6.02 |
| F1RQR3 | CDC42 binding protein kinase gamma | CDC42BPG | -21.0 | 8.14 | 5.94 |
| I3LR64 | iron-sulfur cluster assembly enzyme | ISCU | 16.6 | 8.09 | 5.91 |
| A0A173G6H0 | Septin-5 | SPTAN5 | 2.1 | 8.07 | 5.89 |

| | | | | | |
|---|---|---|---|---|---|
| A0A287B182 | transmembrane protein 263 | TMEM263 | 19.4 | 8.01 | 5.85 |
| B2CZR7 | CFL2b variant 1 | CFL2b | 0.9 | 8.01 | 5.85 |
| K7GNJ2 | integrin subunit alpha L | ITGAL | -1.7 | 7.92 | 5.77 |
| A0A287AXG3 | four and a half LIM domains 2 | FHL2 | 3.0 | 7.84 | 5.71 |
| A0A286ZR49 | Prophenin and tritrpticin precursor | PMAP-23 | -4.1 | 7.82 | 5.71 |
| A0A173G6G4 | Septin 4 | SPTAN4 | 1.7 | 7.80 | 5.70 |
| F1SP36 | PDZ domain containing 2 | PDZD2 | -20.0 | 7.80 | 5.70 |
| A0A287B3S9 | microtubule associated protein 2 | MAP2 | 15.0 | 7.79 | 5.70 |
| A0A286ZW70 | peptidylprolyl isomerase B | PPIB | 1.6 | 7.76 | 5.67 |
| F1S7U3 | chitinase 3 like 1 | CHI3L1 | 5.0 | 7.73 | 5.65 |
| Q95JA4 | 72 kDa gelatinase | MMP-2 | 2.0 | 7.72 | 5.65 |
| Q95N21 | FcgammaRIII a.2 | FCGR3A | -2.0 | 7.59 | 5.53 |
| K7GNN0 | von Willebrand factor | VWF | -2.7 | 7.57 | 5.52 |
| F2Z5B1 | serpin family B member 1 | SERPINB1 | -1.6 | 7.54 | 5.49 |
| F1RWV1 | complement factor properdin | CFP | -3.2 | 7.50 | 5.47 |
| I3LTD0 | lymphatic vessel endothelial hyaluronan receptor 1 | LYVE1 | -3.8 | 7.50 | 5.47 |
| A0A287AUD4 | coiled-coil domain containing 167 | CCDC167 | 14.4 | 7.39 | 5.37 |
| A0A287BHQ8 | caspase 1 | CASP1 | -2.2 | 7.29 | 5.29 |
| A0A286ZNV5 | aldehyde dehydrogenase 9 family member A1 | ALDH9A1 | -1.1 | 7.23 | 5.23 |
| A0A287BA24 | glycine cleavage system protein H | GCSH | 15.6 | 7.21 | 5.22 |
| F1SPH1 | proliferation-associated 2G4 | PA2G4 | 1.6 | 7.13 | 5.14 |
| Q764N2 | CD3d molecule | CD3D | -10.5 | 7.10 | 5.12 |
| A0A1L6ZA05 | Lymphocyte antigen 6 complex locus G6F | ly6g6f | -15.0 | 7.06 | 5.08 |
| A0A286ZSB2 | RRAD, Ras related glycolysis inhibitor and calcium channel regulator | RRAD | 16.1 | 6.99 | 5.02 |
| F1RRU7 | mannose receptor C type 2 | MRC2 | 1.3 | 6.92 | 4.96 |
| F1RGJ3 | heat shock protein family A (Hsp70) member 9 | HSPA9 | 1.6 | 6.88 | 4.92 |
| A0A287BJ70 | coagulation factor V | F5 | -4.1 | 6.87 | 4.92 |
| A0A287BAZ3 | erythrocyte membrane protein band 4.1 | EPB41 | -4.9 | 6.87 | 4.92 |
| A0A287A8W9 | Septin-2 | SPTAN2 | 0.8 | 6.85 | 4.91 |
| A0A286ZI97 | peptidoglycan recognition protein 1 | PGLYRP1 | -4.0 | 6.79 | 4.85 |
| A0PA01 | Serine protease inhibitor 9 | PI-9 | -1.2 | 6.68 | 4.75 |
| F1RQB6 | G3BP stress granule assembly factor 1 | G3BP1 | 1.8 | 6.65 | 4.73 |
| I3LNY6 | nestin | NES | 2.8 | 6.64 | 4.73 |
| A0A287AB49 | mevalonate diphosphate decarboxylase | MVD | 1.8 | 6.63 | 4.72 |
| A0A287A0I8 | drebrin 1 | DBN1 | 2.4 | 6.55 | 4.65 |
| A0A286ZMZ2 | cleavage and polyadenylation specific factor 6 | CPSF6 | 1.8 | 6.55 | 4.65 |
| A0A287BCM3 | IKBKB interacting protein | IKBIP | 2.5 | 6.54 | 4.64 |
| F1S1C3 | carbonic anhydrase 4 | CA4 | -4.9 | 6.40 | 4.51 |
| A0A287AP66 | ribosomal protein L12 | RPL12 | 1.1 | 6.38 | 4.50 |
| F1RSE1 | phosphatidylcholine transfer protein | PCTP | -2.6 | 6.33 | 4.45 |
| E1CAJ6 | Protein disulfide isomerase P5 | pdi-p5 | 1.8 | 6.30 | 4.43 |
| A0A287ARZ1 | thioredoxin domain containing 5 | TXNDC5 | 2.0 | 6.30 | 4.43 |
| F1SL24 | CD9 molecule | CD9 | -1.3 | 6.28 | 4.42 |
| M3VK46 | solute carrier family 44 member 2 | SLC44A2 | -1.7 | 6.25 | 4.39 |
| F1SCY9 | ankyrin repeat domain 22 | ANKRD22 | -10.9 | 6.25 | 4.39 |

| | | | | | |
|---|---|---|---|---|---|
| I3LL91 | CD200 molecule | CD200 | -2.8 | 6.19 | 4.34 |
| A0A287BMM2 | beta 3-glucosyltransferase | B3GLCT | 2.4 | 6.17 | 4.33 |
| F1RF77 | procollagen-lysine,2-oxoglutarate 5-dioxygenase 1 | PLOD1 | 2.9 | 6.16 | 4.32 |
| A0A286ZIJ5 | NADPH oxidase 1 | NOX1 | -1.6 | 6.15 | 4.32 |
| F1RRW5 | angiotensin I converting enzyme | ACE | -2.0 | 6.13 | 4.30 |
| B4YYD8 | nitric oxide synthase 3 | NOS3 | -16.7 | 6.07 | 4.24 |
| A0A287BPW3 | Uncharacterized protein | LOC110258651 | -2.8 | 6.06 | 4.24 |
| P63246 | receptor for activated C kinase 1 | RACK1 | 1.5 | 6.04 | 4.22 |
| G9F6X8 | prolyl 4-hydroxylase subunit beta | P4HB | 1.7 | 6.02 | 4.20 |
| B2ZI35 | F11 receptor | F11R | -1.4 | 5.89 | 4.09 |
| A0A287BT04 | prolyl 3-hydroxylase 1 | P3H1 | 2.6 | 5.88 | 4.07 |
| F1S798 | endothelial cell adhesion molecule | ESAM | -1.2 | 5.87 | 4.07 |
| Q6SZ84 | CD45 antigen isoform 2 | CD45 | -1.5 | 5.87 | 4.07 |
| F1RYJ8 | cysteine and glycine rich protein 2 | CSRP2 | 1.6 | 5.82 | 4.03 |
| A0A287A699 | heterogeneous nuclear ribonucleoprotein K | HNRNPK | 1.6 | 5.77 | 3.99 |
| F1RLG4 | signal recognition particle 19 | SRP19 | 2.1 | 5.69 | 3.91 |
| F8SIP2 | EGF containing fibulin extracellular matrix protein 1 | EFEMP1 | -1.2 | 5.67 | 3.89 |
| I3LS60 | ATP-dependent (S)-NAD(P)H-hydrate dehydratase | NAXD | 0.9 | 5.62 | 3.85 |
| Q9TUN6 | Glycoprotein IIb | GPIIb | -4.2 | 5.62 | 3.85 |
| C3S7K6 | S100 calcium binding protein A9 | S100A9 | -2.6 | 5.60 | 3.83 |
| A0A286ZXM3 | Tryptase | MCT7 | -3.3 | 5.57 | 3.81 |
| F1S0M9 | FKBP prolyl isomerase 10 | FKBP10 | 3.3 | 5.56 | 3.80 |
| F1SC51 | PDZ and LIM domain 1 | PDLIM1 | 1.7 | 5.51 | 3.76 |
| A0A286ZIK9 | phospholipid phosphatase 1 | PLPP1 | -1.1 | 5.51 | 3.76 |
| F2Z567 | ribosomal protein L8 | RPL8 | 2.3 | 5.49 | 3.75 |
| A6M931 | eukaryotic translation initiation factor 4A3 | EIF4A3 | 1.0 | 5.47 | 3.73 |
| I3LP35 | Peptidase S1 domain-containing protein | LOC100154047 | -8.8 | 5.46 | 3.72 |
| F1RNB8 | spectrin alpha, erythrocytic 1 | SPTA1 | -3.6 | 5.45 | 3.71 |
| I3LV51 | zinc finger CCCH-type and G-patch domain containing | ZGPAT | 11.9 | 5.44 | 3.70 |
| F1S3I3 | eukaryotic translation initiation factor 3 subunit G | EIF3G | 1.3 | 5.41 | 3.68 |
| A0A287A2G6 | glutathione peroxidase 7 | GPX7 | 1.9 | 5.40 | 3.67 |
| F1SDK3 | O-6-methylguanine-DNA methyltransferase | MGMT | 1.3 | 5.35 | 3.63 |
| F1SNU4 | Uncharacterized protein | LOC100736951 | -5.5 | 5.35 | 3.63 |
| A0A286ZXA0 | insulin like growth factor binding protein 7 | IGFBP7 | 3.6 | 5.32 | 3.60 |
| F1SS98 | MINDY lysine 48 deubiquitinase 1 | MINDY1 | -6.0 | 5.27 | 3.56 |
| A0A287A080 | annexin A13 | ANXA13 | -4.6 | 5.26 | 3.56 |
| Q29097 | selectin P | SELP | -8.0 | 5.26 | 3.56 |
| I3LMA9 | syntaxin binding protein 5 like | STXBP5L | 11.7 | 5.22 | 3.52 |
| A0A287B2M3 | ribosomal protein SA | RPSA | 1.5 | 5.21 | 3.52 |
| D7RA22 | autophagy related 4B cysteine peptidase | ATG4B | 13.2 | 5.20 | 3.51 |
| A0A287A113 | spectrin beta, erythrocytic | SPTB | -3.9 | 5.17 | 3.48 |
| F1RMK7 | zinc finger protein 787 | ZNF787 | 2.2 | 5.10 | 3.42 |
| I3LG64 | paired immunoglobin like type 2 receptor alpha | PILRA | -15.1 | 5.10 | 3.42 |
| A0A287B3Y1 | eukaryotic translation initiation factor 3 subunit I | EIF3I | 0.8 | 5.09 | 3.41 |
| F1S227 | peptidyl-tRNA hydrolase 2 | PTRH2 | 1.0 | 5.04 | 3.37 |
| I3LSP1 | methionyl aminopeptidase 2 | METAP2 | 1.7 | 5.02 | 3.35 |

| | | | | | |
|---|---|---|---|---|---|
| A0A287B9R2 | matrix metallopeptidase 14 | MMP14 | 13.4 | 5.01 | 3.34 |
| F1S5D8 | protein tyrosine phosphatase receptor type C | PTPRC | -0.9 | 4.96 | 3.30 |
| F1STD6 | zeta chain of T cell receptor associated protein kinase 70 | ZAP70 | -13.3 | 4.96 | 3.30 |
| F1S9Q1 | heat shock protein family A (Hsp70) member 8 | HSPA8 | 1.0 | 4.95 | 3.29 |
| F2Z5T3 | chromobox 5 | CBX5 | 2.1 | 4.93 | 3.27 |
| B6CVD6 | Endoplasmic reticulum resident protein 44 | ERP44 | 1.1 | 4.92 | 3.27 |
| A0A076KWW8 | C-type lectin domain family 8 member A | CLEC8A | -5.9 | 4.92 | 3.27 |
| F1S1U5 | golgi reassembly stacking protein 2 | GORASP2 | 2.5 | 4.91 | 3.27 |
| K7GRK7 | tenascin XB | TNXB | -2.8 | 4.87 | 3.23 |
| I6YMD1 | interferon induced protein with tetratricopeptide repeats 2 | IFIT2 | -16.7 | 4.86 | 3.22 |
| F1RRP1 | myeloperoxidase | MPO | -4.8 | 4.86 | 3.22 |
| A0A287AHH7 | uridine-cytidine kinase 2 | UCK2 | 12.5 | 4.83 | 3.20 |
| I3LNH3 | glucosidase II alpha subunit | GANAB | 0.8 | 4.82 | 3.19 |
| F1RRV1 | tetratricopeptide repeat, ankyrin repeat and coiled-coil containing 2 | TANC2 | -21.9 | 4.81 | 3.18 |
| A0A287A233 | arrestin beta 2 | ARRB2 | -3.7 | 4.80 | 3.18 |
| A0A287A168 | adhesion G protein-coupled receptor E1 | ADGRE1 | -8.2 | 4.80 | 3.18 |
| Q8WNW8 | Nexin-1 | PN-1 | 2.2 | 4.79 | 3.17 |
| A0A286ZL07 | BUB3 mitotic checkpoint protein | BUB3 | 0.9 | 4.76 | 3.15 |
| P32394 | heme oxygenase 1 | HMOX1 | -1.4 | 4.75 | 3.14 |
| A0A287BIL8 | heat shock protein family A (Hsp70) member 5 | HSPA5 | 2.0 | 4.75 | 3.13 |
| Q06AT9 | RNA binding motif protein 4B | RBM4B | 14.7 | 4.72 | 3.11 |
| P30034 | platelet factor 4 | PF4 | -3.7 | 4.71 | 3.10 |
| A5A8V6 | Heat shock 70kDa protein 1A | HSPA1A | 0.7 | 4.70 | 3.10 |
| C4NF76 | bone marrow stromal cell antigen 2 | BST2 | -3.0 | 4.68 | 3.07 |
| P62272 | ribosomal protein S18 | RPS18 | 1.6 | 4.67 | 3.07 |
| F1RYA3 | Cysteinyl-tRNA synthetase | CARS1 | 1.5 | 4.65 | 3.05 |
| F1RLQ5 | CAP10 domain-containing protein | KDELC1 | 2.8 | 4.63 | 3.04 |
| M3V819 | collagen type IV alpha 1 chain | COL4A1 | 2.0 | 4.61 | 3.02 |
| Q03472 | Apolipoprotein R | APOR | -2.9 | 4.61 | 3.02 |
| A0A287A8F1 | filamin A interacting protein 1 | FILIP1 | 11.5 | 4.58 | 2.99 |
| F1SK31 | occludin | OCLN | -2.3 | 4.56 | 2.98 |
| F1S3D5 | lectin, mannose binding 2 | LMAN2 | 1.0 | 4.55 | 2.97 |
| A0A2C9F380 | phosphoglucomutase 3 | PGM3 | 1.2 | 4.52 | 2.94 |
| F2Z5T8 | Mps one binder kinase activator-like 3 | MOBKL3 | 0.8 | 4.51 | 2.94 |
| F1RSW6 | RNA polymerase I and III subunit D | POLR1D | 16.7 | 4.49 | 2.92 |
| F1SAZ6 | tryptophanyl tRNA synthetase 2, mitochondrial | WARS2 | 1.8 | 4.49 | 2.92 |
| F1RX36 | fibrinogen alpha chain | FGA | -3.9 | 4.48 | 2.91 |
| F1SMS8 | lectin, mannose binding 1 | LMAN1 | 1.9 | 4.46 | 2.89 |
| A0A287AG39 | small nuclear ribonucleoprotein polypeptide N | SNRPN | 1.7 | 4.44 | 2.87 |
| I3LT81 | 60S ribosomal protein L17 | RPL17-C18orf | 1.6 | 4.44 | 2.87 |
| A0A287BAC8 | syntaxin binding protein 6 | STXBP6 | -12.1 | 4.42 | 2.86 |
| I3LLQ4 | BMP binding endothelial regulator | BMPER | -2.4 | 4.40 | 2.84 |
| P51636 | caveolin 2 | CAV2 | -1.3 | 4.39 | 2.84 |
| P02543 | vimentin | VIM | 1.5 | 4.37 | 2.82 |

| | | | | | |
|---|---|---|---|---|---|
| F1SMJ6 | complement C9 | C9 | -3.7 | 4.36 | 2.81 |
| F1SH84 | bridging integrator 2 | BIN2 | -2.7 | 4.36 | 2.81 |
| Q52NJ3 | secretion associated Ras related GTPase 1A | SAR1A | 1.1 | 4.35 | 2.80 |
| K7GQL2 | coagulation factor XIII A chain | F13A1 | -2.3 | 4.35 | 2.80 |
| Q6QA76 | PDZ domain containing 11 | PDZD11 | 17.3 | 4.34 | 2.80 |
| A0A287BM03 | Enah/Vasp-like | EVL | -6.3 | 4.33 | 2.79 |
| D0G6X9 | 15-hydroxyprostaglandin dehydrogenase | HPGD | -6.5 | 4.32 | 2.79 |
| F1S765 | CXXC motif containing zinc binding protein | C1orf123 | 1.0 | 4.32 | 2.78 |
| F1RJM2 | endoplasmic reticulum protein 29 | ERP29 | 1.3 | 4.31 | 2.78 |
| F1S870 | proteasome inhibitor subunit 1 | PSMF1 | -2.2 | 4.31 | 2.78 |
| I3LDC7 | isocitrate dehydrogenase (NADP(+)) 1 | IDH1 | 1.3 | 4.26 | 2.74 |
| A0A287AMZ5 | CD46 molecule | CD46 | -1.2 | 4.26 | 2.74 |
| P06867 | plasminogen | PLG | -3.6 | 4.26 | 2.74 |
| A0A287AX97 | collagen type XV alpha 1 chain | COL15A1 | 1.3 | 4.25 | 2.73 |
| A0A287AM78 | purine nucleoside phosphorylase | PNP | -1.6 | 4.23 | 2.72 |
| I3LI80 | advillin | AVIL | -3.6 | 4.24 | 2.72 |
| A0A287A8N4 | C-X-C motif chemokine | LOC100520680 | -4.9 | 4.21 | 2.70 |
| K9IVJ5 | serine dehydratase | SDS | -16.0 | 4.19 | 2.68 |
| A0A287ACK3 | DAB adaptor protein 2 | DAB2 | 2.2 | 4.17 | 2.67 |
| Q29205 | ribosomal protein L11 | RPL11 | 0.8 | 4.17 | 2.67 |
| K7GM61 | solute carrier family 44 member 1 | SLC44A1 | -1.0 | 4.17 | 2.67 |
| F1SLM7 | glutathione peroxidase 8 (putative) | GPX8 | 1.9 | 4.13 | 2.64 |
| F6Q5P0 | ribosomal protein S13 | RPS13 | 0.9 | 4.08 | 2.59 |
| F1RUM4 | inter-alpha-trypsin inhibitor heavy chain 2 | ITIH2 | -3.3 | 4.08 | 2.59 |
| Q8SQ44 | Tryptase | LOC396700 | -13.7 | 4.08 | 2.59 |
| F1RM25 | protein phosphatase 5 catalytic subunit | PPP5C | 1.0 | 4.07 | 2.58 |
| A0A286ZYS2 | Aspartate--tRNA ligase, cytoplasmic | DARS | 1.9 | 4.07 | 2.58 |
| A0A287AJN3 | UPF0687 protein C20orf27 isoform 2 | C17H20orf27P | -1.0 | 4.06 | 2.57 |
| F1SU88 | MER proto-oncogene, tyrosine kinase | MERTK | -7.0 | 4.04 | 2.56 |
| F1SA03 | A-kinase anchoring protein 8 like | AKAP8L | 2.1 | 4.01 | 2.53 |
| P02540 | desmin | DES | 1.6 | 4.01 | 2.53 |
| F1RQ75 | coagulation factor IX | F9 | -2.9 | 4.00 | 2.53 |
| A0A287AKQ5 | LSM4 homolog, U6 small nuclear RNA and mRNA degradation associated | LSM4 | 1.7 | 4.00 | 2.53 |
| K9J6H4 | Sialic acid-binding Ig-like lectin 5-like 1 protein | SIGLEC5L1 | -14.6 | 3.99 | 2.52 |
| I3LR51 | FKBP prolyl isomerase 3 | FKBP3 | 1.1 | 3.97 | 2.50 |
| F1STZ4 | complement C1q A chain | C1QA | -2.0 | 3.97 | 2.50 |
| A0A287AUT0 | GB1/RHD3-type G domain-containing protein | LOC100523668 | -2.2 | 3.97 | 2.50 |
| F1SBI1 | sorting nexin 5 | SNX5 | 0.9 | 3.95 | 2.48 |
| F1RRB7 | acetyl-CoA acyltransferase 1 | ACAA1 | 1.2 | 3.94 | 2.48 |
| F6M2L9 | Integrin alpha-M | Itgam | -10.3 | 3.93 | 2.47 |
| Q9XSH0 | folate receptor alpha | FOLR1 | -3.6 | 3.91 | 2.45 |
| F1SPP8 | cytoskeleton associated protein 4 | CKAP4 | 2.0 | 3.87 | 2.41 |
| Q06A96 | small nuclear ribonucleoprotein polypeptide B2 | SNRPB2 | 1.3 | 3.87 | 2.41 |
| F1RJ25 | aldolase, fructose-bisphosphate C | ALDOC | -1.0 | 3.85 | 2.40 |
| I3LFE2 | chromosome unknown C11orf98 homolog | C2H11orf98 | 13.8 | 3.84 | 2.40 |

| | | | | | |
|---|---|---|---|---|---|
| A0A286ZKH7 | paralemmin | PALM | -2.0 | 3.85 | 2.40 |
| C3S7K5 | S100 calcium binding protein A8 | S100A8 | -2.7 | 3.84 | 2.39 |
| F1S3J8 | mitochondrial ribosomal protein L4 | MRPL4 | 11.5 | 3.83 | 2.39 |
| A0A287BFH9 | alpha tocopherol transfer protein like | TTPAL | 10.9 | 3.83 | 2.39 |
| B2LWN5 | C1q and TNF related 3 | C1QTNF3 | -11.4 | 3.82 | 2.38 |
| K9IWD4 | Hypoxia up-regulated protein 1 | HYOU1_tv1 | 1.2 | 3.81 | 2.38 |
| A0A287BRF1 | C1q domain-containing protein | LOC110258309 | -3.6 | 3.80 | 2.36 |
| F1SH96 | inter-alpha-trypsin inhibitor heavy chain 1 | ITIH1 | -3.3 | 3.77 | 2.34 |
| A0A287A189 | Cytochrome c oxidase assembly protein COX20, mitochondrial | COX20 | 11.5 | 3.77 | 2.34 |
| A0A287BG45 | tetraspanin 15 | TSPAN15 | -12.3 | 3.75 | 2.32 |
| A0A287AJG4 | ankyrin repeat domain 24 | ANKRD24 | 15.7 | 3.74 | 2.31 |
| K9IVT8 | DEAD-box helicase 41 | DDX41 | 15.3 | 3.73 | 2.31 |
| F1S8S6 | WD repeat domain 26 | WDR26 | 13.5 | 3.73 | 2.31 |
| Q9TV77 | protein phosphatase 1 regulatory subunit 12A | PPP1R12A | 1.4 | 3.73 | 2.31 |
| Q1ACV4 | transporter 2, ATP binding cassette subfamily B member | TAP2 | -14.9 | 3.74 | 2.31 |
| I3LSL1 | paired immunoglobin like type 2 receptor alpha | LOC100514951 | -17.9 | 3.73 | 2.31 |
| I3L6R1 | cartilage associated protein | CRTAP | 2.1 | 3.73 | 2.31 |
| A0A287AG70 | glutathione peroxidase 1 | GPX1 | -1.3 | 3.71 | 2.29 |
| F1SB33 | synaptophysin like 1 | SYPL1 | -1.2 | 3.70 | 2.28 |
| C9K507 | CDK-activating kinase assembly factor MAT1 | MAT1 | 10.5 | 3.69 | 2.28 |
| F1SBB3 | laminin subunit alpha 3 | LAMA3 | -2.0 | 3.69 | 2.28 |
| F1RVZ1 | acyl-CoA oxidase 1 | ACOX1 | 2.4 | 3.68 | 2.27 |
| F1SJS8 | transgelin | TAGLN | 1.1 | 3.68 | 2.27 |
| I3LG73 | ribosomal L1 domain containing 1 | RSL1D1 | 14.0 | 3.67 | 2.26 |
| I3LP78 | ribosomal protein L9 | RPL9 | 1.0 | 3.67 | 2.26 |
| F1RJU9 | Septin-8 | SPTAN8 | 1.4 | 3.66 | 2.26 |
| A0A287AXD1 | myocardial zonula adherens protein | MYZAP | -2.3 | 3.64 | 2.24 |
| A1X898 | prolyl 4-hydroxylase subunit alpha 1 | P4HA1 | 1.8 | 3.63 | 2.23 |
| F1RQT6 | VPS51 subunit of GARP complex | VPS51 | -7.6 | 3.63 | 2.23 |
| F1SRA1 | GRB2 related adaptor protein 2 | GRAP2 | -14.0 | 3.62 | 2.22 |
| F1SSG6 | N-acetylneuraminate synthase | NANS | 1.0 | 3.61 | 2.22 |
| A0A287A3C2 | coagulation factor XII | F12 | -3.8 | 3.61 | 2.22 |
| F1SD24 | BPTI/Kunitz inhibitor domain-containing protein | PTI | -10.9 | 3.61 | 2.22 |
| K9IVW2 | integrin subunit alpha X | ITGAX | -1.5 | 3.60 | 2.22 |
| A0A287A8H5 | N-acetylated alpha-linked acidic dipeptidase like 2 | NAALADL2 | -7.9 | 3.60 | 2.22 |
| F1SJ86 | chondroitin sulfate proteoglycan 4 | CSPG4 | 2.5 | 3.60 | 2.21 |
| D6BR76 | Integrin beta | GPIIIa | -2.3 | 3.60 | 2.21 |
| F1RQW5 | negative elongation factor complex member E | NELFE | 1.0 | 3.59 | 2.21 |
| F1RYZ1 | CD151 molecule (Raph blood group) | CD151 | -1.0 | 3.59 | 2.21 |
| I3L816 | heterogeneous nuclear ribonucleoprotein H1 | HNRNPH1 | 1.9 | 3.58 | 2.20 |
| F1SJ68 | mannosidase alpha class 2C member 1 | MAN2C1 | -1.5 | 3.58 | 2.20 |
| F1RL75 | platelet derived growth factor receptor beta | PDGFRB | 1.7 | 3.57 | 2.20 |
| A0A287B8B1 | ST13 Hsp70 interacting protein | ST13 | 0.7 | 3.57 | 2.20 |
| F1SJ07 | pleckstrin | PLEK | -6.4 | 3.57 | 2.20 |
| Q6QAT0 | ribosomal protein L32 | RPL32 | 2.3 | 3.57 | 2.20 |

| | | | | | |
|---|---|---|---|---|---|
| A0A286ZPJ4 | coiled-coil domain containing 85A | CCDC85A | -13.6 | 3.56 | 2.19 |
| A0A287BLW6 | plexin B2 | PLXNB2 | 0.8 | 3.55 | 2.19 |
| I3LH49 | glutaminyl-peptide cyclotransferase | QPCT | -12.9 | 3.55 | 2.19 |
| F1SKI6 | NADH:ubiquinone oxidoreductase complex assembly factor 3 | NDUFAF3 | 11.3 | 3.54 | 2.18 |
| A0A287A2P6 | actin like 6A | ACTL6A | 1.1 | 3.54 | 2.18 |
| F1STM9 | replication protein A2 | RPA2 | 0.7 | 3.53 | 2.18 |
| F1RVI6 | phosphatidylinositol-5-phosphate 4-kinase type 2 alpha | PIP4K2A | -2.2 | 3.52 | 2.17 |
| A0A287BI62 | lymphocyte specific protein 1 | LSP1 | -1.7 | 3.52 | 2.17 |
| B6VAP9 | apurinic/apyrimidinic endodeoxyribonuclease 1 | APEX1 | 1.1 | 3.49 | 2.14 |
| I3L6U9 | G protein subunit alpha 11 | GNA11 | -1.3 | 3.48 | 2.13 |
| I3LIC4 | FLVCR heme transporter 2 | FLVCR2 | -16.3 | 3.48 | 2.13 |
| A0A287A874 | proline-serine-threonine phosphatase interacting protein 2 | PSTPIP2 | -12.1 | 3.48 | 2.13 |
| K7GNP5 | secreted frizzled related protein 1 | SFRP1 | 14.7 | 3.47 | 2.12 |
| A0A287ANN7 | eukaryotic translation elongation factor 1 delta | EEF1D | 1.3 | 3.46 | 2.12 |
| F1RKY2 | serpin family D member 1 | SERPIND1 | -2.4 | 3.46 | 2.12 |
| A0A287AE25 | Non-secretory ribonuclease | LOC102163838 | -9.2 | 3.46 | 2.12 |
| A0A287B6I2 | splicing factor 1 | SF1 | 1.0 | 3.46 | 2.11 |
| A0A286ZWI9 | rhotekin 2 | RTKN2 | -9.6 | 3.45 | 2.11 |
| A0A287B388 | complement factor H | CFH | -3.1 | 3.45 | 2.11 |
| A0A287B929 | ribosomal protein L13a | RPL13A | 2.3 | 3.44 | 2.11 |
| F1SUE0 | BICD cargo adaptor 2 | BICD2 | 1.9 | 3.44 | 2.11 |
| A0A287AXF7 | family with sequence similarity 162 member A | FAM162A | 1.8 | 3.44 | 2.11 |
| A0A287ALU2 | matrix metallopeptidase 8 | MMP8 | -13.9 | 3.44 | 2.11 |
| F1RQW6 | complement factor B | CFB | -3.0 | 3.43 | 2.10 |
| Q8WN93 | calcitonin receptor like receptor | CALCRL | -14.6 | 3.43 | 2.10 |
| I3LJW2 | fibrinogen gamma chain | FGG | -3.2 | 3.43 | 2.10 |
| A0A286ZTE1 | sialic acid binding Ig like lectin 1 | SIGLEC1 | -2.0 | 3.42 | 2.10 |
| F2Z5M9 | signal recognition particle 54 | SRP54 | 1.6 | 3.40 | 2.08 |
| B6E240 | 3-phosphoinositide dependent protein kinase 1 | PDPK1 | 10.8 | 3.40 | 2.08 |
| I3L9M7 | leucine rich repeat containing 59 | LRRC59 | 2.2 | 3.40 | 2.08 |
| Q04967 | heat shock protein family A (Hsp70) member 6 | HSPA6 | 1.5 | 3.39 | 2.08 |
| E1CAJ5 | Protein disulfide-isomerase | grp-58 | 1.4 | 3.39 | 2.08 |
| G8FUN5 | Y-box binding protein 1 | YBX1 | 1.8 | 3.39 | 2.08 |
| I3LPS3 | tetratricopeptide repeat domain 19 | TTC19 | 10.4 | 3.38 | 2.07 |
| A0A287A8T0 | ribosomal protein L7 | RPL7 | 2.7 | 3.38 | 2.07 |
| F1SS34 | phospholipase C beta 2 | PLCB2 | -13.8 | 3.37 | 2.06 |
| I3LMC8 | NOP16 nucleolar protein | NOP16 | 3.3 | 3.36 | 2.06 |
| K7GKC0 | ribosomal protein S16 | RPS16 | 1.5 | 3.35 | 2.05 |
| A0A287BJL8 | procollagen-lysine,2-oxoglutarate 5-dioxygenase 2 | PLOD2 | 14.6 | 3.33 | 2.03 |
| A0A286ZJQ4 | microtubule associated protein 1B | MAP1B | 2.0 | 3.34 | 2.03 |
| I3LL27 | vacuolar protein sorting 4 homolog A | VPS4A | 1.8 | 3.34 | 2.03 |
| F1SI16 | bone morphogenetic protein receptor type 2 | BMPR2 | -14.5 | 3.32 | 2.02 |
| H2F098 | macrophage receptor with collagenous structure | MARCO | -17.9 | 3.30 | 2.01 |
| C3VML1 | claudin 5 | CLDN5 | -3.5 | 3.30 | 2.00 |
| F2Z4Y0 | small nuclear ribonucleoprotein D3 polypeptide | SNRPD3 | 2.6 | 3.30 | 2.00 |

| | | | | | |
|---|---|---|---|---|---|
| A0A287AB21 | protein phosphatase 4 regulatory subunit 1 | PPP4R1 | 9.9 | 3.28 | 1.98 |
| A0A287AR45 | stabilin 1 | STAB1 | 2.1 | 3.27 | 1.98 |
| F1RTV5 | phosphoribosylaminoimidazole carboxylase and phosphoribosylaminoimidazolesuccinocarboxamide synthase | PAICS | 1.5 | 3.27 | 1.98 |
| A0A286ZWI2 | nudix hydrolase 21 | NUDT21 | 1.0 | 3.27 | 1.98 |
| A0A287A3T1 | Toll-interacting protein | Tollip | 0.7 | 3.27 | 1.98 |
| A0A287ATE4 | poly(A) binding protein nuclear 1 | PABPN1 | 1.3 | 3.26 | 1.97 |
| I3LC84 | sulfotransferase family 1C member 4 | SULT1C4 | -2.2 | 3.25 | 1.97 |
| A0A287B1T2 | EH domain containing 3 | EHD3 | -3.0 | 3.25 | 1.96 |
| K9IVK7 | YY1 transcription factor | YY1 | 1.8 | 3.24 | 1.95 |
| Q68RU1 | Ovarian and testicular apolipoprotein N | ApoN | -4.2 | 3.23 | 1.95 |
| A0A287ACE3 | SH3 domain binding glutamate rich protein like 2 | SH3BGRL2 | -1.7 | 3.22 | 1.94 |
| F1SDT0 | junctophilin 2 | JPH2 | 12.2 | 3.22 | 1.94 |
| B7TY21 | Platelet glycoprotein Ib beta | GPIbB | -6.5 | 3.20 | 1.93 |
| A0A287BMU2 | sarcolemma associated protein | SLMAP | 1.1 | 3.20 | 1.92 |
| A0A287B278 | annexin A4 | ANXA4 | 0.7 | 3.19 | 1.92 |
| A0A287AWR7 | paraoxonase 1 | PON1 | -3.8 | 3.19 | 1.92 |
| Q69DK8 | complement C1s | C1S | -4.4 | 3.19 | 1.92 |
| K7ZRK0 | IgA heavy chian constant region | IGHA | -1.2 | 3.19 | 1.92 |
| K7GQI7 | Apoptosis-associated speck-like protein containing a CARD | ASC | -1.9 | 3.18 | 1.91 |
| F2Z561 | neurocalcin delta | NCALD | -1.8 | 3.16 | 1.90 |
| F1SHK2 | MAP3K12 binding inhibitory protein 1 | MBIP | 12.1 | 3.15 | 1.88 |
| F1SII4 | Diadenosine tetraphosphate synthetase | GARS | 1.1 | 3.15 | 1.88 |
| F1SGS2 | Peptidase S1 domain-containing protein | LOC100739080 | -5.7 | 3.14 | 1.88 |
| A0A287B8E3 | 3-hydroxy-3-methylglutaryl-CoA reductase | HMGCR | -17.5 | 3.14 | 1.88 |
| A0A287AYD5 | microtubule associated protein 1A | MAP1A | 2.0 | 3.12 | 1.86 |
| I3LCJ3 | 5'-nucleotidase, cytosolic IIIA | NT5C3A | -8.8 | 3.12 | 1.86 |
| B2ZF47 | aldehyde dehydrogenase 2 family member | ALDH2 | 0.7 | 3.11 | 1.86 |
| F1SNF3 | podocalyxin like | PODXL | -2.6 | 3.11 | 1.86 |
| F1SCV9 | apolipoprotein B | APOB | -2.4 | 3.11 | 1.85 |
| K7GK90 | signal sequence receptor subunit 4 | SSR4 | 0.7 | 3.10 | 1.85 |
| F1RKQ8 | cleavage and polyadenylation specific factor 7 | CPSF7 | 1.9 | 3.09 | 1.84 |
| F1S3G5 | peptidylprolyl cis/trans isomerase, NIMA-interacting 1 | PIN1 | 1.7 | 3.09 | 1.84 |
| F1SN86 | regulator of G protein signaling 3 | RGS3 | -12.5 | 3.09 | 1.84 |
| A0A287AQ20 | complement factor I | CFI | -2.2 | 3.08 | 1.83 |
| F1S1F3 | cytochrome c oxidase assembly factor 3 | COA3 | 13.3 | 3.08 | 1.83 |
| I3LRT6 | FKBP prolyl isomerase 9 | FKBP9 | 1.9 | 3.08 | 1.83 |
| A0A286ZQ63 | SEC14 like lipid binding 3 | SEC14L3 | -1.7 | 3.08 | 1.83 |
| Q9XSI4 | hypoxanthine phosphoribosyltransferase 1 | HPRT1 | -1.9 | 3.06 | 1.82 |
| Q05KQ7 | Chemokine-like receptor 1 | gpcr | -6.7 | 3.06 | 1.82 |
| A0A287BIK6 | pseudopodium enriched atypical kinase 1 | PEAK1 | 10.6 | 3.05 | 1.81 |
| K7GQE8 | ATP-binding cassette sub-family G member 2 | LOC110255210 | -11.8 | 3.05 | 1.81 |
| F1RHG4 | methyltransferase like 16 | METTL16 | 7.3 | 3.04 | 1.80 |
| P04366 | alpha-1-microglobulin/bikunin precursor | AMBP | -2.8 | 3.04 | 1.80 |
| A0A287B2V5 | double PHD fingers 2 | DPF2 | 1.7 | 3.02 | 1.78 |

| | | | | | |
|---|---|---|---|---|---|
| F1RSN3 | 5-oxoprolinase, ATP-hydrolysing | OPLAH | -4.1 | 3.02 | 1.78 |
| I3LFP3 | versican | VCAN | 2.2 | 3.02 | 1.78 |
| A0A2C9F361 | molybdenum cofactor synthesis 3 | MOCS3 | -6.3 | 3.02 | 1.78 |
| F1SFI4 | kininogen 1 | KNG1 | -3.4 | 3.01 | 1.78 |
| F1S771 | tetratricopeptide repeat domain 4 | TTC4 | 10.7 | 3.01 | 1.78 |
| D0G7F6 | triosephosphate isomerase 1 | TPI1 | 1.1 | 2.99 | 1.76 |
| I3L650 | caldesmon 1 | CALD1 | 1.1 | 2.99 | 1.76 |
| Q32YV9 | proteasome 26S subunit, non-ATPase 4 | PSMD4 | 0.8 | 2.98 | 1.76 |
| A0A287BFC6 | glycosylphosphatidylinositol anchored high density lipoprotein binding protein 1 | GPIHBP1 | -4.5 | 2.98 | 1.75 |
| A0A287AUL3 | cortactin | CTTN | 1.6 | 2.96 | 1.74 |
| P62802 | Histone H4 | H4C1 | 1.0 | 2.96 | 1.74 |
| P33198 | isocitrate dehydrogenase (NADP(+)) 2 | IDH2 | 0.8 | 2.96 | 1.74 |
| F2Z5E2 | serpin family C member 1 | SERPINC1 | -2.0 | 2.96 | 1.74 |
| F1SD69 | legumain | LGMN | 2.7 | 2.96 | 1.74 |
| A0A287B9G3 | periaxin | PRX | -10.8 | 2.95 | 1.73 |
| A0A287A7Q7 | eukaryotic translation initiation factor 5 | EIF5 | 1.2 | 2.95 | 1.73 |
| A0A287ADW0 | FAM3 metabolism regulating signaling molecule C | FAM3C | 1.6 | 2.94 | 1.73 |
| A0A286ZQE6 | Kinesin light chain | KLC1 | 1.6 | 2.94 | 1.72 |
| F1RXC2 | carbonic anhydrase 2 | CA2 | -1.9 | 2.93 | 1.72 |
| I3LS74 | Antileukoproteinase | LOC100512873 | -4.0 | 2.93 | 1.72 |
| Q29092 | heat shock protein 90 beta family member 1 | HSP90B1 | 1.4 | 2.93 | 1.72 |
| A0A287B150 | LLGL scribble cell polarity complex component 2 | LLGL2 | -2.3 | 2.93 | 1.72 |
| A4H2R5 | secreted protein acidic and cysteine rich | SPARC | 2.2 | 2.92 | 1.71 |
| F1STR6 | nuclear distribution C, dynein complex regulator | NUDC | 0.7 | 2.92 | 1.71 |
| A0A287BJG3 | 3'-phosphoadenosine 5'-phosphosulfate synthase 1 | PAPSS1 | 1.8 | 2.92 | 1.71 |
| F1S766 | mago homolog, exon junction complex subunit | MAGOH | 1.4 | 2.92 | 1.71 |
| F1S1X3 | Asparagine--tRNA ligase | NARS | 1.2 | 2.92 | 1.71 |
| A5A8W8 | C4a anaphylatoxin | C4A | -2.3 | 2.91 | 1.71 |
| P81649 | ribonuclease A family member k6 | RNASE6 | -13.0 | 2.90 | 1.70 |
| F1RT83 | syndecan binding protein | SDCBP | 0.8 | 2.90 | 1.70 |
| F1SPE9 | DnaJ heat shock protein family (Hsp40) member C13 | DNAJC13 | -2.4 | 2.90 | 1.70 |
| I3LBG5 | regulator of MON1-CCZ1 | RMC1 | -6.1 | 2.89 | 1.69 |
| B6ECP2 | Platelet glycoprotein Ib alpha polypeptide | GPIbA | -12.9 | 2.89 | 1.69 |
| F1SUN0 | arrestin beta 1 | ARRB1 | -0.5 | 2.89 | 1.69 |
| A0A0U2ETD0 | emerin | EMD | 1.1 | 2.89 | 1.69 |
| A0A287BQW1 | methionyl aminopeptidase 1 | METAP1 | 1.0 | 2.88 | 1.69 |
| K9J6J9 | RNA polymerase II associated protein 3 | RPAP3 | 11.3 | 2.88 | 1.69 |
| I3LUP6 | nucleophosmin 1 | NPM1 | 1.1 | 2.88 | 1.69 |
| P26234 | vinculin | VCL | 0.7 | 2.88 | 1.69 |
| B6EAV5 | SLC9A3 regulator 2 | SLC9A3R2 | -1.1 | 2.88 | 1.69 |
| I3LIY2 | inositol polyphosphate-5-phosphatase K | INPP5K | -3.4 | 2.88 | 1.69 |
| F1S3P6 | CCN family member 2 | CTGF | -7.3 | 2.88 | 1.69 |
| A0A287BTI9 | TUB like protein 3 | TULP3 | 14.1 | 2.87 | 1.69 |
| A0A286ZRU9 | serpin family H member 1 | SERPINH1 | 2.3 | 2.87 | 1.68 |
| A0A287ACY7 | proteasome 26S subunit, non-ATPase 10 | PSMD10 | 1.2 | 2.87 | 1.68 |

| | | | | | |
|---|---|---|---|---|---|
| A0A287A6X1 | TRAF3 interacting protein 3 | TRAF3IP3 | -3.4 | 2.87 | 1.68 |
| F1S4E4 | leucine rich repeat containing 8 VRAC subunit C | LRRC8C | -7.4 | 2.86 | 1.67 |
| A0A287BNT2 | junctional adhesion molecule 2 | JAM2 | -1.6 | 2.85 | 1.67 |
| F1SS00 | transmembrane BAX inhibitor motif containing 1 | TMBIM1 | -1.0 | 2.85 | 1.67 |
| I3LBR0 | tolloid like 1 | TLL1 | 18.5 | 2.85 | 1.67 |
| F1SKM1 | collagen type VII alpha 1 chain | COL7A1 | -8.9 | 2.84 | 1.66 |
| F1SP18 | Threonyl-tRNA synthetase | TARS | 1.1 | 2.84 | 1.66 |
| I3LIM2 | UDP-glucose 6-dehydrogenase | UGDH | 1.4 | 2.83 | 1.66 |
| K7GNZ3 | nascent polypeptide associated complex subunit alpha | NACA | 1.3 | 2.83 | 1.66 |
| A0A286ZXZ1 | EMAP like 2 | EML2 | -1.4 | 2.83 | 1.66 |
| K7N7E9 | BUD31 homolog | BUD31 | 0.9 | 2.83 | 1.66 |
| A0A287BF06 | ribosomal protein S27 | RPS27 | 2.0 | 2.83 | 1.66 |
| A0A287A329 | myotubularin 1 | MTM1 | -7.3 | 2.83 | 1.66 |
| I3LRP1 | nuclear autoantigenic sperm protein | NASP | 1.6 | 2.82 | 1.65 |
| F1SA47 | myosin IF | MYO1F | -4.5 | 2.81 | 1.65 |
| F1RND9 | myeloid cell nuclear differentiation antigen | MNDA | -7.6 | 2.81 | 1.65 |
| F1SHI0 | 1-aminocyclopropane-1-carboxylate synthase homolog (inactive) | ACCS | 12.9 | 2.81 | 1.64 |
| I3LEB7 | podocan | PODN | 1.1 | 2.80 | 1.64 |
| F2Z5S5 | biogenesis of lysosomal organelles complex 1 subunit 1 | BLOC1S1 | 13.3 | 2.80 | 1.64 |
| I3LPL9 | makorin ring finger protein 2 | MKRN2 | 12.4 | 2.80 | 1.64 |
| F1S155 | SEC24 homolog D, COPII coat complex component | SEC24D | 1.5 | 2.80 | 1.64 |
| F1SAA5 | ubiquitin C-terminal hydrolase L5 | UCHL5 | 1.2 | 2.80 | 1.64 |
| C3S7K4 | S100 calcium binding protein A12 | S100A12 | -2.0 | 2.80 | 1.64 |
| A0A287AIT2 | reticulon 4 | RTN4 | -5.5 | 2.80 | 1.64 |
| A0A287AL85 | GA binding protein transcription factor subunit alpha | GABPA | 1.9 | 2.79 | 1.64 |
| F1SCY2 | interferon induced protein with tetratricopeptide repeats 3 | IFIT3 | -2.3 | 2.79 | 1.63 |
| I3LGC2 | ribosome binding protein 1 | RRBP1 | 2.6 | 2.79 | 1.63 |
| A0A287B0C2 | G protein-coupled receptor class C group 5 member A | GPRC5A | -1.7 | 2.79 | 1.63 |
| F1S8N1 | HGF activator | HGFAC | -5.3 | 2.78 | 1.63 |
| F1RG45 | angiotensinogen | AGT | -1.9 | 2.78 | 1.63 |
| F1SEQ6 | tetraspanin 14 | TSPAN14 | -3.4 | 2.78 | 1.63 |
| F1RK83 | scavenger receptor class F member 2 | SCARF2 | 12.7 | 2.78 | 1.63 |
| A0A287AT23 | aquaporin 1 (Colton blood group) | AQP1 | -1.0 | 2.78 | 1.63 |
| F1RS49 | ATP binding cassette subfamily E member 1 | ABCE1 | 2.6 | 2.77 | 1.62 |
| F1RNP4 | purinergic receptor P2X 7 | P2RX7 | 12.9 | 2.76 | 1.61 |
| A0A287BI63 | RAB6A, member RAS oncogene family | RAB6A | 0.5 | 2.76 | 1.61 |
| A0A287AII3 | TSC22 domain family member 1 | TSC22D1 | 9.5 | 2.76 | 1.61 |
| A0A287A4H2 | purine rich element binding protein B | PURB | 1.1 | 2.75 | 1.61 |
| B2NJ26 | Beta-2-microglobulin | B2m | -1.2 | 2.75 | 1.61 |
| F1RU31 | spliceosome associated factor 1, recruiter of U4/U6.U5 tri-snRNP | SART1 | 1.8 | 2.75 | 1.61 |
| A0A287BMC7 | C1q and TNF related 1 | C1QTNF1 | 14.8 | 2.75 | 1.61 |
| F1RM56 | RNA binding motif protein 42 | RBM42 | 9.9 | 2.75 | 1.61 |
| B9TRW9 | G protein subunit gamma 12 | GNG12 | -0.9 | 2.75 | 1.61 |
| F1RJ89 | regulator of cell cycle | RGCC | 2.4 | 2.74 | 1.60 |

| | | | | | |
|---|---|---|---|---|---|
| A0A287AQ40 | ZFP91 zinc finger protein, atypical E3 ubiquitin ligase | ZFP91 | 13.8 | 2.74 | 1.60 |
| A0A287A317 | hyaluronidase 2 | HYAL2 | -0.9 | 2.73 | 1.59 |
| A0A287AY64 | GLI pathogenesis related 2 | GLIPR2 | 1.5 | 2.73 | 1.59 |
| F1RU23 | cathepsin W | CTSW | -1.8 | 2.72 | 1.59 |
| A0A287BEA7 | coagulation factor XIII B chain | F13B | -13.2 | 2.73 | 1.59 |
| Q9XSZ6 | Fc fragment of IgE receptor Ig | FCER1G | -1.6 | 2.72 | 1.59 |
| Q06AU4 | RAB34, member RAS oncogene family | RAB34 | 1.7 | 2.72 | 1.59 |
| A0A287AVN5 | C-X9-C motif containing 1 | CMC1 | 11.4 | 2.72 | 1.58 |
| M3UZ93 | synaptogyrin 2 | SYNGR2 | -1.0 | 2.71 | 1.58 |
| A0A287A6Q1 | coiled-coil domain containing 134 | CCDC134 | 11.3 | 2.71 | 1.58 |
| F1RXR3 | ALG12 alpha-1,6-mannosyltransferase | ALG12 | -7.6 | 2.71 | 1.58 |
| A0A287AXU0 | elastin | ELN | 11.8 | 2.70 | 1.57 |
| F1RJR9 | PDZ binding kinase | PBK | 13.2 | 2.69 | 1.57 |
| F2Z5L7 | proteasome 20S subunit alpha 1 | PSMA1 | 0.5 | 2.69 | 1.57 |
| F1RLL9 | collagen type IV alpha 2 chain | COL4A2 | 1.8 | 2.69 | 1.56 |
| F1RWZ1 | PX domain containing 1 | PXDC1 | 11.5 | 2.69 | 1.56 |
| F1RZ28 | ribosomal protein S10 | RPS10 | 1.1 | 2.68 | 1.56 |
| F1SMZ7 | heat shock protein family D (Hsp60) member 1 | HSPD1 | 1.0 | 2.68 | 1.56 |
| Q2LE37 | apolipoprotein M | APOM | -6.4 | 2.68 | 1.56 |
| A0A287A7L4 | leucine rich repeat containing 25 | LRRC25 | -8.8 | 2.68 | 1.56 |
| F1SR51 | NCK associated protein 1 like | NCKAP1L | -8.3 | 2.67 | 1.55 |
| A0A287AFY6 | dedicator of cytokinesis 2 | DOCK2 | -5.8 | 2.67 | 1.55 |
| I3LL77 | chromosome 6 open reading frame 120 | C6orf120 | 11.3 | 2.66 | 1.55 |
| I3LQH7 | biliverdin reductase B | BLVRB | -2.0 | 2.66 | 1.55 |
| F1SIB1 | coagulation factor II, thrombin | F2 | -2.6 | 2.66 | 1.54 |
| A0A287BJ41 | tropomodulin 1 | TMOD1 | -0.9 | 2.66 | 1.54 |
| I3LVD5 | actin gamma 1 | ACTG1 | 1.1 | 2.65 | 1.54 |
| I3LKI1 | serglycin | SRGN | -6.0 | 2.65 | 1.54 |
| I3LIP6 | aspartate beta-hydroxylase | ASPH | 1.2 | 2.64 | 1.52 |
| F1RGI9 | SIL1 nucleotide exchange factor | SIL1 | 7.3 | 2.63 | 1.52 |
| I3LSD3 | ribosomal protein L13 | RPL13 | 2.2 | 2.62 | 1.51 |
| I3LQ17 | PZP alpha-2-macroglobulin like | PZP | -2.8 | 2.61 | 1.51 |
| I3L5U9 | hook microtubule tethering protein 3 | HOOK3 | 1.7 | 2.61 | 1.50 |
| A0A287BAF3 | Adenylosuccinate synthetase isozyme 1 | ADSS1 | 3.0 | 2.61 | 1.50 |
| K7GS36 | neuropilin 1 | NRP1 | -0.7 | 2.60 | 1.50 |
| F1RKR4 | transmembrane p24 trafficking protein 3 | TMED3 | 1.2 | 2.60 | 1.50 |
| O77633 | ADAM metallopeptidase domain 10 | ADAM10 | -0.5 | 2.60 | 1.50 |
| Q1T7A8 | collagen type VI alpha 1 chain | COL6A1 | -1.5 | 2.60 | 1.50 |
| I3LP02 | acetyl-CoA acetyltransferase 1 | ACAT1 | 1.1 | 2.59 | 1.49 |
| A0A287ADK3 | aldehyde dehydrogenase 7 family member A1 | ALDH7A1 | 1.5 | 2.59 | 1.49 |
| A0A287AKZ5 | eukaryotic translation initiation factor 1A X-linked | EIF1AX | 0.7 | 2.59 | 1.49 |
| I3LCH3 | RNA polymerase II subunit C | POLR2C | 1.0 | 2.59 | 1.49 |
| A5GFX4 | ATP synthase, H+ transporting, mitochondrial F1 complex, epsilon su | ATP5E | -0.9 | 2.58 | 1.48 |
| K9IVI0 | solute carrier family 43 member 3 | SLC43A3 | -12.8 | 2.58 | 1.48 |
| F1SM61 | fibulin 1 | FBLN1 | -1.0 | 2.58 | 1.48 |

| | | | | | |
|---|---|---|---|---|---|
| F1RYS9 | starch binding domain 1 | STBD1 | 10.4 | 2.57 | 1.48 |
| F2Z4Y8 | ribosomal protein S11 | RPS11 | 0.9 | 2.56 | 1.47 |
| Q6UAQ8 | electron transfer flavoprotein subunit beta | ETFB | 0.9 | 2.56 | 1.47 |
| F1SNU2 | elongator acetyltransferase complex subunit 6 | ELP6 | 9.0 | 2.56 | 1.47 |
| A0A287AZY6 | TNF alpha induced protein 8 like 1 | TNFAIP8L1 | 8.5 | 2.56 | 1.47 |
| I3LV38 | methionine sulfoxide reductase B3 | MSRB3 | 2.2 | 2.56 | 1.47 |
| F1RNZ1 | Rieske domain-containing protein | LOC100522678 | 1.3 | 2.56 | 1.47 |
| I3LDW9 | small nuclear ribonucleoprotein polypeptide E | SNRPE | 1.1 | 2.56 | 1.47 |
| F1S519 | kinesin family member 3B | KIF3B | -6.5 | 2.55 | 1.47 |
| F1SBS4 | complement C3 | C3 | -2.3 | 2.55 | 1.46 |
| F1SAX5 | CD58 molecule | CD58 | -1.3 | 2.55 | 1.46 |
| M3UZ96 | uridine-cytidine kinase 1 like 1 | UCKL1 | 10.7 | 2.54 | 1.46 |
| A0A287A1X0 | fibronectin type III domain containing 3B | FNDC3B | 10.1 | 2.54 | 1.46 |
| F1SHL0 | C-type lectin domain containing 14A | CLEC14A | -0.9 | 2.54 | 1.46 |
| A0A287AFA3 | Zinc finger, CCHC domain containing 6, isoform CRA_b | ZCCHC6 | -2.1 | 2.53 | 1.45 |
| A0A287BHL7 | extracellular matrix protein 1 | ECM1 | -7.3 | 2.52 | 1.44 |
| A0A2C9F3H8 | BAG cochaperone 6 | BAG6 | 1.8 | 2.52 | 1.44 |
| F2Z5V6 | Myosin regulatory light chain 12A isoform 2 | LOC733637 | 1.2 | 2.52 | 1.44 |
| A0A287BRC0 | cytoplasmic FMR1 interacting protein 2 | CYFIP2 | -4.7 | 2.52 | 1.44 |
| F1SFF4 | RNA transcription, translation and transport factor | RTRAF | 1.2 | 2.51 | 1.43 |
| I3L651 | fibrinogen beta chain | FGB | -3.1 | 2.51 | 1.43 |
| A0A287BSA7 | COPI coat complex subunit zeta 2 | COPZ2 | 0.9 | 2.51 | 1.43 |
| A0A287B4P8 | proteasome 26S subunit, ATPase 4 | PSMC4 | 0.6 | 2.51 | 1.43 |
| A0A286ZY10 | glucose-6-phosphate dehydrogenase | G6PD | -1.2 | 2.51 | 1.43 |
| A0A286ZP79 | src kinase associated phosphoprotein 2 | SKAP2 | -13.4 | 2.51 | 1.43 |
| Q29198 | 40S ribosomal protein S6 | RPS6 | 1.9 | 2.50 | 1.43 |
| F1SJV8 | RNA exonuclease 2 | REXO2 | 2.0 | 2.50 | 1.43 |
| P09571 | transferrin | TF | -1.6 | 2.50 | 1.42 |
| F1SFI6 | fetuin B | FETUB | -2.9 | 2.50 | 1.42 |
| K9IW73 | Dihydrolipoyllysine-residue succinyltransferase component of 2-oxoglutarate dehydrogenase complex, mitochondrial | DLST-tv1 | 0.7 | 2.49 | 1.42 |
| A0A286ZSJ7 | complement C1q C chain | C1QC | -3.2 | 2.49 | 1.42 |
| A9YTX9 | interferon regulatory factor 3 | IRF3 | -1.0 | 2.49 | 1.42 |
| F1RMD4 | thrombospondin type 1 domain containing 1 | THSD1 | -11.2 | 2.49 | 1.42 |
| F1RFI8 | EWS RNA binding protein 1 | EWSR1 | 1.2 | 2.48 | 1.41 |
| A0A287ABK7 | ATP binding cassette subfamily C member 1 | ABCC1 | -7.9 | 2.48 | 1.41 |
| F1SSY0 | TNF alpha induced protein 8 like 2 | TNFAIP8L2 | -2.3 | 2.48 | 1.41 |
| O97506 | kallikrein B1 | KLKB1 | -3.1 | 2.47 | 1.41 |
| A0A287A042 | maltase-glucoamylase | MGAM | -3.8 | 2.47 | 1.41 |
| A0A287A502 | laminin subunit alpha 2 | LAMA2 | -9.9 | 2.47 | 1.41 |
| M3VK30 | nucleolar protein 7 | NOL7 | 4.4 | 2.47 | 1.41 |
| Q95309 | TEGT protein | TEGT | -4.0 | 2.47 | 1.40 |
| I3LU51 | heterogeneous nuclear ribonucleoprotein U like 1 | HNRNPUL1 | 1.3 | 2.46 | 1.40 |
| F1RL04 | protein phosphatase, Mg2+/Mn2+ dependent 1F | PPM1F | -1.0 | 2.46 | 1.40 |
| Q9GK83 | allograft inflammatory factor 1 | AIF1 | -2.5 | 2.46 | 1.40 |

| | | | | | |
|---|---|---|---|---|---|
| A0A287BM51 | ubiquitin family domain containing 1 | UBFD1 | 1.0 | 2.45 | 1.39 |
| A0A286ZXK8 | malonyl-CoA-acyl carrier protein transacylase | MCAT | 6.4 | 2.44 | 1.38 |
| A0A287AAW7 | C1q domain-containing protein | LOC110258312 | -3.1 | 2.44 | 1.38 |
| A7E1T5 | high mobility group box 1 | HMGB1 | 1.7 | 2.43 | 1.37 |
| I3LJC8 | retention in endoplasmic reticulum sorting receptor 1 | RER1 | -5.3 | 2.42 | 1.37 |
| A5H025 | ribonuclease L | RNASEL | -13.7 | 2.42 | 1.37 |
| I3LP21 | potassium voltage-gated channel subfamily A regulatory beta subunit 2 | KCNAB2 | 1.6 | 2.42 | 1.37 |
| I3LNR4 | transmembrane protein 91 | TMEM91 | 1.3 | 2.42 | 1.37 |
| A0A287AUM3 | cysteine rich transmembrane module containing 1 | CYSTM1 | -2.1 | 2.42 | 1.37 |
| A0A287AQP4 | phosphatidylinositol 4-kinase alpha | PI4KA | -4.1 | 2.42 | 1.36 |
| K7GK75 | cofilin 1 | CFL1 | 0.6 | 2.41 | 1.36 |
| F1SAT9 | thrombomodulin | THBD | -1.9 | 2.41 | 1.36 |
| F1S8W9 | pyridine nucleotide-disulphide oxidoreductase domain 2 | PYROXD2 | -1.9 | 2.41 | 1.36 |
| A0SEH0 | complement C6 | C6 | -5.1 | 2.41 | 1.36 |
| A0A286ZJQ9 | caveolin 1 | CAV1 | -0.8 | 2.40 | 1.36 |
| A0A287A4Q7 | actinin alpha 4 | ACTN4 | 1.3 | 2.40 | 1.35 |
| A0A287AG37 | G3BP stress granule assembly factor 2 | G3BP2 | 1.3 | 2.40 | 1.35 |
| F1SI48 | erythrocyte membrane protein band 4.2 | EPB42 | -5.6 | 2.40 | 1.35 |
| F1SLI6 | SWI/SNF related, matrix associated, actin dependent regulator of chromatin subfamily c member 1 | SMARCC1 | 9.1 | 2.40 | 1.35 |
| A0A287A9H4 | heterogeneous nuclear ribonucleoprotein F | HNRNPF | 1.2 | 2.40 | 1.35 |
| F1SC80 | retinol binding protein 4 | RBP4 | -2.3 | 2.40 | 1.35 |
| A0A286ZP86 | TNF receptor associated factor 7 | TRAF7 | -5.8 | 2.40 | 1.35 |
| P67985 | ribosomal protein L22 | RPL22 | 1.6 | 2.40 | 1.35 |
| C0JX97 | Intercellular adhesion molecule-3 | ICAM-3 | -9.1 | 2.39 | 1.35 |
| A0A286ZWW2 | heterogeneous nuclear ribonucleoprotein H2 | HNRNPH2 | 2.4 | 2.39 | 1.35 |
| P28491 | calreticulin | CALR | 1.3 | 2.39 | 1.35 |
| A0A286ZY30 | KH-type splicing regulatory protein | KHSRP | 1.2 | 2.39 | 1.35 |
| A0A287BQU7 | small nuclear ribonucleoprotein D2 polypeptide | SNRPD2 | 0.7 | 2.38 | 1.35 |
| I3LRY7 | DNA methyltransferase 3 alpha | DNMT3A | 9.2 | 2.38 | 1.35 |
| A0A287AZG0 | Rho GTPase activating protein 1 | ARHGAP1 | 0.6 | 2.38 | 1.35 |
| I3LNB4 | aldehyde dehydrogenase 4 family member A1 | ALDH4A1 | 1.2 | 2.38 | 1.35 |
| F1REX8 | huntingtin interacting protein 1 related | HIP1R | -2.5 | 2.38 | 1.34 |
| Q08094 | calponin 2 | CNN2 | 1.1 | 2.38 | 1.34 |
| F1RN76 | CD5 molecule like | CD5L | -2.9 | 2.37 | 1.34 |
| A0A287BQ61 | RUN and FYVE domain containing 3 | RUFY3 | -5.3 | 2.37 | 1.33 |
| A5GFX6 | tubulin beta 1 class VI | TUBB1 | -12.7 | 2.37 | 1.33 |
| A0A287BLN5 | mitochondrial ribosomal protein L9 | MRPL9 | 8.9 | 2.36 | 1.33 |
| A0A287AYX5 | N-acetylneuraminic acid phosphatase | NANP | 8.3 | 2.36 | 1.33 |
| A0A286ZK74 | heterogeneous nuclear ribonucleoprotein C | HNRNPC | 1.1 | 2.36 | 1.33 |
| A0A287BG35 | ring finger protein 170 | RNF170 | -2.3 | 2.36 | 1.33 |
| F2Z5F1 | splicing factor 3b subunit 6 | SF3B6 | 0.8 | 2.35 | 1.33 |
| I3LHW6 | receptor activity modifying protein 3 | RAMP3 | -12.2 | 2.35 | 1.33 |
| A0A287BI02 | collagen type IV alpha 4 chain | COL4A4 | -7.3 | 2.35 | 1.32 |
| I3LF89 | carboxypeptidase N subunit 2 | CPN2 | -4.2 | 2.35 | 1.32 |
| A0A286ZSL2 | allograft inflammatory factor 1 like | AIF1L | 11.3 | 2.34 | 1.32 |

| | | | | | |
|---|---|---|---|---|---|
| F2Z5K5 | tubulin beta 4A class IVa | TUBB4A | 11.2 | 2.34 | 1.32 |
| Q29361 | ribosomal protein L35 | RPL35 | 4.4 | 2.34 | 1.32 |
| F1S431 | Alanine--tRNA ligase | AARS1 | 1.0 | 2.34 | 1.32 |
| F1RUS8 | tubulin folding cofactor C | TBCC | 0.8 | 2.34 | 1.32 |
| F1SJU4 | parvin beta | PARVB | -8.3 | 2.33 | 1.31 |
| F1RGD9 | Histidine--tRNA ligase, cytoplasmic | HARS | 2.0 | 2.33 | 1.31 |
| K7GR37 | Septin-6 | SPTAN6 | -0.9 | 2.33 | 1.31 |
| L8B0S7 | IgG heavy chain | IGHG | -11.4 | 2.33 | 1.31 |
| F1SL58 | dynactin subunit 2 | DCTN2 | 0.8 | 2.32 | 1.30 |
| I3LKV0 | SRA stem-loop interacting RNA binding protein | SLIRP | 1.3 | 2.32 | 1.30 |

**Supplemental Table 2. Significantly Altered Ingenuity® Canonical Pathways at Uncorrected P < 0.05 Level.**

| PVH Artery vs. Sham Artery: Ingenuity Pathways with Uncorrected P < 0.05 ($-log_{10}$ P > 1.3) | | | | | | |
|---|---|---|---|---|---|---|
| | Pulmonary Artery | | Pulmonary Vein | | Whole Lung | |
| Ingenuity Canonical Pathways | $-Log_{10}$ (P-value) | $-Log_{10}$ (FDR P-value) | $-Log_{10}$ (P-value) | $-Log_{10}$ (FDR P-value) | $-Log_{10}$ (P-value) | $-Log_{10}$ (FDR P-value) |
| EIF2 Signaling | 48.10 | 45.60 | 7.68 | 5.41 | 9.83 | 7.91 |
| Regulation of eIF4 and p70S6K Signaling | 13.80 | 11.70 | 5.38 | 3.43 | 4.02 | 2.51 |
| mTOR Signaling | 11.90 | 9.87 | 4.90 | 3.16 | 3.30 | 2.05 |
| Protein Ubiquitination Pathway | 3.21 | 1.39 | 0.91 | 0.55 | 2.24 | 1.26 |
| Arginine Biosynthesis IV | 3.16 | 1.39 | | | | |
| tRNA Charging | 2.62 | 1.05 | 1.91 | 1.12 | 5.52 | 3.85 |
| Agrin Interactions at Neuromuscular Junction | 2.67 | 1.05 | 1.78 | 1.04 | 1.96 | 1.07 |
| Inhibition of Matrix Metalloproteases | 2.62 | 1.05 | | | 1.88 | 1.02 |
| Glycogen Degradation II | 2.53 | 1.01 | 0.86 | 0.53 | 0.60 | 0.26 |
| α-Adrenergic Signaling | 2.36 | 0.94 | 2.17 | 1.27 | 0.00 | 0.00 |
| Glycogen Degradation III | 2.39 | 0.94 | 0.80 | 0.49 | 1.38 | 0.65 |
| Superpathway of Citrulline Metabolism | 2.33 | 0.94 | 0.77 | 0.47 | 0.52 | 0.22 |
| Virus Entry via Endocytic Pathways | 2.19 | 0.88 | 1.98 | 1.14 | 0.96 | 0.44 |
| CDK5 Signaling | 2.18 | 0.88 | 1.35 | 0.82 | 0.00 | 0.00 |
| Unfolded protein response | 2.17 | 0.88 | 0.30 | 0.18 | 4.31 | 2.77 |
| Colorectal Cancer Metastasis Signaling | 2.09 | 0.82 | 1.03 | 0.64 | 0.26 | 0.08 |
| GP6 Signaling Pathway | 2.03 | 0.80 | 1.79 | 1.04 | 7.66 | 5.88 |
| G Beta Gamma Signaling | 2.00 | 0.78 | 3.09 | 1.86 | 0.79 | 0.35 |
| Caveolar-mediated Endocytosis Signaling | 1.86 | 0.77 | 7.66 | 5.41 | 1.02 | 0.49 |
| Ephrin B Signaling | 1.87 | 0.77 | 4.52 | 2.93 | 2.15 | 1.19 |
| IL-8 Signaling | 1.90 | 0.77 | 2.46 | 1.46 | 2.08 | 1.17 |
| Integrin Signaling | 1.80 | 0.77 | 2.29 | 1.36 | 1.51 | 0.76 |
| Gαi Signaling | 1.96 | 0.77 | 1.71 | 1.01 | 0.00 | 0.00 |
| Glioma Invasiveness Signaling | 1.84 | 0.77 | 1.20 | 0.74 | 0.59 | 0.26 |
| Macropinocytosis Signaling | 1.81 | 0.77 | 0.62 | 0.39 | 0.27 | 0.08 |
| Antiproliferative Role of Somatostatin Receptor 2 | 1.79 | 0.77 | 0.21 | 0.13 | 0.00 | 0.00 |
| Glutamate Biosynthesis II | 1.86 | 0.77 | | | | |
| Glutamate Degradation X | 1.86 | 0.77 | | | | |
| Hypusine Biosynthesis | 1.69 | 0.71 | 1.44 | 0.88 | | |
| S-adenosyl-L-methionine Biosynthesis | 1.69 | 0.71 | 1.44 | 0.88 | | |
| G Protein Signaling Mediated by Tubby | 1.72 | 0.71 | 1.25 | 0.76 | 1.44 | 0.70 |
| Cardiac Hypertrophy Signaling | 1.60 | 0.65 | 2.51 | 1.48 | 0.00 | 0.00 |
| CXCR4 Signaling | 1.55 | 0.65 | 2.32 | 1.37 | 0.44 | 0.18 |
| Antigen Presentation Pathway | 1.53 | 0.65 | 1.08 | 0.67 | 0.63 | 0.27 |
| Prostate Cancer Signaling | 1.60 | 0.65 | 0.00 | 0.00 | 0.77 | 0.34 |
| Neuregulin Signaling | 1.54 | 0.65 | 0.00 | 0.00 | 0.00 | 0.00 |
| Melatonin Degradation II | 1.57 | 0.65 | | | | |
| Proline Biosynthesis I | 1.57 | 0.65 | | | | |
| Bladder Cancer Signaling | 1.53 | 0.65 | | | 0.71 | 0.31 |
| Apoptosis Signaling | 1.51 | 0.64 | 0.00 | 0.00 | | |

| | | | | | | |
|---|---|---|---|---|---|---|
| Citrulline-Nitric Oxide Cycle | 1.47 | 0.61 | 1.22 | 0.75 | 0.95 | 0.44 |
| Ephrin Receptor Signaling | 1.45 | 0.61 | 2.15 | 1.27 | 0.87 | 0.40 |
| Oncostatin M Signaling | 1.45 | 0.61 | 0.38 | 0.23 | 0.00 | 0.00 |
| UDP-N-acetyl-D-glucosamine Biosynthesis II | 1.39 | 0.58 | 2.66 | 1.54 | 0.87 | 0.40 |
| Urea Cycle | 1.39 | 0.58 | | | | |
| Leukocyte Extravasation Signaling | 1.33 | 0.55 | 3.10 | 1.86 | 3.58 | 2.23 |
| fMLP Signaling in Neutrophils | 1.34 | 0.55 | 0.76 | 0.47 | 0.53 | 0.23 |
| Natural Killer Cell Signaling | 1.33 | 0.55 | 0.65 | 0.40 | 2.13 | 1.18 |
| Thioredoxin Pathway | 1.33 | 0.55 | | | | |
| **PVH Vein vs. Sham Vein: Ingenuity Pathways with Uncorrected P < 0.05 (-$\log_{10}$ P > 1.3)** | | | | | | |
| | **Pulmonary Vein** | | **Pulmonary Artery** | | **Whole Lung** | |
| **Ingenuity Canonical Pathways** | **-Log10 (P-value)** | **-Log10 (FDR P-value)** | **-Log10 (P-value)** | **-Log10 (FDR P-value)** | **-Log10 (P-value)** | **-Log10 (FDR P-value)** |
| EIF2 Signaling | 7.68 | 5.41 | 48.10 | 45.60 | 9.83 | 7.91 |
| Caveolar-mediated Endocytosis Signaling | 7.66 | 5.41 | 1.86 | 0.77 | 1.02 | 0.49 |
| Regulation of eIF4 and p70S6K Signaling | 5.38 | 3.43 | 13.80 | 11.70 | 4.02 | 2.51 |
| Sertoli Cell-Sertoli Cell Junction Signaling | 5.46 | 3.43 | 0.44 | 0.23 | 3.86 | 2.38 |
| mTOR Signaling | 4.90 | 3.16 | 11.90 | 9.87 | 3.30 | 2.05 |
| RhoGDI Signaling | 4.81 | 3.16 | 0.90 | 0.37 | 1.19 | 0.56 |
| Signaling by Rho Family GTPases | 4.94 | 3.16 | 0.63 | 0.29 | 3.61 | 2.24 |
| Acute Phase Response Signaling | 4.84 | 3.16 | 0.00 | 0.00 | 8.75 | 6.91 |
| Ephrin B Signaling | 4.52 | 2.93 | 1.87 | 0.77 | 2.15 | 1.19 |
| Tight Junction Signaling | 4.33 | 2.78 | 0.00 | 0.00 | 1.73 | 0.92 |
| Actin Cytoskeleton Signaling | 4.05 | 2.54 | 0.72 | 0.34 | 1.82 | 0.99 |
| Agranulocyte Adhesion and Diapedesis | 3.83 | 2.35 | 0.42 | 0.23 | 1.05 | 0.50 |
| Regulation of Actin-based Motility by Rho | 3.78 | 2.34 | 0.32 | 0.22 | 0.74 | 0.33 |
| Remodeling of Epithelial Adherens Junctions | 3.72 | 2.31 | 0.43 | 0.23 | 2.27 | 1.28 |
| Mitochondrial Dysfunction | 3.55 | 2.20 | 0.94 | 0.38 | 0.00 | 0.00 |
| Glycolysis I | 3.56 | 2.20 | | | 0.90 | 0.42 |
| Protein Kinase A Signaling | 3.34 | 2.02 | 0.29 | 0.20 | 0.00 | 0.00 |
| ILK Signaling | 3.21 | 1.92 | 0.43 | 0.23 | 2.71 | 1.60 |
| G Beta Gamma Signaling | 3.09 | 1.86 | 2.00 | 0.78 | 0.79 | 0.35 |
| Leukocyte Extravasation Signaling | 3.10 | 1.86 | 1.33 | 0.55 | 3.58 | 2.23 |
| LXR/RXR Activation | 3.11 | 1.86 | | | 7.56 | 5.83 |
| Coagulation System | 3.06 | 1.85 | | | 16.70 | 14.10 |
| Glutaryl-CoA Degradation | 3.04 | 1.84 | 0.98 | 0.40 | 0.50 | 0.21 |
| FXR/RXR Activation | 3.01 | 1.84 | | | 5.03 | 3.44 |
| Sirtuin Signaling Pathway | 2.99 | 1.83 | 0.00 | 0.00 | 0.60 | 0.26 |
| Germ Cell-Sertoli Cell Junction Signaling | 2.88 | 1.74 | 0.94 | 0.38 | 1.68 | 0.88 |
| UDP-N-acetyl-D-glucosamine Biosynthesis II | 2.66 | 1.54 | 1.39 | 0.58 | 0.87 | 0.40 |
| Ethanol Degradation IV | 2.56 | 1.49 | 0.83 | 0.37 | 3.76 | 2.34 |
| Tryptophan Degradation III (Eukaryotic) | 2.56 | 1.49 | 0.83 | 0.37 | 0.37 | 0.14 |
| Epithelial Adherens Junction Signaling | 2.55 | 1.49 | 0.55 | 0.26 | 1.16 | 0.56 |
| Clathrin-mediated Endocytosis Signaling | 2.55 | 1.49 | 0.00 | 0.00 | 3.67 | 2.28 |
| Cardiac Hypertrophy Signaling | 2.51 | 1.48 | 1.60 | 0.65 | 0.00 | 0.00 |

| | | | | | | |
|---|---|---|---|---|---|---|
| Glycogen Biosynthesis II (from UDP-D-Glucose) | 2.52 | 1.48 | | | | |
| Gap Junction Signaling | 2.48 | 1.47 | 0.40 | 0.23 | 1.00 | 0.47 |
| Neuroprotective Role of THOP1 in Alzheimer's Disease | 2.48 | 1.47 | 0.26 | 0.19 | 2.18 | 1.21 |
| IL-8 Signaling | 2.46 | 1.46 | 1.90 | 0.77 | 2.08 | 1.17 |
| Tec Kinase Signaling | 2.36 | 1.38 | 0.98 | 0.40 | 0.46 | 0.18 |
| CXCR4 Signaling | 2.32 | 1.37 | 1.55 | 0.65 | 0.44 | 0.18 |
| Phagosome Formation | 2.34 | 1.37 | 1.26 | 0.52 | 2.01 | 1.12 |
| Phospholipase C Signaling | 2.32 | 1.37 | 1.00 | 0.40 | 1.07 | 0.50 |
| Integrin Signaling | 2.29 | 1.36 | 1.80 | 0.77 | 1.51 | 0.76 |
| Leucine Degradation I | 2.29 | 1.36 | | | | |
| IL-1 Signaling | 2.27 | 1.35 | 0.89 | 0.37 | 0.00 | 0.00 |
| Cellular Effects of Sildenafil (Viagra) | 2.24 | 1.33 | | | 0.70 | 0.31 |
| α-Adrenergic Signaling | 2.17 | 1.27 | 2.36 | 0.94 | 0.00 | 0.00 |
| Ephrin Receptor Signaling | 2.15 | 1.27 | 1.45 | 0.61 | 0.87 | 0.40 |
| Fatty Acid β-oxidation I | 2.15 | 1.27 | 0.70 | 0.34 | 0.76 | 0.34 |
| Axonal Guidance Signaling | 2.13 | 1.26 | 0.62 | 0.29 | 1.93 | 1.05 |
| Estrogen Receptor Signaling | 2.11 | 1.24 | 0.72 | 0.34 | 1.31 | 0.64 |
| Virus Entry via Endocytic Pathways | 1.98 | 1.14 | 2.19 | 0.88 | 0.96 | 0.44 |
| Relaxin Signaling | 1.96 | 1.14 | 0.56 | 0.26 | 0.55 | 0.24 |
| Complement System | 1.97 | 1.14 | | | 10.50 | 8.51 |
| UDP-N-acetyl-D-galactosamine Biosynthesis II | 1.97 | 1.14 | | | 0.57 | 0.25 |
| Phagosome Maturation | 1.95 | 1.13 | 0.56 | 0.26 | 1.17 | 0.56 |
| Oxidative Phosphorylation | 1.94 | 1.13 | | | 0.00 | 0.00 |
| tRNA Charging | 1.91 | 1.12 | 2.62 | 1.05 | 5.52 | 3.85 |
| Acetyl-CoA Biosynthesis III (from Citrate) | 1.91 | 1.12 | | | | |
| Lanosterol Biosynthesis | 1.91 | 1.12 | | | | |
| Thrombin Signaling | 1.82 | 1.05 | 1.25 | 0.52 | 0.91 | 0.42 |
| Intrinsic Prothrombin Activation Pathway | 1.83 | 1.05 | 0.60 | 0.29 | 11.10 | 8.92 |
| Agrin Interactions at Neuromuscular Junction | 1.78 | 1.04 | 2.67 | 1.05 | 1.96 | 1.07 |
| GP6 Signaling Pathway | 1.79 | 1.04 | 2.03 | 0.80 | 7.66 | 5.88 |
| Hepatic Fibrosis Signaling Pathway | 1.78 | 1.04 | 0.33 | 0.22 | 0.00 | 0.00 |
| Extrinsic Prothrombin Activation Pathway | 1.79 | 1.04 | | | 12.40 | 10.10 |
| Parkinson's Signaling | 1.79 | 1.04 | | | 0.50 | 0.21 |
| AMPK Signaling | 1.76 | 1.03 | 0.00 | 0.00 | 0.00 | 0.00 |
| Apelin Adipocyte Signaling Pathway | 1.73 | 1.02 | 0.37 | 0.23 | 1.35 | 0.64 |
| RhoA Signaling | 1.74 | 1.02 | | | 3.78 | 2.34 |
| Histamine Degradation | 1.74 | 1.02 | | | 3.24 | 2.03 |
| Gαi Signaling | 1.71 | 1.01 | 1.96 | 0.77 | 0.00 | 0.00 |
| PFKFB4 Signaling Pathway | 1.72 | 1.01 | | | | |
| Atherosclerosis Signaling | 1.70 | 1.00 | 0.24 | 0.18 | 1.52 | 0.76 |
| Molecular Mechanisms of Cancer | 1.62 | 0.98 | 0.88 | 0.37 | 0.00 | 0.00 |
| Primary Immunodeficiency Signaling | 1.63 | 0.98 | 0.54 | 0.26 | 1.52 | 0.76 |
| Granulocyte Adhesion and Diapedesis | 1.61 | 0.98 | 0.46 | 0.23 | 0.87 | 0.40 |
| Oxidative Ethanol Degradation III | 1.65 | 0.98 | | | 3.04 | 1.87 |
| Apelin Muscle Signaling Pathway | 1.65 | 0.98 | | | 0.44 | 0.18 |

| | | | | | | |
|---|---|---|---|---|---|---|
| Guanine and Guanosine Salvage I | 1.61 | 0.98 | | | 3.25 | 2.03 |
| Choline Degradation I | 1.61 | 0.98 | | | 1.33 | 0.64 |
| Sulfate Activation for Sulfonation | 1.61 | 0.98 | | | 1.33 | 0.64 |
| Fatty Acid Biosynthesis Initiation II | 1.61 | 0.98 | | | | |
| Palmitate Biosynthesis I (Animals) | 1.61 | 0.98 | | | | |
| Fatty Acid α-oxidation | 1.60 | 0.97 | | | 2.95 | 1.79 |
| Putrescine Degradation III | 1.56 | 0.94 | 0.87 | 0.37 | 2.87 | 1.72 |
| Androgen Signaling | 1.57 | 0.94 | 0.62 | 0.29 | 0.40 | 0.15 |
| Huntington's Disease Signaling | 1.55 | 0.94 | 1.09 | 0.45 | 2.79 | 1.67 |
| Hepatic Fibrosis / Hepatic Stellate Cell Activation | 1.55 | 0.94 | 0.86 | 0.37 | 1.87 | 1.02 |
| Production of Nitric Oxide and Reactive Oxygen Species in Macrophages | 1.53 | 0.92 | 0.00 | 0.00 | 1.10 | 0.52 |
| Cardiac β-adrenergic Signaling | 1.51 | 0.91 | 0.60 | 0.29 | 0.00 | 0.00 |
| Aryl Hydrocarbon Receptor Signaling | 1.49 | 0.90 | 1.12 | 0.46 | 0.91 | 0.42 |
| Endocannabinoid Cancer Inhibition Pathway | 1.49 | 0.90 | 0.20 | 0.17 | 0.00 | 0.00 |
| Hypusine Biosynthesis | 1.44 | 0.88 | 1.69 | 0.71 | | |
| S-adenosyl-L-methionine Biosynthesis | 1.44 | 0.88 | 1.69 | 0.71 | | |
| Glutathione Redox Reactions I | 1.46 | 0.88 | 0.82 | 0.37 | 1.74 | 0.92 |
| Inosine-5'-phosphate Biosynthesis II | 1.44 | 0.88 | | | 1.16 | 0.56 |
| NADH Repair | 1.44 | 0.88 | | | 1.16 | 0.56 |
| Diphthamide Biosynthesis | 1.44 | 0.88 | | | | |
| Oxidized GTP and dGTP Detoxification | 1.44 | 0.88 | | | | |
| Thyroid Hormone Biosynthesis | 1.44 | 0.88 | | | | |
| Semaphorin Signaling in Neurons | 1.42 | 0.87 | 1.19 | 0.50 | 1.87 | 1.02 |
| Tryptophan Degradation X (Mammalian, via Tryptamine) | 1.42 | 0.87 | 0.80 | 0.37 | 2.58 | 1.52 |
| Gluconeogenesis I | 1.39 | 0.85 | | | 0.33 | 0.12 |
| Calcium Signaling | 1.37 | 0.83 | | | 0.00 | 0.00 |
| Gαs Signaling | 1.36 | 0.83 | 0.78 | 0.36 | 0.00 | 0.00 |
| CDK5 Signaling | 1.35 | 0.82 | 2.18 | 0.88 | 0.00 | 0.00 |
| PXR/RXR Activation | 1.33 | 0.81 | | | | |
| Osteoarthritis Pathway | 1.33 | 0.80 | 0.37 | 0.23 | 0.89 | 0.41 |
| Pentose Phosphate Pathway (Oxidative Branch) | 1.32 | 0.80 | | | 1.04 | 0.50 |
| **PVH Whole Lung vs. Sham Lung: Ingenuity Pathways with Uncorrected P < 0.05 (-$\log_{10}$ P > 1.3)** | | | | | | |
| **Ingenuity Canonical Pathways** | **-Log10 (P-value)** | **-Log10 (FDR P-value)** | **-Log10 (P-value)** | **-Log10 (FDR P-value)** | **-Log10 (P-value)** | **-Log10 (FDR P-value)** |
| Coagulation System | 16.70 | 14.10 | | | 3.06 | 1.85 |
| Extrinsic Prothrombin Activation Pathway | 12.40 | 10.10 | | | 1.79 | 1.04 |
| Intrinsic Prothrombin Activation Pathway | 11.10 | 8.92 | 0.60 | 0.29 | 1.83 | 1.05 |
| Complement System | 10.50 | 8.51 | | | 1.97 | 1.14 |
| EIF2 Signaling | 9.83 | 7.91 | 48.10 | 45.60 | 7.68 | 5.41 |
| Acute Phase Response Signaling | 8.75 | 6.91 | 0.00 | 0.00 | 4.84 | 3.16 |
| GP6 Signaling Pathway | 7.66 | 5.88 | 2.03 | 0.80 | 1.79 | 1.04 |
| LXR/RXR Activation | 7.56 | 5.83 | | | 3.11 | 1.86 |
| tRNA Charging | 5.52 | 3.85 | 2.62 | 1.05 | 1.91 | 1.12 |
| Role of PKR in Interferon Induction and Antiviral | 5.39 | 3.76 | 0.72 | 0.34 | 0.37 | 0.22 |

| | | | | | | |
|---|---|---|---|---|---|---|
| Response | | | | | | |
| FXR/RXR Activation | 5.03 | 3.44 | | | 3.01 | 1.84 |
| Unfolded protein response | 4.31 | 2.77 | 2.17 | 0.88 | 0.30 | 0.18 |
| Regulation of eIF4 and p70S6K Signaling | 4.02 | 2.51 | 13.80 | 11.70 | 5.38 | 3.43 |
| Sertoli Cell-Sertoli Cell Junction Signaling | 3.86 | 2.38 | 0.44 | 0.23 | 5.46 | 3.43 |
| RhoA Signaling | 3.78 | 2.34 | | | 1.74 | 1.02 |
| Ethanol Degradation IV | 3.76 | 2.34 | 0.83 | 0.37 | 2.56 | 1.49 |
| Clathrin-mediated Endocytosis Signaling | 3.67 | 2.28 | 0.00 | 0.00 | 2.55 | 1.49 |
| Signaling by Rho Family GTPases | 3.61 | 2.24 | 0.63 | 0.29 | 4.94 | 3.16 |
| Leukocyte Extravasation Signaling | 3.58 | 2.23 | 1.33 | 0.55 | 3.10 | 1.86 |
| Systemic Lupus Erythematosus Signaling | 3.41 | 2.10 | 0.00 | 0.00 | 1.19 | 0.73 |
| Aldosterone Signaling in Epithelial Cells | 3.41 | 2.10 | 1.02 | 0.42 | 0.24 | 0.14 |
| Role of Tissue Factor in Cancer | 3.33 | 2.05 | 0.72 | 0.34 | 0.75 | 0.47 |
| BAG2 Signaling Pathway | 3.31 | 2.05 | 0.59 | 0.29 | | |
| mTOR Signaling | 3.30 | 2.05 | 11.90 | 9.87 | 4.90 | 3.16 |
| Guanine and Guanosine Salvage I | 3.25 | 2.03 | | | 1.61 | 0.98 |
| Histamine Degradation | 3.24 | 2.03 | | | 1.74 | 1.02 |
| Dopamine Degradation | 3.20 | 2.00 | 0.73 | 0.34 | 1.28 | 0.77 |
| Oxidative Ethanol Degradation III | 3.04 | 1.87 | | | 1.65 | 0.98 |
| Fatty Acid α-oxidation | 2.95 | 1.79 | | | 1.60 | 0.97 |
| Putrescine Degradation III | 2.87 | 1.72 | 0.87 | 0.37 | 1.56 | 0.94 |
| eNOS Signaling | 2.84 | 1.70 | 0.53 | 0.25 | 0.87 | 0.54 |
| Huntington's Disease Signaling | 2.79 | 1.67 | 1.09 | 0.45 | 1.55 | 0.94 |
| ILK Signaling | 2.71 | 1.60 | 0.43 | 0.23 | 3.21 | 1.92 |
| Cleavage and Polyadenylation of Pre-mRNA | 2.61 | 1.53 | | | | |
| Guanosine Nucleotides Degradation III | 2.61 | 1.53 | | | | |
| Tryptophan Degradation X (Mammalian, via Tryptamine) | 2.58 | 1.52 | 0.80 | 0.37 | 1.42 | 0.87 |
| D-myo-inositol (1,4,5)-Trisphosphate Biosynthesis | 2.58 | 1.52 | | | 0.57 | 0.36 |
| Urate Biosynthesis/Inosine 5'-phosphate Degradation | 2.50 | 1.46 | | | | |
| Heme Degradation | 2.49 | 1.45 | | | | |
| Mevalonate Pathway I | 2.40 | 1.38 | | | | |
| Adenosine Nucleotides Degradation II | 2.32 | 1.30 | | | | |
| Remodeling of Epithelial Adherens Junctions | 2.27 | 1.28 | 0.43 | 0.23 | 3.72 | 2.31 |
| CMP-N-acetylneuraminate Biosynthesis I (Eukaryotes) | 2.27 | 1.28 | | | | |
| Protein Ubiquitination Pathway | 2.24 | 1.26 | 3.21 | 1.39 | 0.91 | 0.55 |
| Neuroprotective Role of THOP1 in Alzheimer's Disease | 2.18 | 1.21 | 0.26 | 0.19 | 2.48 | 1.47 |
| Ethanol Degradation II | 2.18 | 1.21 | | | 1.23 | 0.75 |
| Ephrin B Signaling | 2.15 | 1.19 | 1.87 | 0.77 | 4.52 | 2.93 |
| Natural Killer Cell Signaling | 2.13 | 1.18 | 1.33 | 0.55 | 0.65 | 0.40 |
| Adenine and Adenosine Salvage III | 2.10 | 1.17 | | | 1.14 | 0.71 |
| IL-8 Signaling | 2.08 | 1.17 | 1.90 | 0.77 | 2.46 | 1.46 |
| Purine Nucleotides Degradation II (Aerobic) | 2.08 | 1.17 | | | | |
| Superpathway of Geranylgeranyldiphosphate Biosynthesis I (via Mevalonate) | 2.08 | 1.17 | | | | |

| | | | | | | |
|---|---|---|---|---|---|---|
| Noradrenaline and Adrenaline Degradation | 2.04 | 1.14 | 0.67 | 0.32 | 1.16 | 0.72 |
| Phagosome Formation | 2.01 | 1.12 | 1.26 | 0.52 | 2.34 | 1.37 |
| Agrin Interactions at Neuromuscular Junction | 1.96 | 1.07 | 2.67 | 1.05 | 1.78 | 1.04 |
| Axonal Guidance Signaling | 1.93 | 1.05 | 0.62 | 0.29 | 2.13 | 1.26 |
| Endoplasmic Reticulum Stress Pathway | 1.90 | 1.03 | 0.87 | 0.37 | | |
| Inhibition of Matrix Metalloproteases | 1.88 | 1.02 | 2.62 | 1.05 | | |
| Hepatic Fibrosis / Hepatic Stellate Cell Activation | 1.87 | 1.02 | 0.86 | 0.37 | 1.55 | 0.94 |
| Semaphorin Signaling in Neurons | 1.87 | 1.02 | 1.19 | 0.50 | 1.42 | 0.87 |
| Airway Pathology in Chronic Obstructive Pulmonary Disease | 1.85 | 1.01 | 1.27 | 0.52 | | |
| Actin Cytoskeleton Signaling | 1.82 | 0.99 | 0.72 | 0.34 | 4.05 | 2.54 |
| Glutathione Redox Reactions I | 1.74 | 0.92 | 0.82 | 0.37 | 1.46 | 0.88 |
| Tumoricidal Function of Hepatic Natural Killer Cells | 1.74 | 0.92 | 0.82 | 0.37 | | |
| Sucrose Degradation V (Mammalian) | 1.74 | 0.92 | | | | |
| Tight Junction Signaling | 1.73 | 0.92 | 0.00 | 0.00 | 4.33 | 2.78 |
| Germ Cell-Sertoli Cell Junction Signaling | 1.68 | 0.88 | 0.94 | 0.38 | 2.88 | 1.74 |
| Serotonin Degradation | 1.68 | 0.88 | 0.43 | 0.23 | 0.70 | 0.43 |
| Melatonin Degradation III | 1.63 | 0.85 | | | | |
| Xanthine and Xanthosine Salvage | 1.63 | 0.85 | | | | |
| Purine Nucleotides De Novo Biosynthesis II | 1.57 | 0.80 | | | 0.89 | 0.55 |
| Atherosclerosis Signaling | 1.52 | 0.76 | 0.24 | 0.18 | 1.70 | 1.00 |
| Primary Immunodeficiency Signaling | 1.52 | 0.76 | 0.54 | 0.26 | 1.63 | 0.98 |
| Superpathway of Cholesterol Biosynthesis | 1.52 | 0.76 | | | 0.52 | 0.32 |
| Integrin Signaling | 1.51 | 0.76 | 1.80 | 0.77 | 2.29 | 1.36 |
| G Protein Signaling Mediated by Tubby | 1.44 | 0.70 | 1.72 | 0.71 | 1.25 | 0.76 |
| Role of IL-17A in Psoriasis | 1.44 | 0.70 | | | 0.83 | 0.51 |
| Chemokine Signaling | 1.39 | 0.65 | 0.37 | 0.23 | 0.59 | 0.37 |
| Glycogen Degradation III | 1.38 | 0.65 | 2.39 | 0.94 | 0.80 | 0.49 |
| Apelin Adipocyte Signaling Pathway | 1.35 | 0.64 | 0.37 | 0.23 | 1.73 | 1.02 |
| Inhibition of Angiogenesis by TSP1 | 1.34 | 0.64 | 0.68 | 0.32 | 0.46 | 0.29 |
| Choline Degradation I | 1.33 | 0.64 | | | 1.61 | 0.98 |
| Sulfate Activation for Sulfonation | 1.33 | 0.64 | | | 1.61 | 0.98 |
| Superpathway of Inositol Phosphate Compounds | 1.33 | 0.64 | | | 0.64 | 0.40 |
| 4-hydroxyproline Degradation I | 1.33 | 0.64 | | | | |
| Adenine and Adenosine Salvage I | 1.33 | 0.64 | | | | |
| UDP-D-xylose and UDP-D-glucuronate Biosynthesis | 1.33 | 0.64 | | | | |
| iCOS-iCOSL Signaling in T Helper Cells | 1.32 | 0.64 | | | 0.00 | 0.00 |
| Isoleucine Degradation I | 1.32 | 0.64 | | | | |
| Estrogen Receptor Signaling | 1.31 | 0.64 | 0.72 | 0.34 | 2.11 | 1.24 |

**Supplemental Table 3. Ingenuity® Predicted Upstream Regulators with FDR p Value <0.01 (-log p > 2.0) AND I Z-score I ≥ 2.0**

| Upstream Regulator Gene Symbol | Molecule Type | Predicted Activation State | Z-score | -Log10 (FDR P-value) | Predicted Activation State | Z-score | -Log10 (FDR P-value) | Predicted Activation State | Z-score | -Log10 (FDR P-value) |
|---|---|---|---|---|---|---|---|---|---|---|
| PVH Artery vs. Sham Artery: Ingenuity® Predicted Upstream Regulators with FDR p Value <0.01 (-log p > 2.0) AND I Z-score I ≥ 2.0 | | | | | | | | | | |
| | | Pulmonary Artery | | | Pulmonary Vein | | | Whole Lung | | |
| MLXIPL | transcription regulator | Activated | 6.2 | 49.08 | Activated | 2.6 | 4.63 | Activated | 4 | 6.01 |
| MYCN | transcription regulator | Activated | 5.3 | 38.34 | Activated | 2.1 | 13.58 | | 1.14 | 13.86 |
| MYC | transcription regulator | Activated | 6.1 | 31.66 | Activated | 2.9 | 16.16 | Activated | 3.315 | 12.77 |
| TCR | complex | Activated | 2.7 | 9.37 | | 1.4 | 2.31 | Activated | 2.334 | 5.27 |
| SYVN1 | transporter | Activated | 2.1 | 3.36 | | 1.9 | 3.40 | | 0.535 | 3.41 |
| EGFR | kinase | Activated | 2.0 | 2.74 | Activated | 2.2 | 2.77 | | 0.994 | 2.63 |
| CD3 | complex | Activated | 2.4 | 2.25 | | 1.2 | 4.43 | | 1.81 | 4.49 |
| PVH Vein vs. Sham Vein: Ingenuity® Predicted Upstream Regulators with FDR p Value <0.01 (-log p > 2.0) AND I Z-score I ≥ 2.0 | | | | | | | | | | |
| | | Pulmonary Vein | | | Pulmonary Artery | | | Whole Lung | | |
| TGFB1 | growth factor | Activated | 2.7 | 19.11 | | 1.614 | 3.91 | | 1.973 | 18.52 |
| MYC | transcription regulator | Activated | 2.9 | 16.16 | Activated | 6.087 | 31.66 | Activated | 3.315 | 12.77 |
| MYCN | transcription regulator | Activated | 2.1 | 13.58 | Activated | 5.283 | 38.34 | | 1.14 | 13.86 |
| ETV5 | transcription regulator | Activated | 2.7 | 8.33 | | 1 | 2.58 | | | |
| ERBB2 | kinase | Activated | 2.1 | 5.86 | | -0.577 | 0.90 | | -0.592 | 3.78 |
| KDM8 | enzyme | Activated | 2.8 | 4.69 | | | | | | |
| MLXIPL | transcription regulator | Activated | 2.6 | 4.63 | Activated | 6.164 | 49.08 | Activated | 4 | 6.01 |
| Insulin | group | Activated | 2.5 | 4.24 | Activated | 2.351 | 1.09 | | 1.655 | 5.31 |
| XBP1 | transcription regulator | Activated | 2.6 | 4.13 | | 1.761 | 1.63 | Activated | 2.531 | 6.16 |
| PTK2 | kinase | Activated | 2.2 | 3.56 | | | | | 0.216 | 2.10 |
| STK11 | kinase | Activated | 2.4 | 3.25 | | -0.447 | 0.91 | | 0.832 | 0.99 |

| | | | | | | | | | | |
|---|---|---|---|---|---|---|---|---|---|---|
| Esrra | transcription regulator | Activated | 2.1 | 3.22 | | | | Activated | 2.219 | 0.63 |
| EGFR | kinase | Activated | 2.2 | 2.77 | Activated | 2.001 | 2.74 | | 0.994 | 2.63 |
| MRTFA | transcription regulator | Activated | 2.4 | 2.67 | | | | Activated | 2.098 | 7.54 |
| IPMK | kinase | Activated | 2.0 | 2.57 | | | | | | |
| FOS | transcription regulator | Activated | 2.9 | 2.31 | | 0.378 | 2.69 | | 0.629 | 1.65 |
| ERN1 | kinase | Activated | 2.4 | 2.23 | | | 1.34 | Activated | 2.153 | 3.94 |
| LEP | growth factor | Inhibited | -2.8 | 4.06 | | | | | -0.956 | 1.18 |
| IFNG | cytokine | Inhibited | -2.4 | 3.94 | | -1.744 | 1.88 | Inhibited | -2.271 | 6.05 |
| ACOX1 | enzyme | Inhibited | -2.3 | 2.72 | | | | | -1.098 | 1.21 |
| miR-1-3p (+others GGAAUGU) | mature microRNA | Inhibited | -2.6 | 2.52 | | -1.964 | 0.77 | | | |
| NR4A1 | ligand-dependent nuclear receptor | Inhibited | -2.1 | 2.42 | | | | | | |
| **PVH Whole Lung vs. Sham Whole Lung: Ingenuity® Predicted Upstream Regulators with FDR p Value <0.01 (-log p > 2.0) AND I Z-score I ≥ 2.0** | | | | | | | | | | |
| | | **Whole Lung** | | | **Pulmonary Artery** | | | **Pulmonary Vein** | | |
| MYC | transcription regulator | Activated | 3.3 | 12.77 | Activated | 6.087 | 31.66 | Activated | 2.913 | 16.16 |
| MRTFB | transcription regulator | Activated | 2.6 | 8.84 | | | | | 1.769 | 2.95 |
| MRTFA | transcription regulator | Activated | 2.1 | 7.54 | | | | Activated | 2.368 | 2.67 |
| XBP1 | transcription regulator | Activated | 2.5 | 6.16 | | 1.761 | 1.63 | Activated | 2.611 | 4.13 |
| MLXIPL | transcription regulator | Activated | 4.0 | 6.01 | Activated | 6.164 | 49.08 | Activated | 2.551 | 4.63 |
| TCR | complex | Activated | 2.3 | 5.27 | Activated | 2.735 | 9.37 | | 1.408 | 2.31 |
| SRF | transcription regulator | Activated | 3.1 | 4.91 | | | 0.80 | | 0.953 | 2.57 |
| Immunoglobulin | complex | Activated | 2.4 | 4.36 | | | | | 1.571 | 2.14 |

| | | | | | | | | | | |
|---|---|---|---|---|---|---|---|---|---|---|
| ERN1 | kinase | Activated | 2.2 | 3.94 | | | 1.34 | Activated | 2.449 | 2.23 |
| PDGF (family) | group | Activated | 2.0 | 2.93 | | | | | | |
| NOSTRIN | transcription regulator | Activated | 2.4 | 2.59 | | -0.064 | 2.57 | | 0.618 | 3.94 |
| EPO | cytokine | Activated | 2.0 | 2.55 | | | 1.65 | | -0.131 | 3.17 |
| CD38 | enzyme | Activated | 2.2 | 2.04 | | | | | | |
| HNF4A | transcription regulator | Inhibited | -2.8 | 9.94 | | | | | -0.292 | 3.24 |
| TCL1A | transcription regulator | Inhibited | -2.2 | 8.82 | | | | | | 2.61 |
| IFNG | cytokine | Inhibited | -2.3 | 6.05 | | -1.744 | 1.88 | Inhibited | -2.413 | 3.94 |
| CEBPA | transcription regulator | Inhibited | -2.4 | 5.10 | | | | | -0.613 | 5.21 |
| miR-338-3p (+others CCAGCAU) | mature microRNA | Inhibited | -3.0 | 5.08 | | | 1.54 | | -1.342 | 2.86 |
| miR-29b-3p (+others AGCACCA) | mature microRNA | Inhibited | -2.9 | 4.75 | | -1.964 | 1.31 | | -0.686 | 2.00 |
| KITLG | growth factor | Inhibited | -2.6 | 4.57 | | | 3.86 | | -1.982 | 3.18 |
| IL6 | cytokine | Inhibited | -3.0 | 3.93 | | | | | -1.657 | 2.20 |
| IL17A | cytokine | Inhibited | -2.7 | 3.91 | | | | | -1.656 | 1.15 |
| miR-335-3p (+others UUUUCAU) | mature microRNA | Inhibited | -2.6 | 3.72 | | | 1.72 | | -1 | 2.24 |
| TGM2 | enzyme | Inhibited | -3.1 | 3.72 | | | | Inhibited | -2 | 0.60 |
| SMARCA4 | transcription regulator | Inhibited | -2.2 | 3.55 | | 0.378 | 0.80 | | 0.87 | 2.52 |
| CSF3 | cytokine | Inhibited | -2.3 | 3.53 | | | | | -0.664 | 2.23 |
| OSM | cytokine | Inhibited | -2.4 | 3.43 | | 0.632 | 0.91 | | -0.984 | 2.36 |
| Mmp | group | Inhibited | -2.2 | 3.43 | | | | | | 1.25 |
| PKD1 | ion channel | Inhibited | -2.1 | 3.40 | | | | | 0.508 | 2.36 |
| CSF2 | cytokine | Inhibited | -2.1 | 2.32 | | | | | | |
| SPI1 | transcription regulator | Inhibited | -2.8 | 2.25 | | | 0.97 | | -0.751 | 1.09 |

| | | | | | | | | | | |
|---|---|---|---|---|---|---|---|---|---|---|
| HIPK2 | kinase | Inhibited | -2.4 | 2.21 | | | | | -1.412 | 1.71 |

**Figure 1. Animal Study Design**

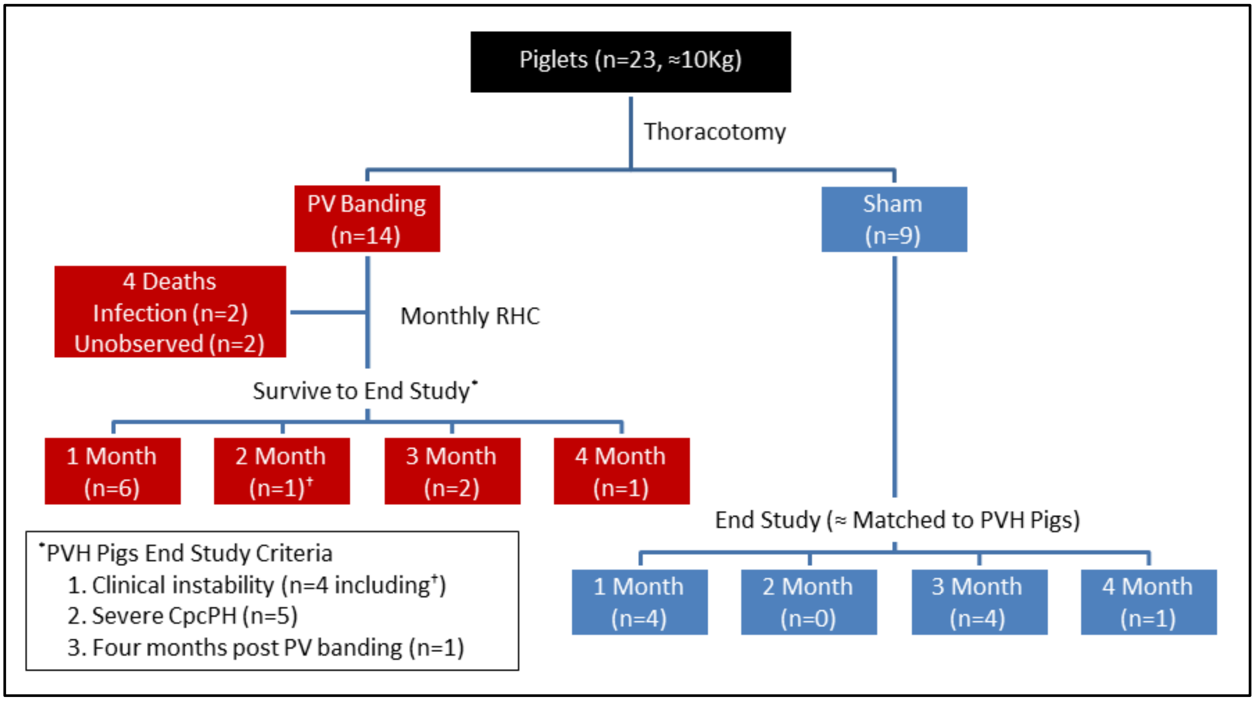


†Unobserved death 2 days after 2 month RHC; included in End Study Survivors but with hemodynamic data only. CpcPH indicated combined pre- and post-capillary pulmonary hypertension; PV, pulmonary vein; and RHC, right heart catheterization.

**Figure 2. Whole slide digital microscopic scanning of porcine lung specimens with annotation of pulmonary vessels with quantitative histomorphometry measurement.**

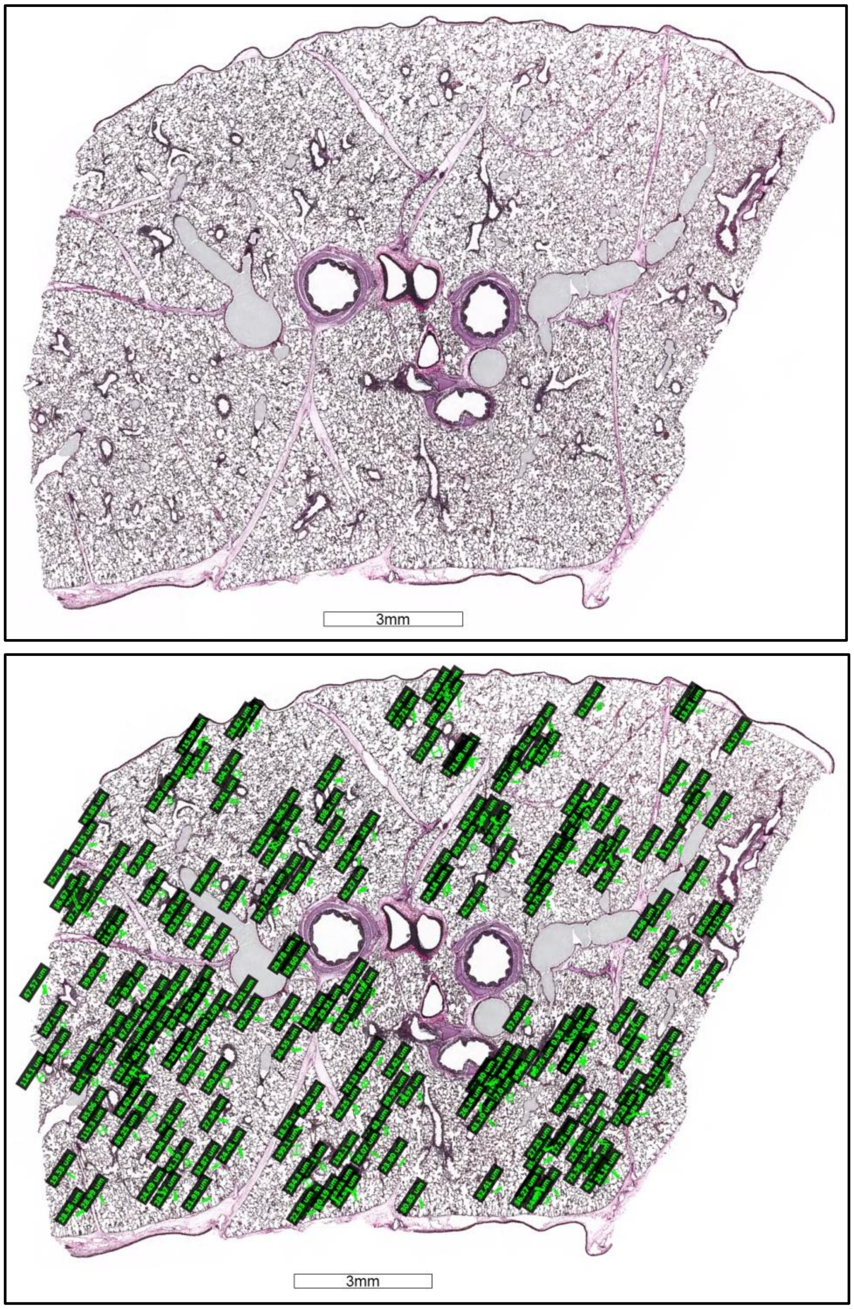


Upper: Whole slide digital microscopic scanning of lung specimen, displaying the Barium-agarose gel (light grey) in the lumens of pulmonary veins.

Lower: A representative example of whole slide digital scan of the entire lung specimen displaying the annotations of quantitative histomorphometry. Each vessel meeting the criteria for morphometric analysis was annotated and analyzed at higher power (20x–40x).

**Figure 3. Workflow of laser capture microdissection of porcine pulmonary arteries and veins using Zeiss PALM Microbream system.**

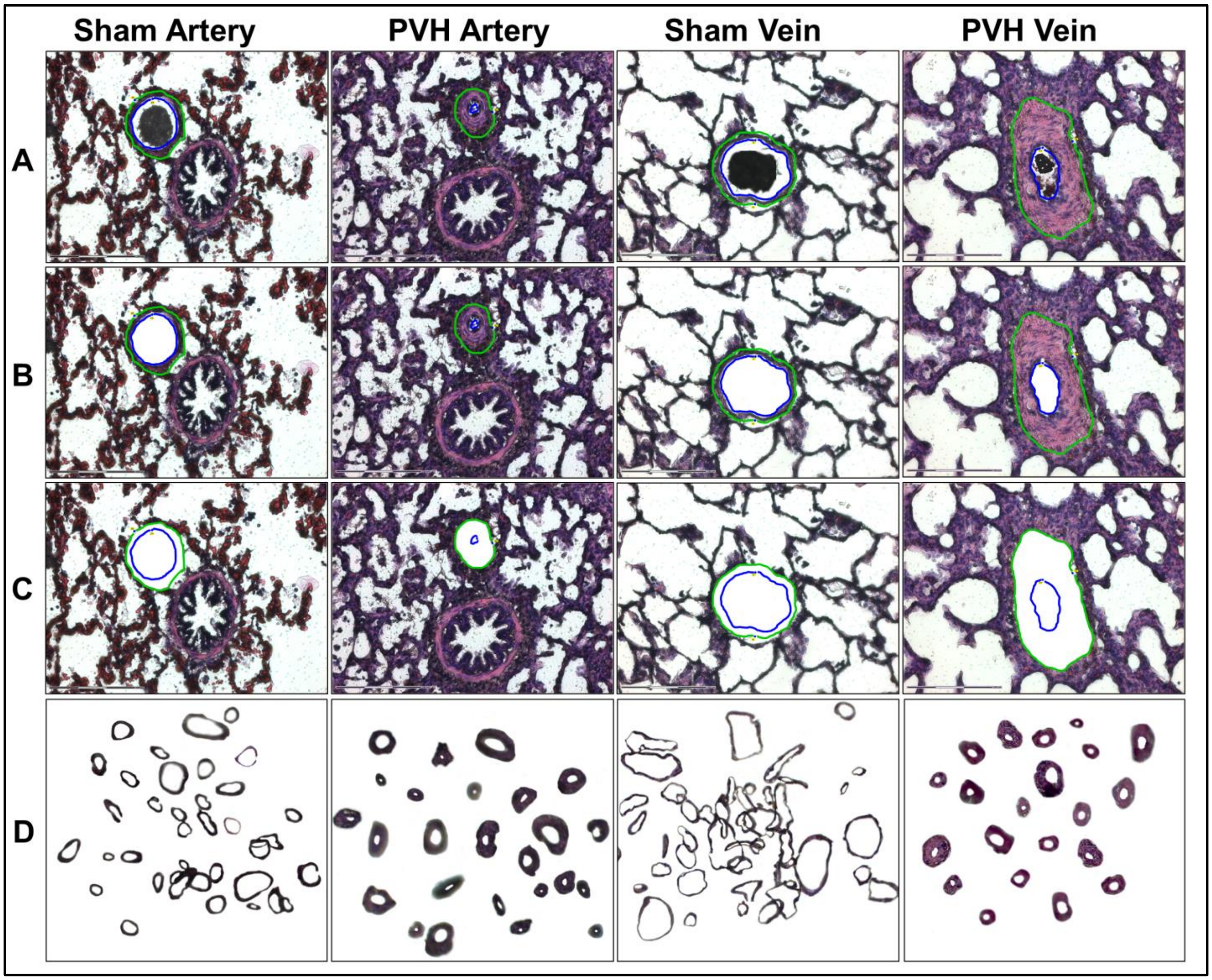


**(A)** H&E stained, thick (10 µm) formalin-fixed paraffin embedded tissues were mounted on the laser microdissection slides (ArcturusXT$^{TM}$) coated with polyethylene napthalate (PEN) membrane. To sterilize and to overcome the hydrophobic nature of PEN membrane, prior to mounting the tissue, slides were irradiated with UV light at 254 nm for 30 minutes. **(B)** When present, labelling material and/or blood cells were cleared off the lumen of the vascular profile (blue perimeter). **(C)** Each vascular profile was cut along the outer edge of the adventitia. For each vascular specimen, approximately 500,000 µm$^2$ area of vascular tissue was collected. Tissue area was calculated by subtracting the luminal area (blue perimeter) from the outer area (green perimeter) using Zeiss PALM RoboSoftware 4.6 (Bernried, Germany). **(D)** Tissue Capture Check: Micro-dissected vascular tissue was catapulted and collected in opaque adhesive caps of 0.5µL microfuge tubes (Zeiss), invertedly placed on the slide. Tubes were stored at -80 °C until mass spectrometry.

**Figure 4. Conscious Pulmonary Artery (PA) Pressure Assessment in Pigs**

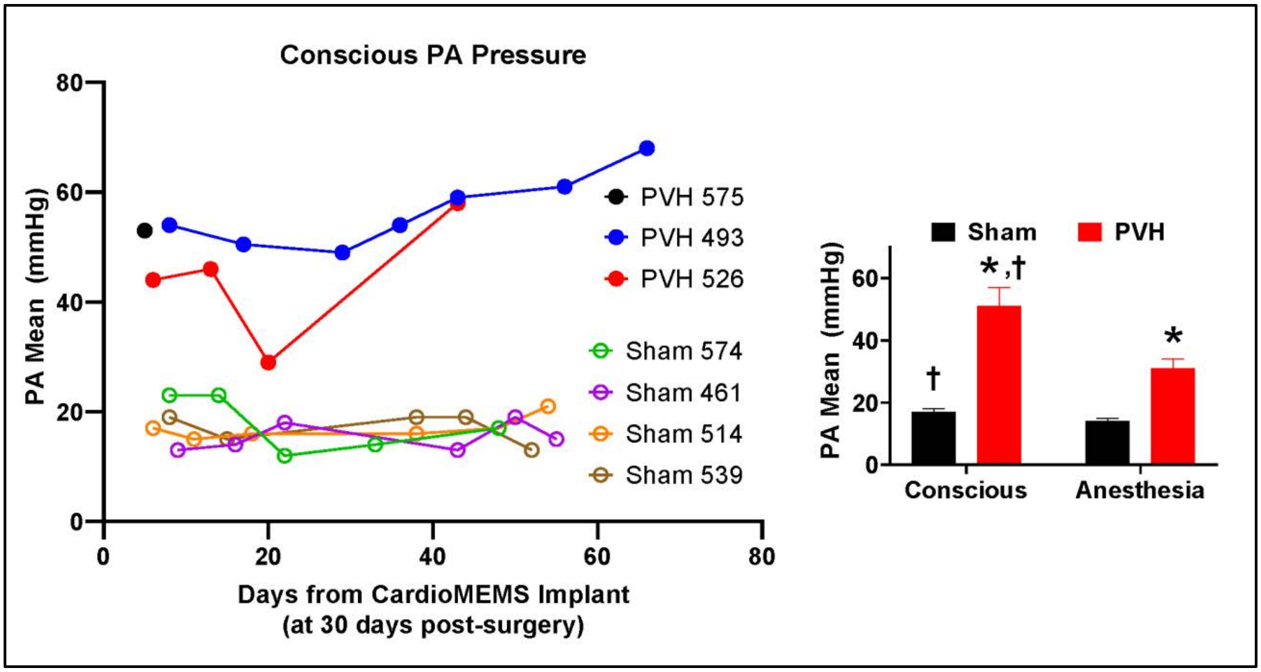


Left: Individual conscious measurements over time in PVH and Sham pigs.
Right: Average of averages of all measurements in PVH and Sham pigs at conscious and corresponding similar timed anesthetized measurements.
* P<0.001 PVH vs Sham; † p < 0.03 vs anesthetized measurements (Students t test without correction for multiple comparisons).

**Figure 5. Histological demonstration of Verhoeff-van Gieson (VVG) staining.**

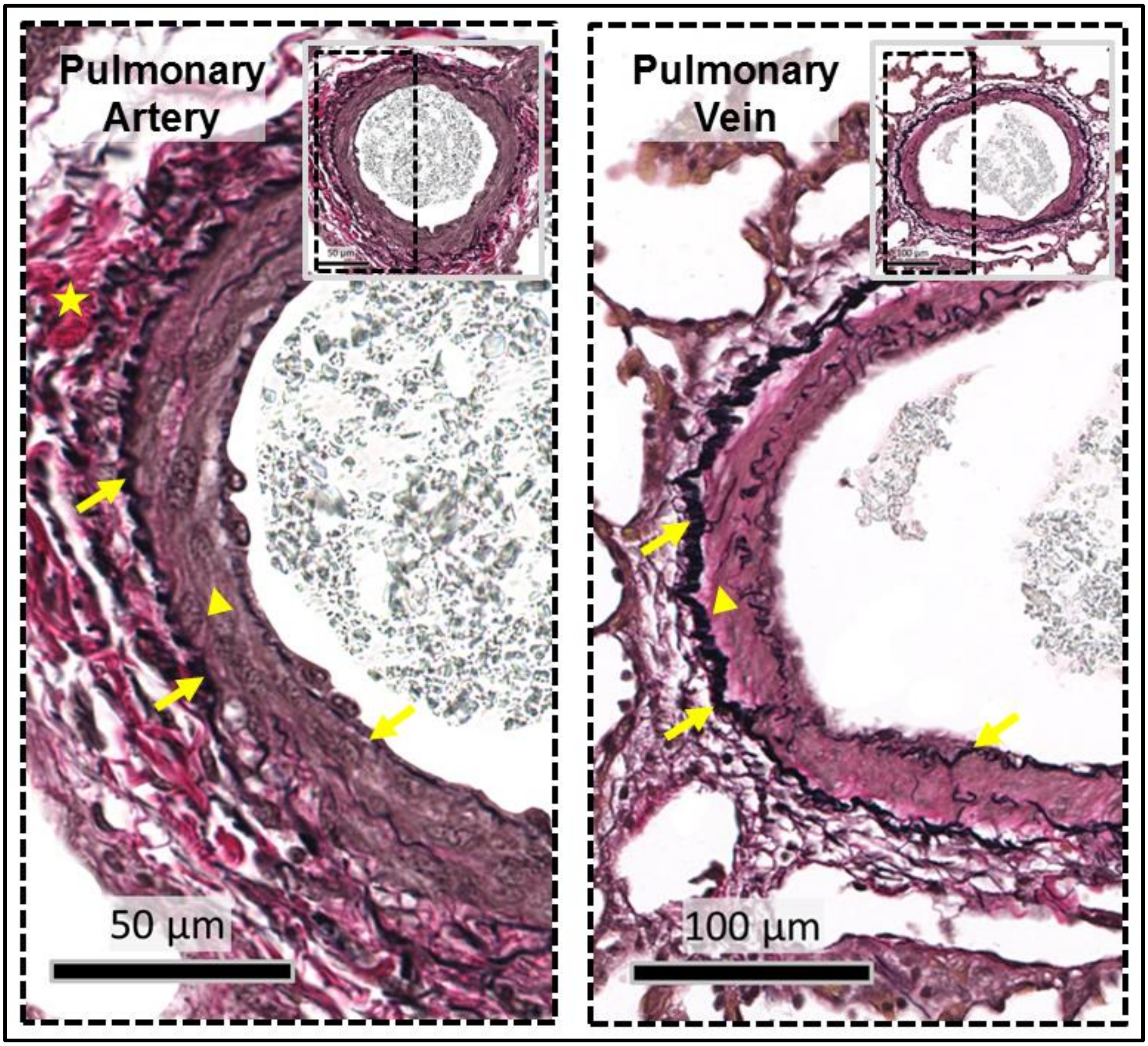


A representative photomicrograph displaying the Verhoeff-van Gieson (VVG) stained pulmonary artery and pulmonary vein. VVG stains collagen/fibrosis (star) red, smooth muscle cells (arrow head) brown and elastic tissue (arrows) black.
